# Unraveling the Web of Life: Incomplete lineage sorting and hybridization as primary mechanisms over polyploidization in the evolutionary dynamics of pear species

**DOI:** 10.1101/2024.07.29.605463

**Authors:** Ze-Tao Jin, Xiao-Hua Lin, Dai-Kun Ma, Richard G.J. Hodel, Chen Ren, Liang Zhao, Lei Duan, Chao Xu, Jun Wu, Bin-Bin Liu

## Abstract

In contrast to the traditional Tree of Life (ToL) paradigm, the Web of Life (WoL) model provides a more nuanced and precise depiction of organismal phylogeny, particularly considering the prevalent incongruence observed among gene/species trees. The lack of a generalized pipeline for teasing apart potential evolutionary mechanisms—such as Incomplete Lineage Sorting (ILS), hybridization, introgression, polyploidization, and Whole-Genome Duplication—poses significant challenges to the delineation of the WoL. The pear genus *Pyrus*, characterized by extensive hybridization events, serves as an excellent model for investigating the WoL. This study introduces a novel Step-by-Step Exclusion (SSE) approach to deciphering the complexities inherent in the WoL. Our findings indicate: 1) ILS, rather than polyploidization, is identified as the primary driver behind the origin of *Pyrus* from the arid regions of the Himalayas-Central Asia; 2) the two subgenera of *Pyrus* have independent evolutionary trajectories, facilitated by the geographical barriers that arose via the uplift of the Tibetan Plateau and increased aridity in Central Asia; 3) ILS and hybridization have facilitated the diversification of Oriental pears, while hybridization alone has driven the reticulate evolution of Occidental pears; 4) the establishment of the Silk Road during the Han Dynasty acted as a conduit for genetic exchange between Occidental and Oriental pears. The novel SSE approach provides a universally applicable framework for investigating evolutionary mechanisms defining the WoL paradigm.

## Introduction

Recent advances in phylogenomic research have increasingly indicated that a “Web of Life” (WoL) framework, as opposed to the traditional “Tree of Life” (ToL) model, offers a more comprehensive and accurate representation of organismal phylogeny (Swithers et al. 2009; Mao et al. 2017). This paradigm shift is substantiated by the growing body of evidence indicating widespread discordance between gene trees and species trees across diverse taxa, including animals and plants (Feng et al. 2022; Liu et al. 2022; Duan et al. 2023a; Gardner et al. 2023; Jin et al. 2023; Rivas-González et al. 2023; Xue et al. 2023). Additionally, bioinformatic tools have paved the way for more detailed phylogenomic discordance analyses, substantially improving our understanding of the intricate patterns inherent in the WoL (Liu et al. 2022; Xu et al. unpublished data). Moreover, a variety of factors, including both non- biological elements such as stochastic and systematic errors, as well as biological mechanisms including incomplete lineage sorting (ILS), horizontal gene transfer (HGT), hybridization, polyploidization, and introgression, have been identified as pivotal drivers of netlike phylogenies (Duan et al. 2023a; Steenwyk et al. 2023). This evolving perspective not only challenges the traditional bifurcating tree model but also can guide how we conceptualize and study the reticulate nature of evolution.

ILS frequently co-occurs with species’ rapid radiations, characterized by the stochastic retention of ancestral polymorphisms in certain genes across multiple divergent species (Meleshko et al. 2021). This phenomenon has been empirically observed in diverse groups, and primates (Rivas-González et al. 2023). Although many studies recognize the contribution of ILS to reticulate evolution, differentiating it from other evolutionary processes remains a complex challenge (Duan et al. 2023a; Nie et al. 2023; Yu et al. 2023). Allopolyploidization, often arising from one or multiple hybridization events, not only drives speciation, but also adaptation to extreme environmental conditions (Soltis PS and Soltis DE 2009; Lamichhaney et al. 2015). Advanced bioinformatic tools enable the concurrent analysis of hybridization and allopolyploidization (Thomas et al. 2017; Li et al. 2018; Yang et al. 2018; Morales-Briones et al. 2022). Case studies leveraging these tools have enhanced our understanding of specific Angiosperm lineages, exemplified by studies in the Vitaceae (Yu et al. 2023) and Gesneriaceae families (Yang et al. 2023). Accurate identification and verification of polyploidy events are necessary to reconstruct accurate evolutionary relationships among taxa, and to assess the impact of hybridization/polyploidization on the expansion of distribution ranges (Mu et al. 2023). Introgression, characterized by the repeated backcrossing of hybrid offspring with one parental species, results in nuclear genes in hybrids resembling those from one parent, a phenomenon frequently observed in sympatric species (Gardner et al. 2023).

Similarly, chloroplast capture events may occur, wherein hybrid offspring acquire the maternal chloroplast genome through continuous backcrosses with the paternal parent (Rieseberg and Soltis 1991; Okuyama et al. 2005; Liu et al. 2020; Duan et al. 2023b). This process has been documented in various plant lineages, such as the *Amelanchier- Malacomeles-Peraphyllum* clade within the Rosaceae family (Liu et al. 2020), the *Quercus* and Quercoideae groups within Fagales (Yang et al. 2021), and the *Taxus* genus within a Gymnosperm lineage (Qin et al. 2023). The widespread occurrence of gene introgression complicates phylogenetic analysis among closely related species by obscuring signals of ancient gene flow. Our awareness of the prevalence of reticulate evolution necessitates a shift from the traditional Tree of Life (ToL) evolutionary model to a network-based one (Zhang et al. 2021).

The genus *Pyrus*, a member of the Rosaceae family within the apple tribe (Maleae), stands as a paradigmatic model for exploring evolutionary mechanisms (Wu et al. 2018). This genus, globally recognized for its extensive array of cultivated varieties and significant commercial value, comprises perennial deciduous trees or shrubs primarily native to the temperate regions of the Eurasian continent (Fedorov 1954; Browicz 1993; Zamani et al. 2012). The evolutionary history of pear species is remarkably complex, predominantly driven by factors such as hybridization, introgression, and polyploidy. Furthermore, there is disruption of the self-incompatibility mechanisms in polyploids, with hybridization playing a critical and central role in the evolution of the *Pyrus* genus (Rubtsov 1944; Zielinski and Thompson 1967; Bell and Hough 1986). The classification of *Pyrus* species has long been contentious, with varying proposals from different researchers worldwide, suggesting species numbers ranging from 21 to 40 (Challice and Westwood 1973; Browicz 1993; Bell et al. 1996; Teng et al. 2004; Wu et al. 2013) or even as high as 50-80 species (Phipps et al. 1990; Gu and Spongberg 2003; Govaerts et al. 2021). This high species diversity arose due to reliance on traditional morphological characteristics for *Pyrus* species identification, compounded by hybridization leading to intermediate morphological phenotypes (Rubtsov 1944; Westwood and Challice 1978; Jang et al. 1992; Korotkova et al. 2018). The impact of a history of hybridization on the taxonomic classification and inferred evolutionary history of *Pyrus* species is evident, presenting challenges to our understanding of this group.

While evidence such as pollen structure, isoenzyme markers, and chemical data has been used to delineate *Pyrus* species (Challice and Westwood 1973; Westwood and Challice 1978; Zou et al. 1986; Jang et al. 1992), these characters did not provide sufficient information for robust phylogenetic reconstruction. DNA molecules, including RFLP (Kawata et al. 1995; Iketani et al. 1998), RAPD (Teng et al. 2001, 2002), and AFLP (Monte-Corve et al. 2000; Bao et al. 2008) markers, were used to characterize the genetic diversity of certain pear species.

However, due to the complex evolutionary history of *Pyrus*, these variable DNA molecular markers failed to reconstruct its evolutionary history accurately. Zheng et al. (2008) used nuclear gene ITS sequences to conduct a phylogenetic analysis of *Pyrus* species primarily from East Asia, revealing monophyletic species and indicating potential extensive ancient hybridization events. Building upon this, the researchers utilized non-coding sequences from two chloroplast regions and low-copy nuclear genes to reconstruct the phylogenetic relationships within *Pyrus*. The phylogenetic trees revealed significant cytonuclear discordance, underscoring the complexity of the evolutionary network that *Pyrus* has experienced (Zheng et al. 2014). Results from Wu et al. (2018), based on SNP data, also indicate the clustering of the *Pyrus* into two major groups: Asian pear and European pear.

The network-like diversification presents challenges in determining the spatiotemporal origins of the genus *Pyrus*. Paleontological evidence suggests that *Pyrus* likely emerged in the Paleocene, around 65-55 million years ago (Mya) or even earlier (Rubtsov 1944; Jang et al. 1992; Silva et al. 2014). Mountainous regions in western and southwestern China have been identified in prior studies as potential origins and domestication centers for *Pyrus*, subsequently spreading westward and eastward. Two additional diversification centers were hypothesized to be in the region from Asia Minor to the Middle East in the Caucasus Mountains, and Central Asia (Vavilov 1951; Zukovskij 1962; Vavilov 1992; Silva et al. 2014). These distribution patterns imply that *Pyrus* formed two geographically-define lineages: Eastern pear and Western pear (Rubtsov 1944). Lo and Donoghue (2012) estimated the age of *Pyrus*, determining the MRCA age to be around 27 or 33 Mya. This aligns with the findings of Korotkova et al. (2018), who observed an earlier divergence time for East Asian pear compared to Western pear, suggesting a potential correlation with the decline of the Turgai Strait. Wu et al. (2018) proposed a model outlining the differentiation, spread, and independent domestication of Asian pear and European pear, proposing that pears originated in southwest China, spread across Central Asia, and eventually reached West Asia and Europe.

In our study, we utilize the pear genus, characterized by ILS and hybridization, as a model to develop a comprehensive pipeline for untangling the complexities of the Web of Life. This innovative methodology will enable us to address three critical objectives to deepen our understanding of the pear genus: 1) establish a robust phylogenetic backbone for *Pyrus*, utilizing a carefully curated dataset comprising 801 nuclear SCN genes and 77 plastid CDSs; 2) explore the evolutionary mechanisms that have driven the dynamics of pear species; 3) elucidate the biogeographical origin of *Pyrus* given significant gene tree and cytonuclear discordance.

## Results

### Comprehensive screening and assembly of nuclear genes and plastome datasets

In this study, our assembly of 801 SCN genes revealed an uneven retrieval of genes across all samples, ranging from 619 (77.3%) to 801 (100%) genes (supplementary fig. S1; supplementary table S2). Despite stringent selection criteria for SCN gene screening, the ‘hybpiper paralog_retriever’ from HybPiper v. 2.0.1 (supplementary fig. S2) revealed the presence of paralogs in some genes across various Maleae samples. A substantial fraction of genes (330) exhibited paralogous sequences in at least one sample. To reduce the effects of paralogs and enhance the signal-to-noise ratio, two distinct datasets of orthologous genes were generated: 771 MO orthologs and 905 RT orthologs, using the tree-based orthology inference method. Following a series of rigorous data filtration steps, the final alignments of these two datasets yielded genome-level sequences of 1,040,446 and 1,197,809 bp, respectively.

A total of 72 plastomes were successfully assembled to complement an additional 20 plastomes sourced from GenBank. Of these, 29 samples underwent assembly utilizing either NOVOPlasty or GetOrganelle, whereas the remaining 43 samples, characterized by lower-quality data, were processed using the SARD assembly method. All 92 plastomes were incorporated into the subsequent phylogenetic analyses. Plastid CDSs were extracted from the assembled plastomes, and the final alignment of these plastid CDS datasets spanned a length of 68,280 bp.

### Nuclear/plastid phylogenetic analysis provides insights into clade delineation of *Pyrus*

Phylogenetic inference in *Pyrus* was performed using both concatenated and coalescent-based methods, which were applied to nuclear MO and RT gene datasets. These approaches yielded six phylogenetic trees with nearly congruent topologies (supplementary figs. S3-S8). Given this congruence, the RAxML tree estimated from the nuclear MO dataset was then selected for further analysis (fig. 1a). The nuclear phylogeny indicated that *Pyrus* is monophyletic, forming two highly supported, distinct clades (bootstrap support (BS) = 100, SH-aLRT support/Ultrafast bootstrap (SH- aLRT/UFBoot) = 100/100, and local posterior probabilities (LPP) = 1.0). The phyparts analysis revealed that a substantial proportion of informative gene trees, specifically 377 out of 435 (ICA = 0.697), displayed concordance with the target tree. Furthermore, QS analyses uniformly supported this finding, with all sampled quartet partitions corroborating this node (1/-/1). The sister relationship between *Pyrus* and *Malus*, however, garnered support from only five out of 287 informative gene trees (ICA = 0.019; fig. 1a; supplementary figs. S9, S10), although it was supported in ML (BS = 97; fig. 1a; supplementary fig. S3) and QS analyses (0.011/0.88/0.97; fig. 1a; supplementary fig. S11).

**Figure 1.**
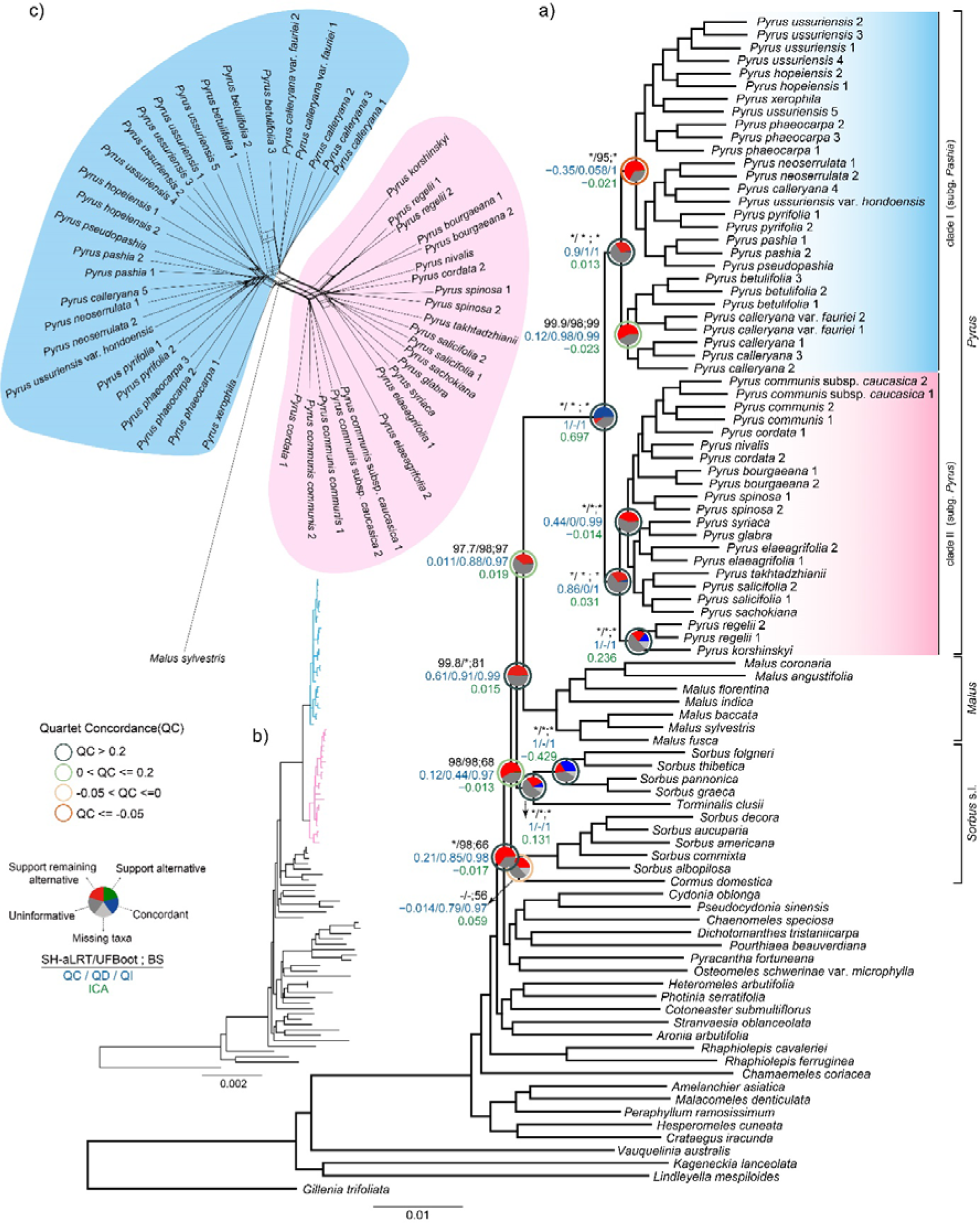
**a)** Maximum likelihood (ML) phylogeny of *Pyrus* in the framework of Maleae inferred from RAxML analysis using the concatenated 771 nuclear MO orthologs supermatrix. Summarized phylogenetic supports of the focal nodes from two trees based on the nuclear MO dataset are presented above the branch. From left to right (labeled in black above branch), the SH-aLRT support and Ultrafast Bootstrap estimated from IQ-TREE2 (details referring to Supplementary Fig. S4); the bootstrap support (BS) values from RAxML analysis (details referring to Supplementary Fig. S3) (e.g., 97.7/98; 97); asterisks (*) indicates full support (100/100; 100). Pie charts on the nodes represent the following data: the proportion of gene trees that support that clade (blue), the proportion that support the main alternative bipartition (green), the proportion that support the remaining alternatives (red), the proportion (conflict or support) that have less than 50% bootstrap support (BS, dark grey), and the proportion that have missing taxa (light grey). The value of partial sampling ICA is presented below (labeled in green) (details referring to Supplementary Fig. S9). The color of the circle around the pie chart represents the value range of Quartet Concordance (QC), where QC > 0.2 is painted in dark green, 0 < QC ≤ 0.2 is painted in light green, -0.05 < QC ≤ 0 is painted in yellow, and QC ≤ -0.05 is painted in red. Values for Quartet Concordance/ Quartet Differential/ Quartet Informativeness estimated from QS analysis are provided below branches (e.g., 1/-/1, labeled in blue) (details referring to Supplementary Fig. S11). **b)** RAxML tree of *Pyrus* in the framework of Maleae using the concatenated 92 CDS supermatrix. Branches in blue indicated clade I and pink indicates clade II (details referring to Supplementary Fig. S12). **c)** Supernetwork inferred with SplitsTree based on 771 rooted Maximum likelihood (ML) gene trees, where parallelograms indicate incongruences among gene trees. Clade I is shown in blue, clade II is depicted in pink.

The monophyly of the genus *Pyrus* also received robust support, with two highly supported clades in plastid phylogenetic relationships (BS = 100; SH-aLRT/UFBoot = 100/100; LPP = 1.0; fig. 1b; supplementary figs. S12-S14). In addition, concordance between ML tree and informative gene trees was significant, with 17 out of 21 gene trees exhibiting consistency (ICA = 0.533; supplementary figs. S15-16), and full QS support (1/-/1; fig. 1b; supplementary fig. S17). However, monophyly among species with multiple sampled individuals did not receive support. Moreover, nodes in the plastid phylogeny were not well-resolved in these two clades, and strong gene tree conflicts and weak QS scores were prevalent in our conflict analyses (supplementary figs. S15-S17). Branch lengths were also extremely short in most terminal nodes, which may be due to the limited informative sites in the plastid CDS sequences.

### Conflict analyses suggest reticulate relationships between *Pyrus* and its related genera

Conflicts were observed in the relationships between *Pyrus* and related genera across datasets and tree-inference methods (fig. 2a). An examination of nuclear phylogenies inferred using ASTRAL-III revealed a sister relationship between *Pyrus* and a large clade comprising *Aria*, *Micromeles*, and *Torminalis*, albeit with relatively low support (LPP = 0.35 for MO in supplementary fig. S5, LPP = 0.33 for RT in supplementary fig. S8). Furthermore, only eight out of 283 informative gene trees were congruent with this species tree topology (ICA = 0.697), as determined by the phyparts analysis (supplementary figs. S18-19). The QS result also indicated counter-support (-0.084/0.5/0.97; supplementary fig. S20). Conversely, concatenation-based trees strongly supported *Pyrus* as sister to *Malus* (BS = 97; SH-aLRT/UFBoot = 97.7/98; fig. 2a; supplementary figs. S3-S4). Plastome-derived phylogenies recovered a sister relationship between *Pyrus* and a combined clade consisting of *Sorbus* s.s., *Micromeles*, and *Cormus* within concatenation-based frameworks (BS = 91; SH- aLRT/UFBoot = 97.8/100, supplementary figs. S12-S13), while *Pyrus* inferred as sister to *Cormus* in the coalescent-based species tree (LPP = 0.63, supplementary fig. S14). The phylogenetic relationships among the five clades within *Sorbus* s.l. varied significantly, particularly concerning the positions of *Sorbus* s.s. and *Cormus* (fig. 2a; supplementary figs. S3-S8, S12-S14).

**Figure 2.**
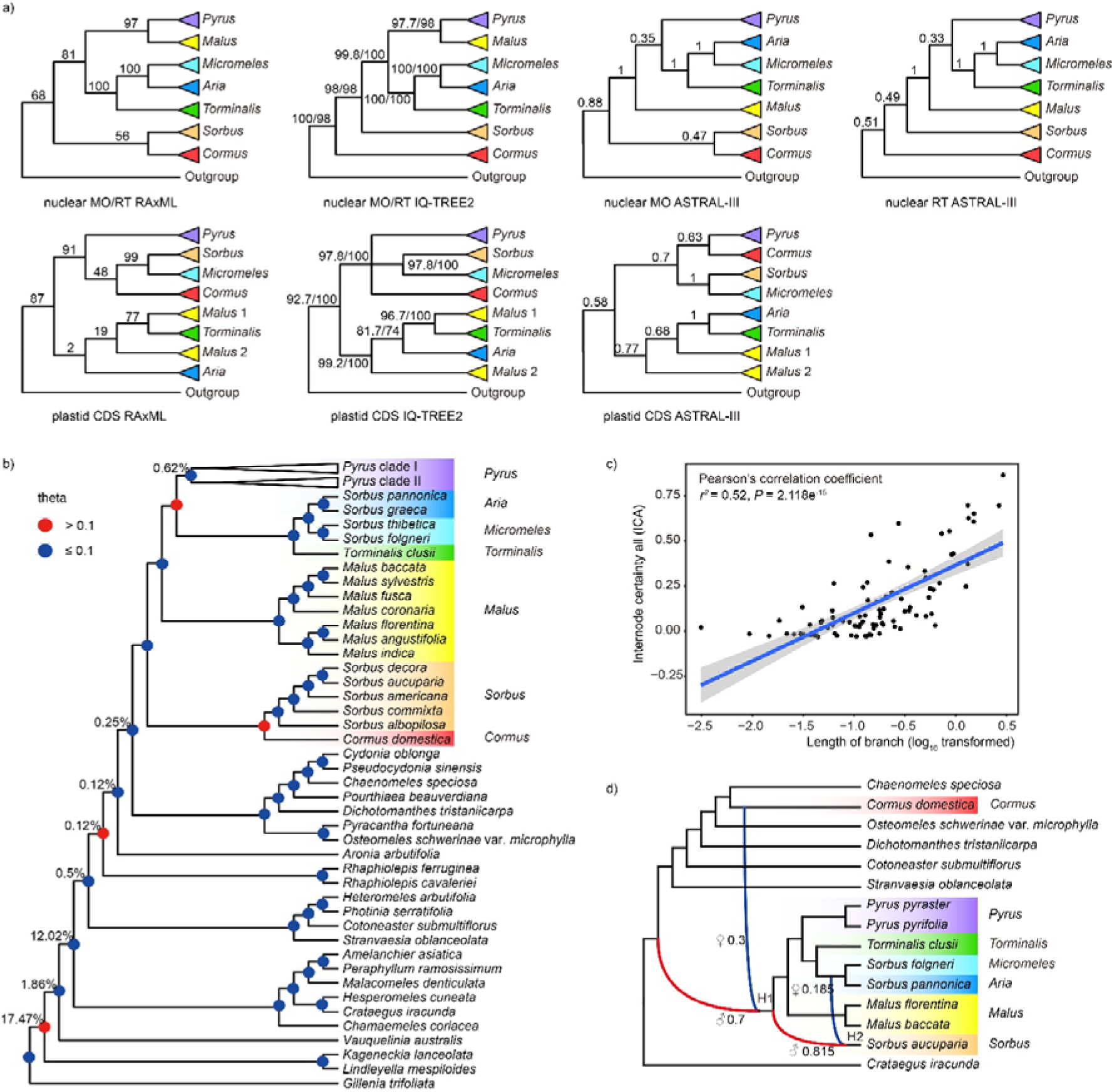
**a)** Comparative visualization of conflicting topologies from different datasets and inference methods. Phylogenetic supports of the focal nodes from trees are presented next to the branch. The bootstrap support (BS) values from RAxML analysis (e.g., 97; details referring to Supplementary Fig. S3); the SH-aLRT support and Ultrafast Bootstrap estimated from IQ-TREE2 (e.g., 97.7/98; details referring to Supplementary Fig. S4); the local posterior probability (LPP) from ASTRAL-III (e.g., 0.35; details referring to Supplementary Fig. S5). For MO/RT nuclear trees with the same topology recovered by different tree-building methods, only the support value from the MO dataset is shown. **b)** A cladogram of Maleae from ASTRAL analysis of MO ortholog trees with nodes colored by population mutatuin parameter theta. Percentages next to nodes denote the proportion of duplicated genes when using orthogroups from the homologs (details referring to Supplementary Fig. S21). **c)** The correlation between length of branch and internode certainty (ICA) value. **d)** Phylogenetic network analysis from the 15-taxa sampling of Maleae. Blue and red curved branches indicate the possible hybridization events with the corresponding inheritance probabilities from the parental lineages marked beside.

### Comparative analyses emphasize the reticute relationship within *Pyrus*

*Pyrus* was delimited into two major clades with maximum support (BS = 100, 100; SH- aLRT/UFBoot = 100/100, 100/100; LPP = 1.0, 1.0), designated as clade I (subg. *Pashia*) and clade II (subg. *Pyrus*) for the convenience of discussion. These clades demonstrated a pronounced association with their respective geographical distributions, further substantiated by a global network analysis conducted using SplitsTree (fig. 1c).

Clade I, an East Asian clade, is further subdivided into three subclades (fig. 3a). Subclade A comprises four species predominantly from northern China: *Pyrus ussuriensis*, *P. hopeiensis*, *P. xerophila*, and *P. phaeocarpa*. Notably, with the exception of one individual of *P. ussuriensis* collected in Japan, which is sister to *P. xerophila*, all species within this subclade are monophyletic. Subclade B includes six species, four from southern China and two from Japan. Subclade C consists of *P. calleryana* and *P. betulifolia*, widely distributed across China, along with the Korean endemic *P. calleryana* var. *fauriei*. Phylogenetically, subclade A is sister to subclade B, and these two subclades collectively are sister to subclade C. This specific topology is maximally supported, by bootstrap values, SH-aLRT/UFBoot scores, and LPP (100; 100/100; 1.0), and is further reinforced by strong QS support (0.9/1/1). Nonetheless, this topology is corroborated by only a minority of informative gene trees—nine out of 249, with an ICA score of 0.013. Moreover, the phyparts analysis did not support a sister relationship between subclades A and B, as reflected by a negative ICA score (-0.021). This dissent is mirrored in the QS results, which show counter-support (-0.35/0.058/1). These conflicts are manifested in the coalescent-based tree (right panel in fig. 3a), where the monophyly of subclades A and B is not evident in the species tree.

**Figure 3.**
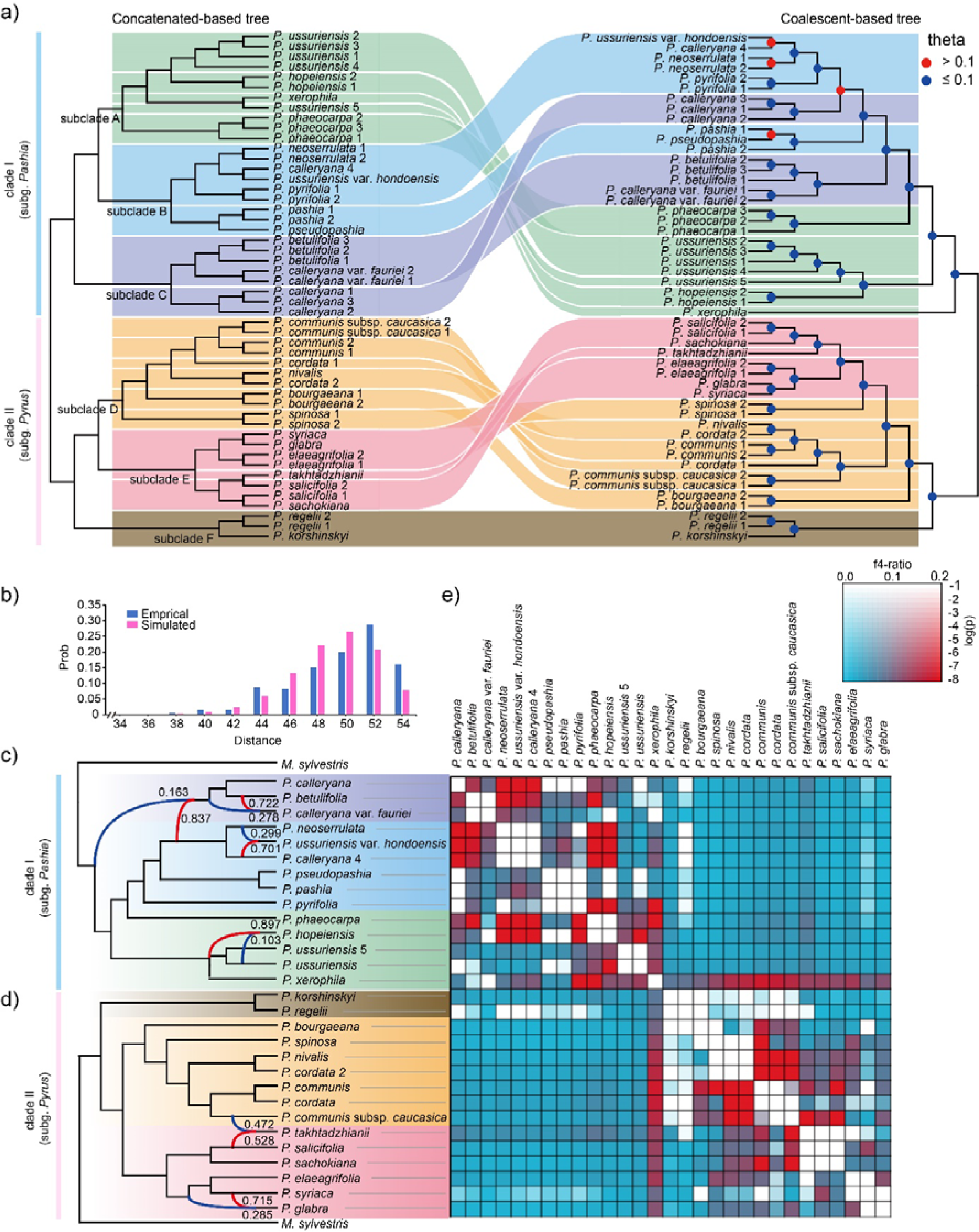
**a)** Comparison of the species tree and gene tree topologies based on nuclear MO orthologs of the *Pyrus* with 50 samples. Maximum likelihood (ML) tree inferred from a supermatrix of 771 concatenated MO orthologs with RAxML (Concatenation-based tree; Left), and the species tree topology inferred from individual nuclear gene trees using a multispecies summary coalescent approach performed with ASTRAL-III (Coalescent-based tree; Right). The pear genus was divided into two clades (clade I in blue and clade II in pink) and six subclades (subclade A in lime green, subclade B in soft blue, subclade C in grayish blue, subclade D in soft orange, subclade E in soft red, subclade F in dark moderate). **b)** Distribution of tree-to-tree distance between empirical gene trees and the ASTRAL species tree, compared to those from the coalescent simulation. **c)** Phylogenetic network analysis from the 15-taxa sampling of clade I within *Pyrus* and one outgroup. Blue and red curved branches indicate the possible hybridization events with the corresponding inheritance probabilities from the parental lineages marked beside. **d)** Phylogenetic network analysis from the 16-taxa sampling of clade II within *Pyrus* and one outgroup. Blue and red curved branches indicate the possible hybridization events with the corresponding inheritance probabilities from the parental lineages marked beside. **e)**. Heat map showing statical support for gene flow between pairs of species inferred from *D*suite package. The shaded scale in boxes represents the estimated f4-ratio branch value.

The Occidental Clade II is further delineated into three subclades with full support (100, 100, 100; fig. 3a). Subclade F is phylogenetically sister to a clade comprised of D and E, a topology substantiated by 27 out of 257 informative gene trees (ICA = 0.031) and strong QS support despite a discordant skew (0.86/0/1). Subclade F comprises species endemic to Central Asia, namely *P. regelii* and *P. korshinskyi*. Meanwhile, five species from Europe & North Africa—*P. communis*, *P. cordata*, *P. nivalis*, *P. bourgaeana*, and *P. spinosa*—along with the Caucasian species *P. communis* subsp. *caucasica*, constitute subclade D. Subclade E is comprised of six species: *P. syriaca*, *P. glabra*, *P. elaeagrifolia*, *P. takhtadzhianii*, *P. salicifolia*, and *P. sachokiana*.

However, the phylogenies within *Pyrus* constructed using nuclear gene datasets with different phylogenetic methods also showed significant discordance (fig. 3a). In clade I, our phylogenetic analysis revealed an alternative topology in the species tree (right panel in fig. 3a; supplementary fig. S5), which did not exhibit three subclades as recovered by RAxML (left panel in fig. 3a; supplementary fig. S3). In the coalescent-based tree, *P. xerophila* formed a sister relationship with all other species in clade I (fig. 3a; supplementary fig. S5). This topology was corroborated by eight out of 248 informative trees (ICA = 0.013; supplementary figs. S18-S19) from the phyparts analysis and received strong QS support with discordant skew (QS = 0.67/0/1; supplementary fig. S20). The phylogenetic position of *P. xerophila* revealed significant conflicts between the coalescent-based and concatenation-based trees, suggesting its potential hybrid origin. The subclade in clade I, except *P. xerophila*, was then delineated into two major clades. In the first major clade, *P. ussuriensis* is identified as the sister species to *P. hopeiensis*, with robust phylogenetic support (LPP = 1 in supplementary fig. S5; no concordant tree out of 357 informative trees, ICA = -0.021 in supplementary figs. S18- S19; QS = 0.37/0.059/0.98 in supplementary fig. S20). The concatenation-based tree further corroborates this relationship. Furthermore, both species, predominantly found in northern China, exhibit a high degree of morphological similarity. The other clade was composed of three individuals of *P. phaeocarpa* (fig. 3a, in lime green; LPP = 1 in supplementary fig. S5; 10 concordant trees out of 286 informative gene trees, ICA = 0.037 in supplementary figs. S18-S19; QS = 0.17/0/0.99 in supplementary fig. S20) and a large clade.

In clade II, the central Asian clade, *P. regelii* and *P. korshinskyi* were consistently recovered as sister to all other species in both the concatenated and coalescent-based trees (fig. 3a). Notably, significant conflict was observed in the phylogenetic relationship of the species indigenous to Europe & North Africa and West Asia (fig. 3a). In the species tree, species endemic to West Asia (except *P. communis* subsp. *caucasica*) were monophyletic with strong support (LPP = 0.98) and an abundance of conflict trees (433 out of 434 informative gene trees, ICA = -0.023; QS = 0.28/0.55/0.97) (fig. 3a, color in soft red; supplementary figs. S5, S18-S20). However, this West Asian clade was observed to be sister to two individuals of *P. spinosa* (right panel, fig. 3a), primarily distributed in Europe, instead of forming a sister relationship with the entire Europe & North Africa clade as seen in the concatenation-based tree. Two species of *P. communis* and *P. communis* subsp. *caucasica*, were not sister, as indicated in the concatenation-based tree. *Pyrus communis* was sister to a lineage containing *P. nivalis* and *P. cordata* 2, and all three species were successively sister to *P. cordata* 1 and *P. communis* subsp. *caucasica.* Two individuals of *P. bourgaeana* formed one distinct clade.

This species is primarily distributed in North Africa. *Pyrus takhtadzhianii* was sister to one individual of *P. salicifolia* with moderate support (BS = 87; 19 concordant trees out of 317 informative gene trees, ICA = 0.055), and QS counter-support (-0.52/0.19/1) in concatenation-based trees (fig. 3a; supplementary figs. S3, S9-S11). However, *Pyrus takhtadzhianii* was recovered as a sister lineage to a clade composed of *P. salicifolia* and *P. sachokiana* with maximum support (LPP = 1), a few concordant trees (six out of 377 informative gene trees, ICA = 0.0331) and strong QS support with discordant skew (0.63/0/1) in coalescent-based phylogenies (fig. 3a; supplementary figs. S5, S18-S20).

### Incomplete lineage sorting analysis highlights its role in the origin of *Pyrus*

Our findings revealed a significant increase in theta value (theta = 0.778) in the node clustering *Pyrus* and the clade containing *Aria*, *Micromeles*, and *Torminalis*, indicating the significant impact of ILS in the diversification of this large clade. Furthermore, a notable theta value (theta = 0.172) was observed at the node of the *Sorbus* and *Cormus* lineages (fig. 2b; supplementary fig. S21). Regression analysis conducted across the Maleae tribe revealed a weak positive correlation (Pearson’s correlation coefficient *r*^2^ = 0.52, *P* = 2.118e^-15^; Pearson’s correlation test, two-sided) between branch lengths and ICA values (fig. 2c). This correlation partially consistent with the expectation that ILS contributed to the genetic discrepancies observed within *Pyrus*.

However, nodes exhibiting low theta values (value < 0.1) were predominantly located within the *Pyrus* lineage (fig. 3a; supplementary fig. S21). Complementary ILS analysis, employing intraspecific coalescent-based simulations, demonstrated minimal overlap between empirical and simulated distance distributions (fig. 3b). This suggests that ILS alone may not fully account for the genetic incongruence detected in *Pyrus* based on analyses of nuclear SCN gene datasets.

### Network and gene flow analysis reveal the intricate gene flow within *Pyrus*

In the network inference using the “Maleae 15-taxa data” at the genus level, we identified two instances of hybridization based on pseudodeviance score ranking (fig. 2d; supplementary fig. S22). The network analysis revealed a hybridization event between MRCA of *Cormus domestica* and the MRCA of a diverse clade encompassing *Aria*, *Chaenomeles*, *Cormus*, *Cotoneaster*, *Dichotomanthes*, *Malus*, *Micromeles*, *Osteomeles*, *Pyrus*, *Sorbus* s.s., *Stranvaesia*, and *Torminalis*, with γ values of 0.3 and 0.7, respectively. Another detected hybridization event suggests a hybrid origin for *Sorbus* s.s., involving the MRCA of *Micromeles* and *Aria* (γ = 0.185) and the MRCA of a lineage that includes *Pyrus*, *Torminalis*, *Micromeles*, *Aria*, *Malus*, and *Sorbus* s.s. (γ = 0.815).

Previous research has highlighted the crucial role of hybridization in the evolutionary history of *Pyrus*. To further explore this, we included 29 individuals representing all *Pyrus* species and one outgroup species (*Malus sylvestris*) for phylogenetic network inference (fig. 3c,d). Within Clade I, utilizing the “*Pashia* 15-taxa data”, the SNaQ network analysis identified four optimal hybridization events (fig. 3c; supplementary fig. S23). This analysis suggested that the MRCA of *P. calleryana*, *P. betulifolia*, and *P. calleryana* var. *fauriei* likely resulted from hybridization between the MRCA of *P. neoserrulata*, *P. ussuriensis* var. *hondoensis*, and *P. calleryana* 4 (83.7%) and the MRCA of *Pyrus* subg. *Pashia* (16.3%).

Additionally, the origin of *P. ussuriensis* var. *hondoensis* and *P. hopeiensis* was inferred to involve hybridization events, indicating complex genetic relationships within this clade.

In Clade II, analyzed with “*Pyrus* 16-taxa data”, two hybridization events were detected according to SNaQ network analysis (fig. 3d; supplementary fig. S24). *P. takhtadzhianii* appeared to have genetic contributions from *P. communis* subsp*. caucasica* (47.2%) and *P. salicifolia* (52.8%). Another event involved *P. syriaca* and the ancestral clade of *P. elaeagrifolia* and *P. syriaca*, contributing significantly to the genetic material of *P. glabra*.

Further exploration using the f4-ratio statistics in *D*suite revealed multiple introgression events within *Pyrus* species, corroborating the hybridization events suggested by the phylogenetic network analyses (fig. 3e supplementary fig. S25). These gene flow instances, particularly among clades I and II, underscore the significant impact of gene introgression on the phylogenetic inconsistencies observed between concatenated and coalescent-based trees, emphasizing the complex evolutionary dynamics within the *Pyrus* genus.

### Polyploidy analysis reveals the minimal contribution of polyploidization to the origin of *Pyrus*

In response to the frequent association between polyploidization events and hybridization, we employed on a WGD analysis. By mapping duplication events identified within orthogroups onto the MO orthology trees, we observed a marked increase in gene duplication rates at several nodes among the basal groups of the Maleae tribe (fig. 2b).

Notably, only 0.62% of the 807 genes analyzed showed signs of duplication within the clade that encompasses the MRCA of *Pyrus*. To further discern the haplotype structure of the species, we analyzed heterozygous k-mer pairs using Smudgeplot. The prevalent distribution of AB k-mers indicated that the genomes of *Pyrus* species are predominantly diploid (supplementary fig. S26). These findings suggest that polyploidy did not significantly contribute to the origin or speciation of *Pyrus* species.

### Biogeographic analyses revolutionize our understanding of *Pyrus* origin

The divergence time was estimated using SCN genes and plastid CDS datasets, and the resulting time trees were used to conduct ancestral area reconstruction with the biogeography models based on the AICc_wt value (supplementary figs. S27-S32). According to the nuclear gene dataset, *Pyrus* originated from East Asia, Central Asia, and West Asia in the early Oligocene (29.72 Mya, 95% HPD: 27.6-31.84 Mya) and then diversified in the early Miocene (21.85 Mya, 95% HPD: 19.36-24.41 Mya; fig. 4; supplementary figs. S27-29) due to the vicariance. The crown clade of *Pyrus* subg. *Pyrus* formed around 19.32 Mya (95% HPD: 17.09-21.66 Mya); the crown clade of *P.* subg. *Pashia* was estimated to have diversified between Central Asia and West Asia in the middle Miocene (17.61 Mya, 95% HPD: 15.22- 20.06 mya; fig. 4; supplementary figs. S27-29). Some species further diversified in Europe & North Africa after the Pliocene (5.27 Mya, 95% HPD: 3.04-7.74 Mya; fig. 4; supplementary figs. S27-29). The CDS dataset suggested similar origin time and dispersal patterns of *Pyrus* as the nuclear gene dataset, but the divergence time of most lineages within the *Pyrus* was much younger (supplementary figs. S30-S32).

**Figure 4.**
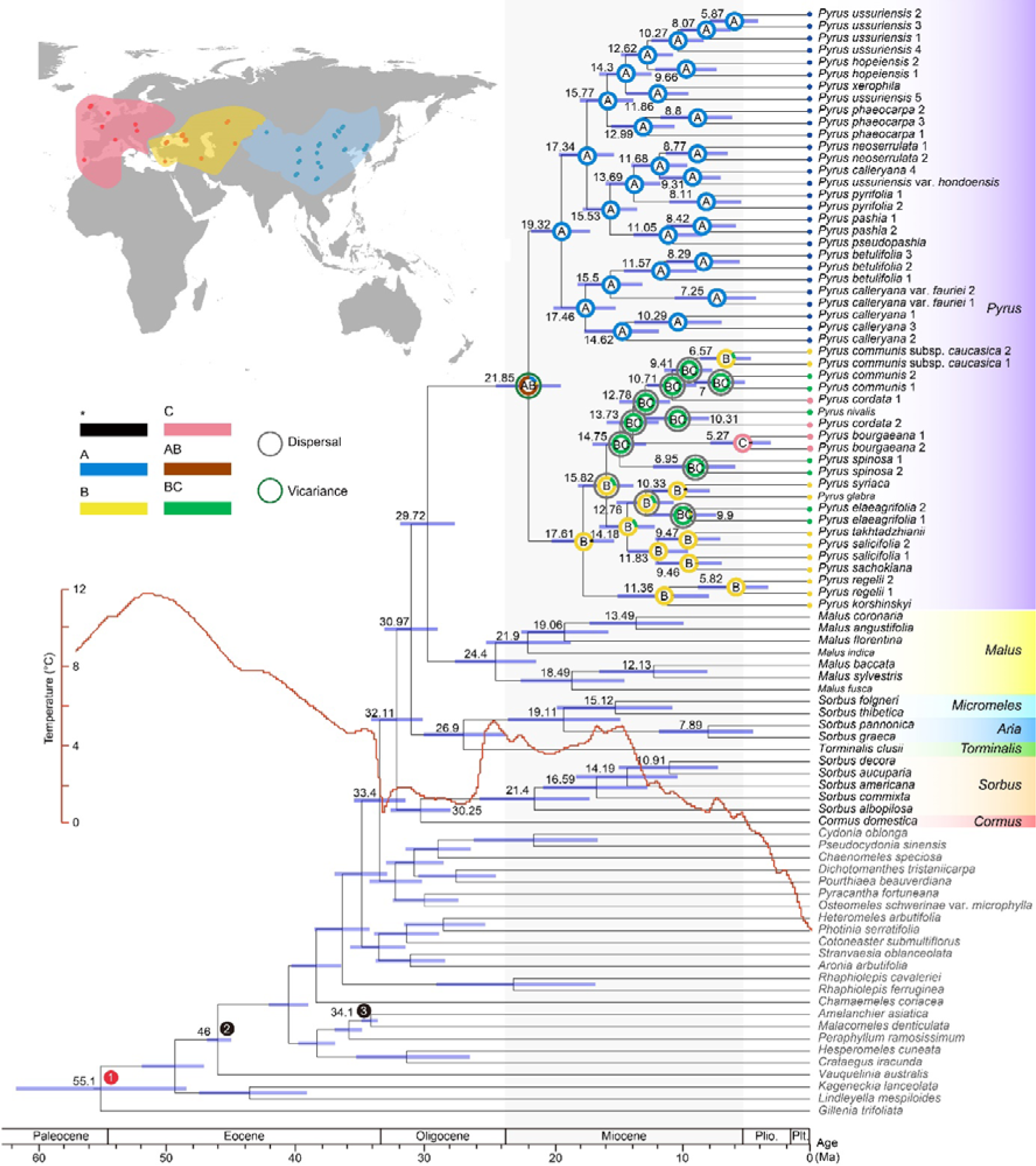
Divergence time estimation and geographical range evolution of the Maleae. Dated chronogram of the Maleae inferred from PAML based on the nuclear MO dataset. Focal nodes feature estimated divergence times and ancestral geographical ranges. The inset map in the upper left outlines the three distribution areas used for geographical analysis. (A), East Asia (blue); (B), Central Asia and West Asia (yellow); (C), Europe and North Africa (pink). The dots on the map indicate the geographical origin of species. Two fossils, colored black (nodes 2-3), and one divergence time estimate based on previous research, colored red (node 1), are used as constraints. The red line represents the global temperature changes (from Westerhold et al., 2020).

We conducted BAMM analysis utilizing dated trees inferred from nuclear SCN gene and plastid CDS datasets, respectively (fig. 5). The analysis based on CDS dataset showed a pattern of increase in the net diversification and speciation rate (fig. 5a). In contrast, the speciation rate and the net diversification rate, as estimated from nuclear datasets, demonstrated an decreased incline (fig. 5b). Although the extinction rates remained relatively stable in both datasets, a higher rate was observed in the CDS dataset (fig. 5).

**Figure 5.**
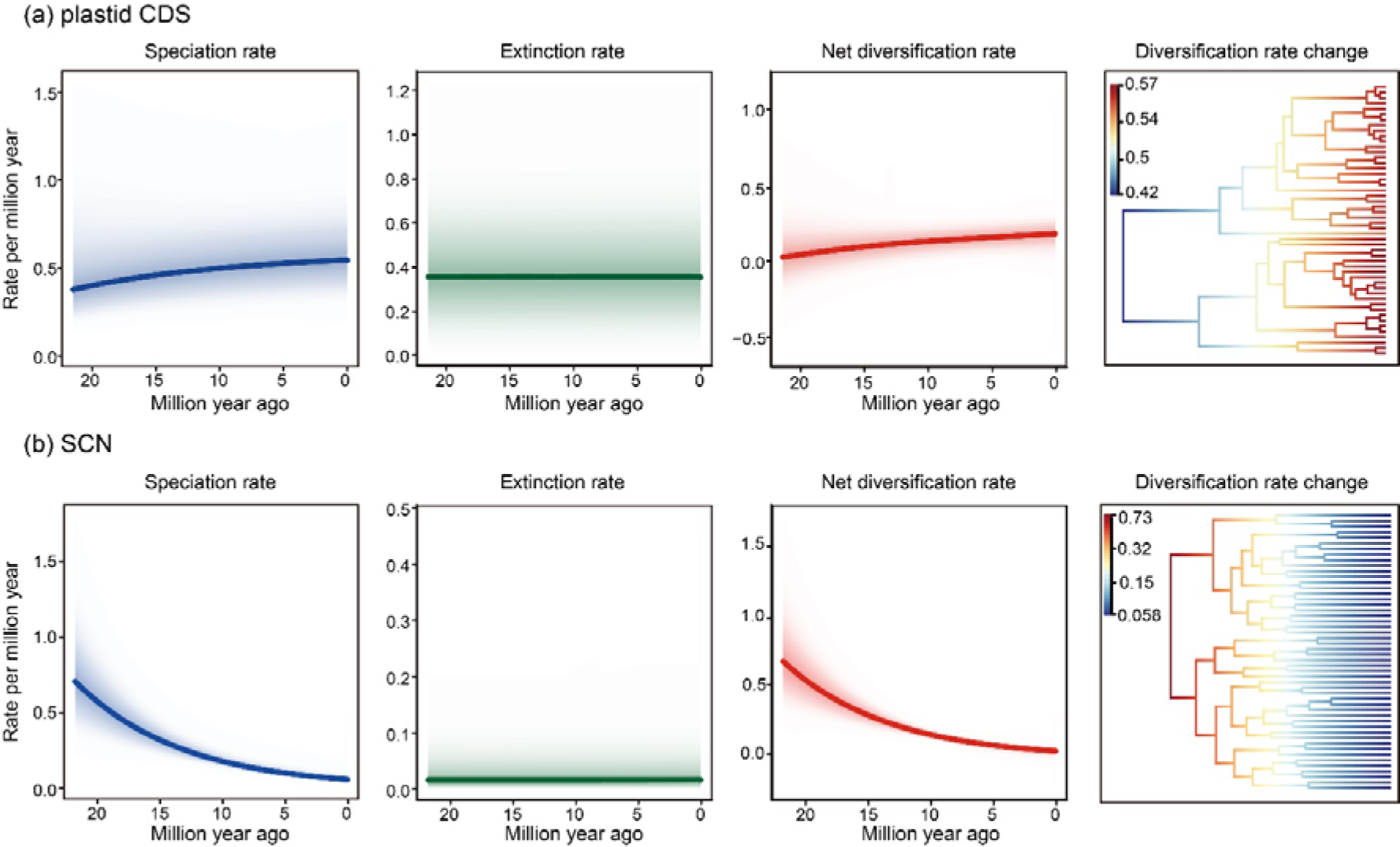
Speciation rates, extinction rates, net diversification rates and changes in speciation rates among major lineages within *Pyrus* over time base on BAMM analysis. Shaded regions show the 90% credible intervals for the rate. **a)** BAMM analysis based on CDS dataset. **b)** BAMM analysis based on nuclear MO dataset.

## Discussion

### A Step-by-Step Exclusion approach for unraveling the Web of Life

Untangling the evolutionary histories of plant lineages remains a major objective within the field of systematics; however, achieving this requires distinguishing between the various mechanisms leading to reticulation. Contemporary analytical programs typically analyze individual factors in isolation (Morales-Briones et al., 2021), a practice that hinders a comprehensive understanding of evolution due to the absence of an integrated framework for examining multiple factors simultaneously. To address this limitation, we propose a Step-by-Step Exclusion (SSE) approach, designed to unravel the complexities inherent in the Web of Life (WoL) (Fig. 6). This study focuses on the pear genus, known for its significant hybridization driven by self-incompatibility, to explore its complex evolutionary relationships and potential paleobiogeographical history. When phylogenetic discordance within the focused lineage was identified, the SSE approach was applied to investigate the potential evolutionary mechanisms driving diversification, adhering to a step-by-step pipeline based on the if-not principle. After bifurcating phylogenetic inference using nuclear and plastid datasets, we assessed the extent of conflict among nodes using phyparts and QS metrics. Comparisons between nuclear and plastid trees were conducted to detect potential chloroplast capture events. Moreover, two alternative approaches were employed to assess the role of ILS: coalescent simulation analysis (Liu et al., 2022) and analysis based on the population mutation parameter theta (Cai et al., 2021), each providing insights into deep coalescence from a global perspective or focusing on specific nodes, respectively.

**Figure 6.**
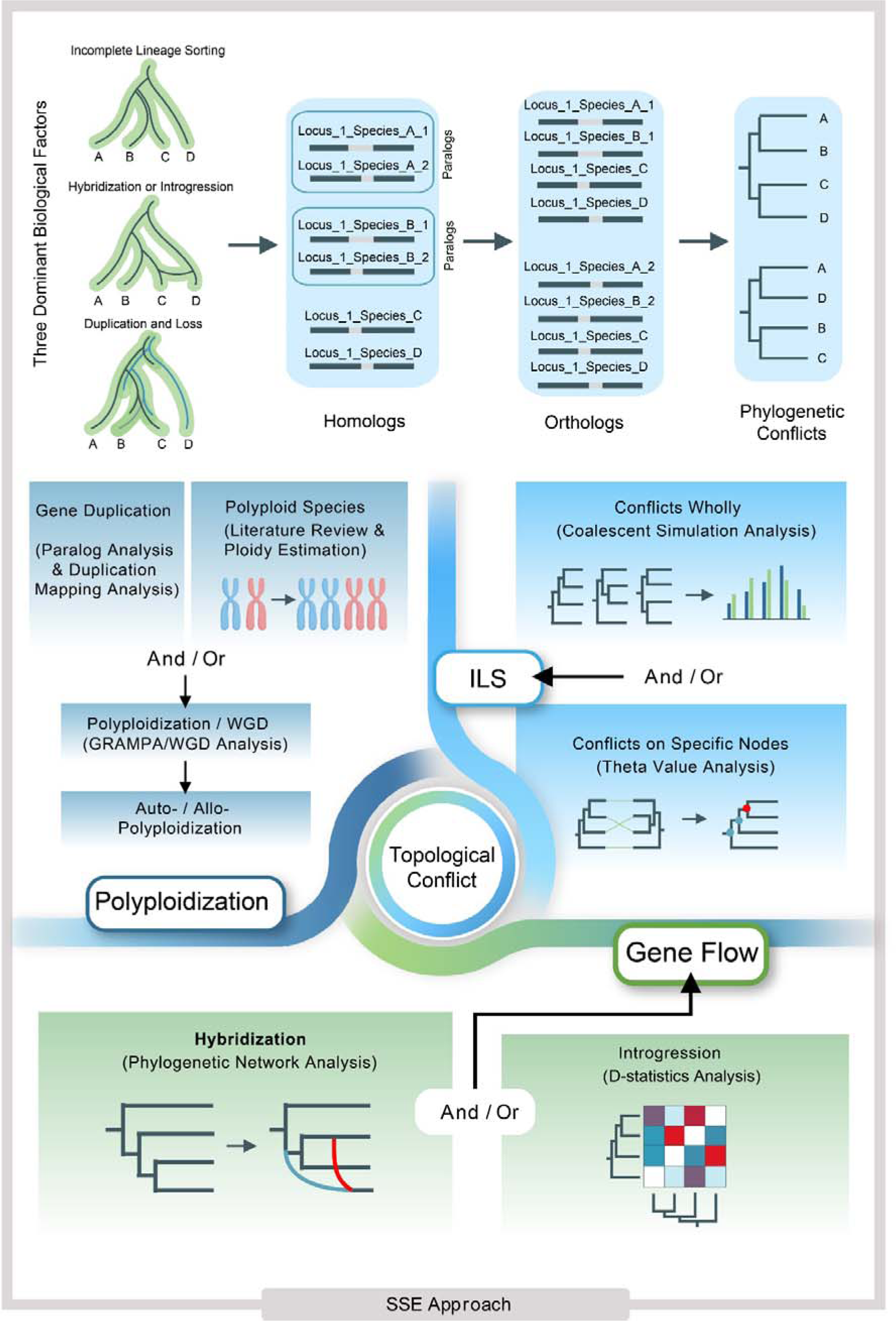
Workflow of the Step-by-Step Exclusion (SSE) approach for unraveling the complexities inherent in the Web of Life. An illustrated evolutionary mechanism for generating phylogenomic conflicts is presented (top). These three biological factors lead to the admixture of genetic information among species, particularly through paralogs, thereby complicating the identification of evolutionary relationships among species. The step-by-step pipeline (bottom), based on the if-not principle, individually analyzes the role of these three factors in the evolutionary process and infers the evolutionary history of the group by synthesizing multiple lines of evidence.

Ploidy estimation, from the literature or using newly collected data, enables a preliminary insight into the historical occurrences of polyploidization/WGD within the focal lineage. Subsequent analyses for detecting paralogs and mapping gene duplication events onto phylogenetic trees provide additional evidence of polyploidization/WGD events. GRAMPA and WGD analyses can further substantiate occurrences of autopolyploidy or allopolyploidy. The potential for hybridization-driven reticulation is assessed through tools such as PhyloNetworks, PhyloNet, and *D*-statistics, which elucidate the reticulate relationships among samples. By constructing an implicit phylogenetic network and analyzing gene flow, we can infer possible patterns of hybridization/gene flow. The temporal and geographic context of these reticulation events may be deduced through historical biogeographic analysis.

This sequential analytical approach enables a thorough investigation of the multifaceted factors shaping the intricate evolutionary history of taxa, providing a robust framework for elucidating the evolutionary narratives of complex groups. Crucially, our case study involving *Pyrus* revealed that despite frequent genome doubling in the Maleae, the SSE approach identified ILS, and not polyploidy, better explains the diversification of the pear genus. This study established a foundational analytical framework leveraging DGS datasets, suggesting that ongoing advancements in software development could further enhance and refine this framework, leading to increasingly accurate outcomes in future studies.

### ILS, rather than polyploidization, drives the origin of *Pyrus*

Phylogenetic studies have repeatedly recovered an ambiguous phylogenetic placement of *Pyrus*, with some studies positing it is sister to *Malus* (Xiang et al. 2017; Liu et al. 2022), while others suggest a closer relationship to *Sorbus* s.l. (Lo and Donoghue 2012; Zhang et al. 2017; Sun et al. 2018; Liu et al. 2022; Zhang et al. 2023). Recent improvements within the Maleae clade have verified cytonuclear discordance in *Pyrus* (Liu et al. 2022; Jin et al. 2023), with our findings further validating topological discordance across diverse datasets using various tree inference methodologies (fig. 2a). To measure the potential influence of non-biological factors as underlying causes for the observed genealogical and cytonuclear conflict regarding the position of *Pyrus*, an array of methodical strategies have been deployed to minimize these analytical impacts. These strategies encompass comprehensive taxon sampling of *Pyrus* and its closest relatives, application of a tree-based orthology inference method, adoption of complementary tree inference techniques based on concatenated and coalescent-based methods, and a series of sequence processing approaches. Subsequent analyses will be dedicated to investigating the evolutionary mechanisms attributable to biological factors.

Analysis of nuclear gene tree discordance revealed that the sister relationship between *Pyrus* and *Malus* was supported by only five out of 287 gene trees (supplementary fig. S9), albeit with substantial BS support in the ML tree (97.7/98; 97; fig. 1a; supplementary figs. S3- 4). This observation is consistent with the QS result, indicating only a weak majority of from inadequate informative sites (fig. 1a). Phylogenetic network analyses have detected two distinct hybridization events between *Pyrus* and its closely related genera within the Maleae tribe (fig. 2d; supplementary fig. S22). Nonetheless, neither event directly involved *Pyrus*, unveiling a complex evolutionary pattern within *Sorbus* s.l. Interestingly, the cytonuclear and genealogical discordances observed in the placement of *Pyrus* (fig. 2a) may be partially clarified through these reticulate events associated with *Sorbus* s.l. Consequently, we can speculate that the hybridization-induced ambiguous position of these five *Sorbus*-related genera indirectly complicates the phylogenetic placement of *Pyrus*. The MRCA of *Pyrus* captured the chloroplast genome of the MRCA of *Cormus* through the H1 hybridization event, which may have occurred in the late Eocene (fig. 4; supplementary figs. S27-S32).

Furthermore, given the widespread occurrence of WGDs across Rosaceae (Xiang et al. 2017), gene duplications and subsequent losses frequently result in the emergence of paralogs post-WGD and/or polyploidization events (Lynch and Conery 2000; Panchy et al. 2016). Despite rigorous selection criteria for SCN genes, paralogous genes were identified in numerous samples within Maleae (supplementary fig. S2). By mapping the MRCA of gene duplication events from orthogroup trees onto the ASTRAL species trees, two nodes within the apple tribe, characterized by an elevated proportion of gene duplications (fig. 2b), were pinpointed as indicative of potential WGD events. This finding corroborates the polyploidy origin hypothesis for the combined clade of Maleae and Gillenieae (Hodel et al. 2022). Conversely, the basal node of *Pyrus* exhibited a minimal gene duplication rate (0.62%), which may reflect *Pyrus*-specific gene duplication and loss events after the polyploidization event, marking the origin of Maleae and Gillenieae. Ploidy estimation was performed via Smudgeplot analysis of heterozygous k-mer pairs in genomes, and all samples were ascertained as diploid (supplementary fig. S26). An exhaustive literature review of chromosome ploidy within *Pyrus* further indicated that all documented wild *Pyrus* species are diploid. In conclusion, the collective evidence presented herein suggests that gene duplication and polyploidization events may not significantly contribute to the origin of *Pyrus*.

ILS has been identified as a principal source of gene tree discordances (Koenen et al. 2020; Feng et al. 2022; DeRaad et al. 2023; Rivas-González et al. 2023), a phenomenon widespread across multi-locus phylogenetic datasets (e.g., Morales-Briones et al. 2022). Despite the challenges in detecting ILS and its often-underappreciated role in facilitating species diversification, documented instances of ILS span diverse phylogenetic groups, including plants, birds, marsupials, and primates. Our comprehensive ILS analyses, employing regression analysis and theta values, suggest that ILS only partially explains the observed conflicts within the Maleae tribe (fig. 2b,c; supplementary fig. S21). Notably, the node associating *Pyrus* with a clade comprising three genera in *Sorbus* s.l. (*Aria*, *Micromeles*, and *Tominalis*) exhibited a high theta value of 0.788 (fig. 2b; supplementary fig. S21), signifying the considerable influence of ILS in the origin of *Pyrus*. The prominent role of ILS has also been observed in the rapid evolution of the crown clade in Fagaceae (Zhou et al. 2022). The anomaly zone, initially defined by Degnan and Rosenberg (2006) as a collection of short internal branches within the species tree indicative of pronounced ILS instances, was later expanded by Rosenberg (2013) to include a requirement for two successive short internal branches in the ancestor–descendant relationship. Our species tree inference from nuclear genes distinctly exhibits these consecutive short internal branches between *Pyrus* and its closely related genera, *Aria*, *Micromeles*, and *Torminalis* (supplementary figs. S5, S8). Based on this observation, it is postulated that ILS predominates as the predominant mechanism influencing the nuclear evolution of *Pyrus*.

Our dating analyses, employing two distinct topologies (nuclear ML tree and plastid CDS ML tree), consistently supports the rapid radiation of *Pyrus* and related genera during the early Oligocene (fig. 4; supplementary figs. S27-28, S3-31). This radiation may be associated with the climate transition during the Eocene-Oligocene transition (EOT) (Zachos et al. 2001). The rapid decline in temperature during this period may have compelled the diversification of the MRCA of *Pyrus*, *Sorbus* s.l., and *Malus*, facilitating their adaptation to extreme environmental conditions via mechanisms such as hybridization and/or ILS (Loiseau et al. 2021; Liu et al. 2024). The ancestral area reconstruction analysis supported the East-to-West Asia origin of *Pyrus*. We concluded that the MRCA of *Pyrus* may have originated from the arid region of Himalayas-Central Asia. In summary, we postulate that the origins of *Pyrus* can be attributed to ILS and rapid radiation, driven by the climatic change during the EOT.

### Frequent hybridization and rampant introgression promoted the diversification of pear species

Although pears have gained considerable attention due to their economic importance as a fruit tree, the evolutionary patterns of *Pyrus* has never been well resolved. Previous studies have shown that the inherent self-incompatibility of pears facilitates widespread interspecific and intraspecific hybridization events within the genus, thereby blurring the morphological boundaries among species (Wu et al. 2018). Furthermore, numerous instances of species non-monophyly have been corroborated across a variety of studies (Kawata et al. 1995; Iketani et al. 1998; Monte-Corvo et al. 2000; Teng et al. 2001, 2002; Bao et al. 2008; Zheng et al. 2008, 2014; Korotkova et al. 2018). However, the evolutionary dynamics responsible for cytonuclear discordance and species non-monophyly within *Pyrus* have not been thoroughly investigated. In the ensuing discussion, we will employ the Step-by-Step Exclusion (SSE) methodology to dissect the evolutionary factors underpinning the diversification of pear species.

All nine trees, inferred from three distinct datasets (two nuclear and one plastid) utilizing three different inference methods (two ML and one species tree), consistently corroborate two major clades in *Pyrus*, i.e., the Occidental (European; *Pyrus* subg. *Pyrus*) and the Oriental (Asian; *P.* subg. *Pashia*) clade. The gene tree discordance analyses robustly validate this bifurcation, with negligible conflict observed within the crown clade of *Pyrus*. The biogeographic analysis supports the diversification of the crown clade of *Pyrus* in the early Miocene, as evidenced by the nuclear ML tree (21.85 Mya, 95% CI: 19.36-24.41 Mya) and the plastid topology (21.48 Mya, 95% CI: 14.81-28.08 Mya). This temporal framework is further corroborated by an exhaustive review of fossil evidence presented in the literature (supplementary table S3). The divergence of these two subgenera is posited to have been influenced by contemporaneous climatic and tectonic shifts, characterized by the retreat of the Paratethys sea, the formation of arid regions in Central Asia, and the orogenic activities near the Tibetan Plateau, culminating in the geographical segregation of East and Central Asia (Harrison et al. 1992; Guo et al. 2002; Xiao et al. 2012).

We used the SSE approach to explore the evolutionary processes underpinning the diversification patterns observed within the two principal clades of *Pyrus*. Polyploidy analysis via Smudgeplot revealed that all specimens were diploid (supplementary fig. S26), thereby negating the influence of polyploidization on *Pyrus* dynamics. Further exploration into gene tree discordance has uncovered significant conflict within *Pyrus* (supplementary figs. S9-S11, S15-S20). The data processing pipeline (referring to the part of orthology inference for the nuclear genes in Materials and Methods) has effectively ruled out the influence of non- biological factors as potential causes for the observed topological discordances. Coalescent simulation analysis, aimed at investigating ILS, demonstrated a significant divergence between the distance distribution predicted by the multispecies coalescent (MSC) model and the empirical distance distribution (fig. 3b). Several nodes exhibited high Theta values (theta > 0.1), suggesting the substantial influence of ILS in accounting for these discordances (supplementary fig. S21). Notably, these nodes with elevated Theta values predominantly pertain to clade I (*Pyrus* subg. *Pashia*). The presence of multiple consecutive short branch lengths provides further evidence supporting the significant contribution of ILS to the pervasive conflicts within the subgen. *Pashia*.

Hybridization influenced the diversification of *Pyrus*, as demonstrated by frequent hybridization events within the genus (i.e., Wu et al. 2018). Investigations into genetic introgression unveiled a considerable degree of genetic material exchange within species of the two major clades. Nonetheless, gene flow between these two clades was not observed, with the exception of *Pyrus xerophila*, which warrants further discussion. Our in-depth analysis of hybridization corroborated numerous events of gene introgression. For instance, *Pyrus hopeiensis* has been identified as a hybrid between *P. ussuriensis* and the MRCA of *P. ussuriensis* and *P. xerophila*. The morphological continuum between *P. hopeiensis* and *P. ussuriensis* poses significant challenges for species delineation. Drawing upon morphological similarity and phylogenomic analysis from SNP data and entire plastid genome sequences, a recent investigation merged *P. hopeiensis* into *P. ussuriensis* (Mu et al. 2022). However, our findings propose an alternative hypothesis, positioning it either as sister to *P. calleryana* var. *fauriei* in the plastid tree or to *P. ussuriensis* in the nuclear tree. This discrepancy may stem from the insufficient taxon sampling in Mu et al. (2022)’s study. This hybridization-driven divergent position of *Pyrus hopeiensis* indicates the need to reassess its taxonomic classification. Moreover, introgression analyses have uncovered extensive gene flow between *Pyrus hopeiensis* and other species of subg. *Pashia*, implying its complex evolution (fig. 3e; supplementary fig. S25). The analysis of genetic introgression suggests that numerous species may have experienced repeated hybridizations and successive backcrosses, leading to disparate genetic compositions (fig. 3c,d,e). In essence, it is postulated that hybridization, coupled with widespread introgression, has been a pivotal driver in the diversification of *Pyrus*.

The progenitor of the Oriental clade may have expanded eastward beyond the Hengduan Mountains in southwestern China, whereas the ancestral lineage of the Occidental clade was situated to the west. These bifurcations led to opposed geographical expansions. The orogeny of the Himalayas and Hengduan Mountains may have played a pivotal role in the genesis of the East Asian monsoon and the Yangtze River water system (He et al. 2021), thereby creating novel environmental niches that facilitated the further diversification of Oriental species. The lack of reproductive barriers and genomic recombination fostered the adaptation to these new environments contributed to speciation in the *Pyrus* subg. *Pashia*, predominantly through ILS and hybridization. Conversely, another lineage, potentially originating in Central Asia, underwent gradual adaptation to an increasingly arid climate; however, rapid radiation evolution did not occur due to unfavorable climatic conditions. Certain species expanded westward as a response to aridification (Sun et al. 2017), diversifying in the Caucasus region, a distribution center of the Occidental clade. With the retreat of the Paratethys sea (Sun et al. 2013) and the consequent establishment of a land bridge between Europe and Asia, the ancestors of Occidental pears further disseminated into Europe, adapting to local climatic conditions and initiating diversification processes. Remarkably, some pear species even migrated to North Africa from Europe via island-hopping strategies. Extensive gene flow, i.e., hybridization, may explain discrepancies in dating time estimations between nuclear and plastid datasets. This assertion is corroborated by temperature fluctuations since 5 Mya (Yu et al. 2023).

In the context of the extensive reticulation observed within wild *Pyrus* species, we identified a notable instance of an artificial hybrid species, *P. xerophila* (fig. 3e). This species exhibits a highly variable position between the concatenated and coalescent-based tree, either nested in subg. *Pashia* or sister to all members of subg. *Pashia* (fig. 3a). Phylogenetic network analysis using Splitstree provided alternative evidence for the intermediate position of this species. Previous research has suggested that *P. xerophila* may possess admixed genetic materials from the Occidental and Oriental groups (Wu et al. 2018). Our genetic introgression analysis revealed that *P. xerophila* contains genetic contributions from more than two pear species, posing significant challenges to the precise identification of its parental lineages. We proposed one possible gene flow pattern for the origin of *P. xerophila* as follows. Considering the geographical distribution of these two major clades, the Oriental pears are predominantly distributed to the east Tianshan Mountains, while the Occidental pears are distributed in Central-West Asia, Europe, and North Africa (Fedorov 1954; Browicz 1993; Zamani et al. 2012). The formation of arid regions in Central Asia may have impeded genetic exchange between these two subgenera, leading to divergent evolutionary trajectories. Given that *P. xerophila*, primarily located in northwest China, incorporates genetic elements from both major clades, it is speculated to have arisen from one or several hybridization events between Oriental and Occidental pears, potentially during the ancient Silk Road era since the Han Dynasty, similar to the case of *Pyrus sinkiangensis*, a confirmed artificial hybrid species (Wu et al. 2018). The lack of intermediate phenotypic traits from both subgenera complicates the determination of its hybridization mechanism, in contrast to *P. sinkiangensis*, which exhibits a pear shape characteristic of Occidental pears and sandy pulp typical of Oriental pears. In comparison, *Pyrus xerophila* predominantly displays morphological traits associated with the Oriental pear group, possibly due to a reduced proportion of genetic material inherited from the MRCA of Occidental pears through successive backcrosses with the paternal lineage (referring to the scenario in Liu et al. 2020).

In conclusion, the two major clades of *Pyrus* have evolved independently. The ongoing orogeny of the Himalayas, coupled with the establishment of an arid zone in Central Asia, has served as a geographical barrier, limiting genetic exchange between the Occidental and Oriental clades. Within these distinct evolutionary histories, ILS and hybridization have been instrumental in facilitating the diversification of Oriental pears, whereas hybridization alone has predominantly driven the diversification of Occidental pears. However, the inception of the Silk Road during the Han Dynasty emerged as a conduit for genetic interchange between the Occidental and Oriental pears, paralleling cultural exchanges. This period saw multiple artificial hybrid species, e.g., *Pyrus xerophila* and *P. sinkiangensis*, underscoring human-mediated cross-breeding and its contribution to the phylogenetic complexity of *Pyrus*.

## Conclusion

In this study, we employed *Pyrus* as a model system to introduce and apply the novel SSE approach, designed to dissect the complexity of the WoL with a focus on deciphering the reticulation obscured by complex genetic interrelations. Our results reveal that ILS, rather than polyploidization, primarily drives the origin of *Pyrus* in the arid Himalayas-Central Asia region, with its evolutionary path closely linked to climatic fluctuations during the EOT, highlighting environmental changes as key to the evolution of *Pyrus*. The research further reveals that two subgenera within *Pyrus* have pursued distinct evolutionary paths, a divergence accelerated by vicariance introduced by the uplift of the Tibetan Plateau and the prevailing aridity in Central Asia. Remarkably, the establishment of the Silk Road during the Han Dynasty emerges as a pivotal conduit for genetic exchange between Occidental and Oriental pears, giving rise to numerous artificial hybrid cultivars. The diversification within the Oriental *Pyrus* subg. *Pashia* is attributed to ILS and hybridization, while the reticulation of Occidental *P.* subg. *Pyrus* is predominantly driven by hybridization.

The SSE approach offers profound insights into the complex evolutionary mechanisms underpinning the *Pyrus* lineage. It underscores the importance of considering both genetic and geographical factors in understanding speciation and diversification processes. This new approach significantly advances our comprehensive understanding of plant evolutionary biology and sets a solid platform for future research into the evolutionary mechanisms that underlie complex plant lineages. Utilizing this approach allows for a deeper insight into the multifaceted evolutionary forces that shape the WoL. It opens avenues for further scholarly work that synergizes historical biogeography with phylogenomic analyses, aiming to elucidate the intricate evolutionary tales embedded within the natural world’s biodiversity.

## Materials and Methods

### Taxon sampling, DNA extraction, and Sequencing

In this investigation, a thorough phylogenetic analysis of the genus *Pyrus* was performed, encompassing all seven subsections as delineated in the comprehensive taxonomic classification by Phipps et al. (1990), which stands as the most detailed taxonomic framework for pear species globally. Given the prevalent self-incompatibility within the genus that leads to the formation of hybrids, we intentionally excluded species recognized as artificial hybrids, such as *P. sinkiangensis*. To accurately estimate the divergence times for the *Pyrus* lineage, a total of 41 outgroups were selected, representing 26 distinct genera in the apple tribe Maleae, covering a broad spectrum of the currently recognized genera. We especially emphasized extensive taxon sampling on the close relatives of *Pyrus*, namely *Malus* and *Sorbus* sensu lato (*Aria*, *Cormus*, *Micromeles*, *Sorbus*, and *Torminalis*), to gain insights into their phylogenetic relationships. Additionally, *Gillenia trifoliata*, belonging to the tribe Gillenieae, was incorporated as an outgroup to root the phylogeny of the tribe Maleae. Of the 92 samples analyzed in this study, 58 were derived from Whole Genome Sequencing (WGS) and/or Deep Genome Skimming (DGS) data obtained from the Sequence Read Archive (SRA) in the National Center for Biotechnology Information (NCBI), while 34 DGS data (2 × 150bp) were generated as part of this research. Detailed accession information regarding these samples is available in the supplementary table S1.

Total genomic DNA extractions were performed using silica-gel dried leaves and herbarium/museum specimens. This extraction employed a modified cetyltrimethylammonium bromide (CTAB) method, known as a mCTAB protocol (Li et al. 2013). DNA extraction was performed in the State Key Laboratory of Plant Diversity and Specialty Crops at the Institute of Botany, Chinese Academy of Sciences (IBCAS). Post-extraction, the integrity and quality of the extracted DNAs were assessed using agarose gel electrophoresis. Following this quality assurance step, DNAs that met the high-quality criteria were transported to the Novogene laboratory in Beijing, China. In this facility, libraries were prepared using the NEBNext^®^ Ultra^™^ II DNA Library Prep Kit. The DNA libraries were then sequenced on the Illumina NovaSeq Platform (2 × 150bp), also located at Novogene, Beijing.

### Reads processing, plastome assembly, and annotation

Raw sequencing reads were trimmed by Trimmomatic v. 0.39 (Bolger et al. 2014). This step removed low-quality bases and trimming adapters that were inadvertently introduced during the NGS sequencing procedure. Subsequently, quality assessment was conducted using FastQC v. 0.11.9 (Andrews 2010); this analysis ensured the reliability and accuracy of the clean data for downstream analyses.

We used a two-step approach to enhance the efficiency and precision of plastome assembly. The first step involved using two automated plastome assembly programs: NOVOPlasty v. 3.6, a seed-based program (Dierckxsens et al. 2017), and GetOrganelle, a toolkit based on *de novo* assembly principles (Jin et al. 2020). NOVOPlasty was executed with default parameters, including “Genome Range 120000-200000” and “K-mer 31”, to assemble plastomes for all *Pyrus* samples. The *rbc*L sequence (accession: KP088778) and the chloroplast genome (accession: KX450880) of *Pyrus × bretschneideri* from GenBank served as the seed and the reference for sequence extension, respectively. For GetOrganelle, default settings, including Maximum extension rounds (-R) set to 15 and ORGANELLE_TYPE (-F) set to embplant_pt, were used, except for adjustments to the default SPAdes kmer settings to 21, 45, 65, 85, 105. After integrating assembly results from both approaches, a subset of samples remained unassembled into circular plastomes, primarily due to sequencing gaps in read coverage. For these specific cases, we adopted an alternative method developed by our PhyloAI team, known as the Successive Approach, combining Reference-based and De novo assembly (SARD; Liu et al. 2023). The SARD approach has proven effective across various angiosperm lineages, particularly in the plastome assembly of low-quality raw reads. Briefly, Bowtie2 (Langmead and Salzberg 2012) was initially utilized to align the reads of these unsuccessfully assembled samples to the plastomes mentioned in the preceding NOVOPlasty plastome assembly. Subsequently, a consensus sequence for each sample was generated using Geneious Prime v. 2023.0.1 (Kearse et al. 2012). The scaffolds and contigs of the corresponding sample retrieved from NOVOPlasty and GetOrganelle were then realigned to their draft plastome to rectify errors and ambiguities, resulting in high-quality complete plastomes. The assembled plastomes were annotated using the Plastid Genome Annotator (PGA) (Qu et al. 2019). All assembled chloroplast genomes were visualized in Geneious Prime (Kearse et al. 2012) to confirm start and stop codons of each coding gene, with manual correction of any discrepancies in the annotations. All assembled plastomes in this study were submitted to the GenBank database, with their respective accessions listed in supplementary table S1.

### Single-copy nuclear marker development and sequence assembly

To retreive SCN genes for all samples in this investigation, we employed a custom-designed nuclear SCN marker set encompassing 801 SCN genes, with the reference genes available on Dryad [DOI: 10.5061/dryad.hx3ffbghm]. This particular SCN gene set has been validated across all lineages within the apple tribe Maleae. The development of these 801 nuclear SCN markers was executed utilizing MarkerMiner v. 1.0 (Chamala et al. 2015), based on the genomic sequences of *Malus domestica* (GenBank accession no. GCF_002114115.1), *Pyrus ussuriensis* × *P. communis* (GenBank accession no. GCA_008932095.1), and *Prunus persica* (GenBank accession no. GCA_000346465.2). Comprehensive details regarding the marker design methodology and the parameters employed are described in our previous publication (Jin et al. 2023).

The assembly of nuclear loci was conducted using HybPiper v. 2.0.1 (Johnson et al. 2016), an integrative software suite that wraps various bioinformatics tools. We extracted lineage-specific SCN sequences from NGS reads using default parameters in the HybPiper pipeline. Briefly, we first used the ‘hybpiper assemble’ command to facilitate the mapping and sorting of trimmed reads against the aforementioned SCN gene reference sequences using BWA v. 0.7.17 (Li and Durbin 2009) and SAMtools v. 1.17 (Li et al. 2009). The sorted reads were subsequently *de novo* assembled into contigs or supercontigs using SPAdes v. 3.15.5 (Bankevich et al. 2012), applying a coverage cutoff value of 5. Further analysis involved the use of ‘hybpiper stats’ and ‘hybpiper recovery_heatmap’ commands for the quantitative summarization and visualization of gene recovery efficiency across different species.

Additionally, the ‘hybpiper paralog_retriever’ command was employed for the paralog detection and retrieval of gene sequences with flagged paralogs, a critical step considering the potential influence of chimeric sequences on subsequent orthology inference. Consequently, any sequences deemed putatively chimeric were excluded.

### Orthology inference for the nuclear genes

To accurately estimate evolutionary history through orthologs, we utilized the tree-based orthology inference method for SCN genes proposed by Yang and Smith (2014), generating two ortholog datasets, MO and RT. The specific procedures for this method are detailed in Morales-Briones et al. (2022). Additionally, uneven sequencing coverage in this study resulted in some orthologs exhibiting outlier loci and short sequences, which could potentially impact phylogenetic inference. To address these issues, our PhyloAI team innovatively developed a comprehensive pipeline to refine these ortholog sequences. This pipeline has been successfully applied in several studies, including those by Liu et al. (2021, 2022) and Jin et al. (2023, 2024). For detailed descriptions of these procedures, refer to the Supplementary Methods.

### Multiple inference methods for phylogenetic analyses

This study employed multiple phylogenetic inference approaches to accurately estimate the plastid/nuclear phylogeny of the genus *Pyrus*, using both concatenated and coalescent-based inference methods. We used two Maximum Likelihood (ML) programs for phylogenetic inference, designed explicitly for the concatenated supermatrix derived from nuclear and/or plastid datasets. All orthologs in each dataset were combined into a supermatrix with AMAS v. 1.0 (Borowiec 2016). Next, the optimal partitioning scheme and evolutionary models for each partition were determined using PartitionFinder2 (Stamatakis 2006; Lanfear et al. 2017). This process was guided by the Corrected Akaike Information Criterion (AICc) and the rcluster algorithm (Lanfear et al. 2014). The branch lengths were uniformly set to be linked across the analyses. The estimated optimal partitioning scheme and evolutionary models for each SCN gene were then used to conduct ML inference. This was executed using IQ-TREE2 v. 2.1.3 (Minh et al. 2020), which included 1000 replicates for the SH approximate likelihood ratio test and ultrafast bootstrap, and RAxML v. 8.2.12 (Stamatakis 2014), utilizing the GTRGAMMA model along with 200 bootstrap replicates.

We verified the boundaries of the two inverted repeats in each plastome utilizing the Repeat Finder plugin within the Geneious Prime software suite (Kearse et al. 2012).

Subsequently, one of the duplicated repeats was removed from further analysis. The well-annotated 77 plastid CDSs were then extracted from these processed plastomes and then aligned by MAFFT v. 7.475 with “--auto” option. Following sequence alignment, all generated alignments were concatenated using AMAS v. 1.0 (Borowiec 2016). This concatenated dataset served as the input for PartitionFinder2 (Stamatakis 2006; Lanfear et al. 2017), which was employed to identify the best-fit partitioning schemes and nucleotide substitution models. It is important to note that the parameter settings used here were consistent with those applied in the nuclear phylogenetic inference, with the exception of employing the greedy algorithm (Lanfear et al. 2012) for this specific analysis. The resultant supermatrix and the identified partitioning scheme were then utilized to conduct ML phylogenetic inference. This inference was carried out using two software tools: IQ-TREE2 v. 2.1.3 (Minh et al. 2020) and RAxML v. 8.2.12 (Stamatakis 2014), and the parameters follow the settings in the nuclear phylogeny. The aligned plastid supermatrix was deposited in the Dryad Digital Repository https://doi.org/10.5061/dryad.3ffbg79r8.

A coalescent-based approach was employed for species tree estimation using the ASTRAL-III software (Zhang et al. 2018), applied to both nuclear and plastid CDSs datasets. For each gene within the nuclear and plastid datasets, individual gene trees were estimated using RAxML v. 8.2.12 (Stamatakis 2014), with the GTRGAMMA model and 100 fast bootstrap (BS) replicates. To enhance the reliability of these gene trees, branches exhibiting support values below 10 were collapsed using the ‘pxcolt’ command in *phyx* toolkit (Brown et al. 2017). Subsequently, the refined gene trees, having undergone the process of branch collapse, were integrated and summarized into a species tree with ASTRAL-III with default parameters. The study also ensures accessibility and transparency of its findings by making all nine resultant trees available through the Dryad Digital Repository: https://doi.org/10.5061/dryad.3ffbg79r8, thus providing an opportunity for further scrutiny and analysis by the scientific community.

### Detecting and visualizing nuclear gene tree discordance

We employed various approaches to evaluate congruence among gene trees and the inferred phylogeny. First, we utilized the program phyparts (Smith et al. 2015) to examine phylogenetic conflict. This approach maps individual gene trees to the target tree and quantifies the number of discordant and congruent bipartitions. The gene trees and the target tree were both rooted using the ‘pxrr’ command in *phyx* (Brown et al. 2017), followed by a full concordance analysis (-a 1) using phyparts. Notably, nodes with bootstrap support (BS) less than 50% in each gene tree were considered uninformative and excluded from analysis. Additionally, to mitigate the impact of missing taxa in some loci due to uneven recovery efficiency of nuclear genes, a rapid concordant analysis (-a 0) was conducted. The results were amalgamated with the previous full concordance analysis to generate the final outcomes, visualized using a Python script (available at https://bitbucket.org/dfmoralesb/target_enrichment_orthology/src/master/phypartspiecharts_m issing_uninformative.py). The proportion of discordant and congruent topologies for each node was presented as a pie chart. Furthermore, we also calculated the internode certainty all (ICA) value to summarize the degree of inconsistency on each node in our dataset using phyparts.

Additionally, Quartet Sampling (QS, Pease et al. 2018) was employed to distinguish conflicting support from nodes with weak support. By subsampling the target tree and the combined supermatrix of all the genes, QS assesses the reliability of internal tree relationships and each terminal branch rapidly, producing several values to reflect the degree of conflict.

We conducted the QS analysis using 100 replicates with a log-likelihood threshold set to 2 and visualized the results with an R script (available at https://github.com/ShuiyinLIU/QS_visualization).

### SNP calling and gene flow analyses

The latest, high-quality genome assembly of *Pyrus pyrifolia* (accession number: GCA_016587475.1) was downloaded as a reference genome for single nucleotide polymorphism (SNP) calling. Clean reads of each sample were mapped to this reference genome using BWA v. 0.7.17 (Li and Durbin 2009). Subsequently, SAMtools v. 1.6 (Li et al. 2009) was employed to convert and sort the aligned results into bam files. Duplicate reads were marked, and variants were identified using the ‘MarkDuplicates’ and ‘HaplotypeCaller’ functions in the GATK v. 4.3.0.0 (McKenna et al. 2010), respectively. Criteria for base calling included a minimum base quality threshold of 30, while variant sites were retained only if they surpassed a quality score of 30. Haplotypes for each sample were combined using the ’‘CombineGVCFs’ and ‘GenotypeGVCFs’ functions in GATK to produce genotype files (gVCF). We performed two-step filtering for the quality guarantee. The VCF file underwent initial filtering using the ‘VariantFiltration’ function in GATK, applying parameters: QD < 2.0, FS > 60.0, MQ < 40.0, MQRankSum < -12.5, and ReadPosRankSum < −8.0. All SNPs were then extracted using the ‘SelectVariants’ function in GATK. A secondary filtration was performed using VCFTOOLS (Danecek et al. 2011), following the parameters of minQ 30, max-missing 0.67, max-alleles 2, max-meanDP 500, and min-meanDP 10, to obtain the final variant sites for downstream gene flow analysis.

The f4-ratio was calculated using *D*suite v. 0.5 (Malinsky et al. 2021) to investigate potential gene flow between species. This analysis utilized SNPs as the input data, and an employed as the guiding tree. Visualization of the f4-ratio statistics results were accomplished using the ‘plot_f4ratio.rb’ Ruby script, which is publicly available at https://github.com/mmatschiner/tutorials/tree/master/analysis_of_introgression_with_snp_data. We executed a series of parallel analyses to evaluate the potential influence of outgroup selection on our analytical outcomes. In these, we alternated the outgroup, employing either a representative individual from the *Malus* genus or one from the *Sorbus* genus. The comparative analysis revealed high consistency in the results, irrespective of the outgroup used.

### Incomplete lineage sorting analyses

To thoroughly investigate the role of ILS in shaping the evolutionary trajectory of *Pyrus*, this study employed two alternative approaches, integrating both the population mutation parameter theta (Cai et al. 2021) and coalescent simulation analysis (Liu et al. 2022). In the first approach, we analyzed the population mutation parameter theta (Cai et al. 2021) at each nodal point. Theta was calculated by dividing the branch length in mutation units, inferred by IQTREE, by the length in coalescent units as estimated through ASTRAL-III. The determination of branch lengths in mutation units utilized the ASTRAL-III tree as a fixed topology, a methodological choice designed to ensure consistent topology between the compared trees. Furthermore, additional analysis of the correlation between the branch lengths and the ICA values of the ASTRAL-III tree were conducted to assess the impact of ILS. A strong positive correlation between branch lengths and ICA values can be interpreted as ILS being responsible for tree conflicts (Zhou et al. 2022).

To elucidate the evolutionary processes underlying the observed phylogenetic discrepancies between gene trees and the species tree in *Pyrus*, the present study adopted an alternative approach, i.e., coalescent simulation analysis approach (Liu et al. 2022), aimed at assessing the contribution of ILS in resolving these phylogenetic conflicts. A dataset of 29 high-quality samples was subsampled based on nuclear gene recovery. Subtrees for these species were extracted from all gene trees, and a species tree was then inferred using ASTRAL-III. Utilizing Phybase v. 1.5 (Liu and Yu 2010), 10,000 gene trees were simulated under the multi-species coalescent model with the species tree as the input. Distances among simulated gene trees, empirical gene trees, and the species tree were quantified using DendroPy v. 4.5.2 (Sukumaran and Holder 2010), and then visually compared. The disparity in distance distributions between simulated and empirical gene trees was analyzed to assess the effect of ILS on gene tree incongruence.

### Polyploidy analyses

We integrated multiple sources of evidence to investigate the possible impact of polyploidy or whole-genome duplication (WGD) on phylogenetic discrepancies in *Pyrus*. In the first step, a comprehensive literature review was undertaken, and we collected all chromosome-related data, including the ploidy level of all *Pyrus* species. This information was extracted from existing scholarly publications and digital resources, notably the International Plant Chromosome Number (IPCN) database (http://legacy.tropicos.org/Project/IPCN). In addition, we employed Smudgeplot (Ranallo-Benavidez et al. 2020) to infer the ploidy of individual samples by analyzing the k-mers within sequencing reads. All these two methods can thoroughly identify the potential polyploid species in *Pyrus*.

To further explore the incidence of polyploidization and WGD events within the deep phylogenetic history of *Pyrus*, a sophisticated analysis, initially conceptualized by Morales-Briones et al. (2022), was conducted. This methodology involved extracting rooted ortholog trees from homolog trees. Adhering to a stringent filtering criterion that required an average bootstrap value of no less than 50% per ortholog tree, gene duplications identified within these rooted ortholog trees were then mapped onto a rooted species tree. After this mapping, the proportions of gene duplication at the nodes correlating with the most recent common ancestors (MRCAs) were meticulously documented. The computational scripts implemented for this analysis are publicly available and can be accessed at https://bitbucket.org/blackrim/clustering.

### Inference of global split networks

SplitsTree is an ideal tool for global split network computation, especially for deriving unrooted phylogenetic networks from molecular sequence data. This utility is achieved through various sophisticated methods, including split decomposition, neighbor-net, consensus network, and super networks techniques. Specifically focusing on the *Pyrus* case study, we employed SplitsTree v. 4.19.0 (Huson and Bryant 2006) to investigate and clarify the complex, web-like evolutionary trajectory of the *Pyrus* genus. For this analysis, our primary dataset comprised well-aligned SCN genes derived from the MO dataset, and this dataset included a diverse representation of 50 *Pyrus* samples, alongside an individual from *Malus* as the outgroup. This investigation entailed an in-depth inference of the implicit network, employing parameters such as uncorrected_P distances, the EqualAngle network construction algorithm, and the NeighborNet method.

### Phylogenetic network analyses

To advance our understanding of the complex reticulate evolutionary processes in the *Pyrus* genus, we used an explicit network analysis utilizing the Species Networks applying Quartets (SNaQ) algorithm. This software, developed in the Julia programming language and integrated within the PhyloNetworks (Solís-Lemus et al. 2017), employs maximum pseudolikelihood methods for inferring phylogenetic networks from multi-locus datasets.

SNaQ offers increased efficiency and tractability, particularly when scaling up the number of taxa or hybridization events, compared to full likelihood-based methodologies. In this analysis, gene flow and ILS were considered potential sources of discordance among gene trees, and these factors were duly accounted for in the species network estimation process. To reduce the computational burden, three distinct datasets were generated to investigate potential reticulate evolutionary events within the origin and diversification of *Pyrus*. Each dataset was constrained to fewer than 20 samples to maintain computational feasibility. The first dataset, termed “Maleae 15-taxa data,” aimed to test the possible hybridization origin of *Pyrus*, encompassing 15 representative species from genera closely related to *Pyrus*, such as *Malus* and *Sorbus* s.l. For the “coalescent simulation analysis” part, we selected 29 high-quality samples representing 26 currently recognized *Pyrus* species. This selection encompassed 15 taxa from *P.* subg. *Pyrus* and 14 taxa from *P.* subg. *Pashia*. The second dataset, named “*Pyrus* 16-taxa data,” included 15 taxa from *P.* subg. *Pyrus*, with an additional *Malus* sample serving as the outgroup. Similarly, the third dataset, called “*Pashia* 15-taxa data,” comprised the 14 taxa from *P.* subg. *Pashia*, plus one *Malus* sample as the outgroup.

Notably, the species selected for each dataset were either representative of major lineages or known for their cytonuclear discordance, thus offering a comprehensive view of the genetic diversity and evolutionary dynamics within *Pyrus*.

For each dataset, we utilized all SCN gene trees to summarize quartet concordance factors (CFs) using the ‘readTrees2CF’ package within the PhyloNetworks software. Next, we reconstructed the species tree with ASTRAL-III (Zhang et al. 2018). The derived CFs and the species tree were subsequently employed as input data for inferring the optimal phylogenomic network. The maximum number of reticulation events (*h_max_*) was set from 0 to 6, and the inheritance probabilities, denoted as γ and 1-γ, were calculated and plotted near the hybridization edges. Subsequently, we summarized the pseudo-deviance score of the optimal network of all six runs, followed by their visualization. The optimal network was identified by selecting the *h_max_* value corresponding to a stable score in the analytical metrics, marked by a plateau following an initial rapid decrease, indicating consistency in network scoring.

### Dating analysis and ancestral area reconstruction

In the context of the Maleae tribe, and particularly the genus *Pyrus*, there has been a notable absence of temporal dating analyses, resulting in an imprecise determination of the stem age of *Pyrus*. To address this gap, we adopted a two-step strategy for the divergence time estimation within *Pyrus*. Despite the discovery of various *Pyrus* fossils across different epochs and localities (supplementary table S2), most of these fossils as leaf specimens present a limitation, as leaf morphology alone is insufficient for accurate species identification.

Additionally, the generic classification of some leaf fossils remains ambiguous, posing challenges in distinguishing between *Malus*, *Pyrus*, or other related genera within the Maleae tribe. We first used the MCMCTree, a program implemented in PAML v. 4.9j (Yang 2007), to estimate the divergence times across the Maleae phylogenetic backbone, incorporating two fossil species. The inferred most recent common ancestor (MRCA) age between *Pyrus* and *Malus* was used to set the boundary for the stem age of *Pyrus* in our subsequent analyses. In the following step, we employed BEAST2 (Bouckaert et al. 2014) to refine our estimation of divergence times among *Pyrus* species. This analysis utilized the stem age estimated in the first step of *Pyrus* as a secondary calibration point, thereby enhancing the precision of our temporal divergence estimates within the *Pyrus* genus. The detailed parameter settings for these two software programs can be found in the Supplementary Methods.

The biogeographic analysis of *Pyrus* was conducted using the BioGeoBEARS v. 1.1.1 (Matzke 2018), integrated within RASP v. 4.2 (Yu et al. 2015). The time tree, inferred by analyses. In this study, based on the distribution patterns of the extant *Pyrus* species and the paleotectonic histories of continents, we categorized the geographic areas into three regions: (A) East Asia, (B) Central and West Asia, and (C) Europe and Northern Africa. During the analysis, a constraint was imposed wherein the maximum number of areas assignable to any given phylogenetic node was restricted to two. The selection of the optimal biogeographic model was based on the highest Akaike Information Criterion corrected for small sample sizes (AICc_wt) value, following the comprehensive evaluation of all models available in the BioGeoBEARS toolkit. This methodological approach facilitated a rigorous and data-driven determination of the most plausible biogeographic scenario for the *Pyrus* genus.

### Diversification Analyses

The BAMM (Rabosky 2014) was used to estimate the diversification rate of *Pyrus*. Given the extensive geographical distribution of *Pyrus*, it was impractical to sample all known pear species comprehensively in this study. To address the potential bias from incomplete sampling, we adjusted the default setting to a non-random incomplete taxon sampling strategy. The sampling proportions for the two *Pyrus* clades were specifically determined to mitigate this issue. The configuration of the BAMM was set to operate a species-extinction model, sampling every 1,000 generations. We ran four independent MCMC chains, each for 10,000,000 generations, with the initial 1,000,000 generations discarded as burn-in. We then used the R package BAMMTOOLS (Rabosky et al. 2014) to analyze the BAMM outputs. This analysis helped us calculate and visualize the evolutionary rates over time and identify the 95% credible set of shift configurations.

## Data Availability

All the DGS data are deposited in the NCBI Sequence Read Archive (SRA) under the BioProject PRJNA1031385, and the detailed information for each sample is referred to supplementary table S1. Sequence alignments, phylogenetic trees, and other data files generated in this study have been deposited in the Dryad Digital Repository (https://doi.org/10.5061/dryad.3ffbg79r8). Editors and reviewers can access these files via the following URL: https://datadryad.org/stash/share/wlPQra8K7U7Vj9d_4tvRF3E8fnMxiS3nmDGO0JLcoAE.

## Acknowledgements

The computational analyses in this study were performed on the PhyloAI supercomputer (https://doi.org/10.12282/PhyloAIHPC), under the ownership of Bin-Bin Liu. All the molecular experiments were performed on the Plant DNA and Molecular Identification Platform (PDMIP) of IBCAS. We thank An-Qi Song (Computer Network Information Center, Chinese Academy of Sciences) for her valuable contributions to revising the SSE approach. Financial support for this work was provided by the National Natural Science Foundation of China (grant number: 32270216 & 32000163 to BBL and 32170381 to LZ) and the Youth Innovation Promotion Association CAS (grant number: 2023086 to BBL).

## Author Contributions

B.B.L. and J.W. conceived and designed the study. B.B.L. led and supervised the project. Z.T.J., X.H.L., and D.K.M. wrote the draft manuscript. Z.T.J. carried out the phylogenomic analyses. C.X. performed the deep genome skimming sequencing. R.G.J.H., L.Z., C.R., and L.D. provided suggestions for structuring the paper. All the authors contributed to the writing and interpreting of the results and approved the final manuscript.

## Notes

### Competing Interest Statement

The authors have declared no competing interest.

