## Supplementary material for "Unraveling the Web of Life: Incomplete lineage sorting and hybridization as primary mechanisms over polyploidization in the evolutionary dynamics of pear species": sup_figs.pdf

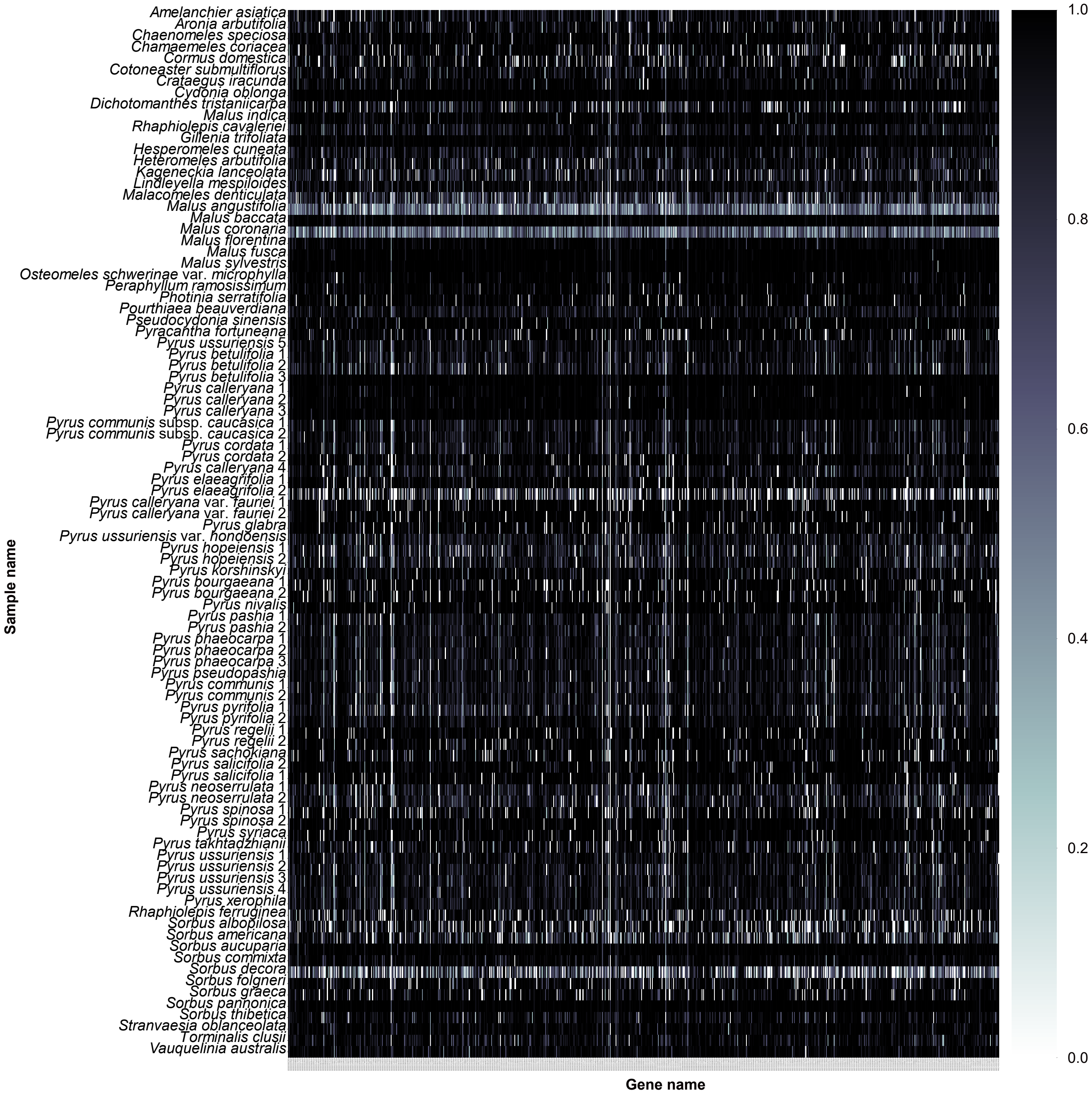


Supplementary Fig. S1. Heat map showing percentage length recovery for SCNs genes recovered by HybPiper. Each row shows a sample, and each column is a gene. The amount of shading in each box corresponds to the length of the gene recovered for that sample by the pipeline, relative to mean reference length (maximum of 1.0).

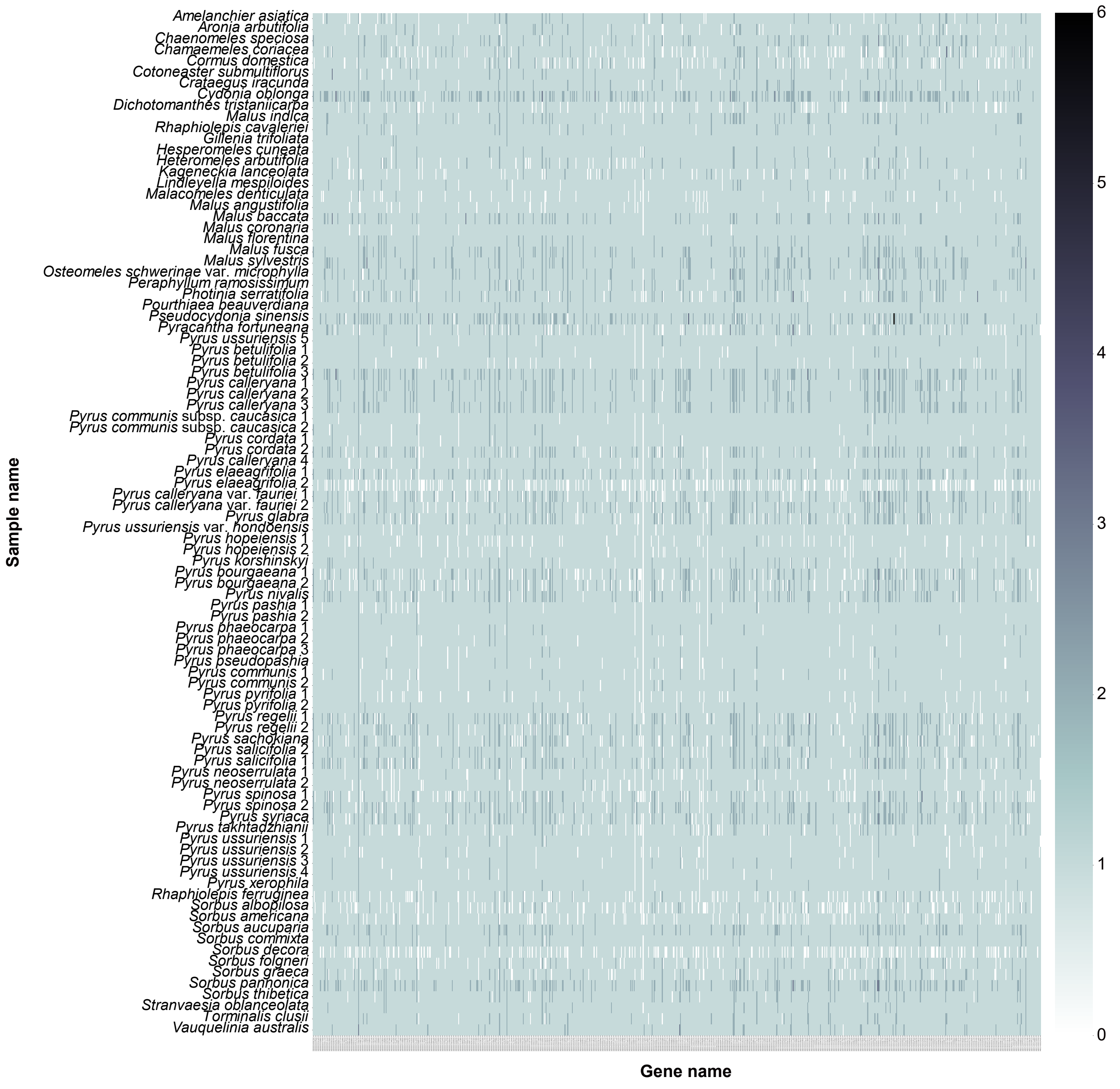


Supplementary Fig. S2. Heat map showing number of paralog sequences for each gene and each sample recovered by HybPiper. Each row shows a sample, and each column is a gene. The amount of shading in each box corresponds to the number of the gene recovered for that sample by the pipeline.

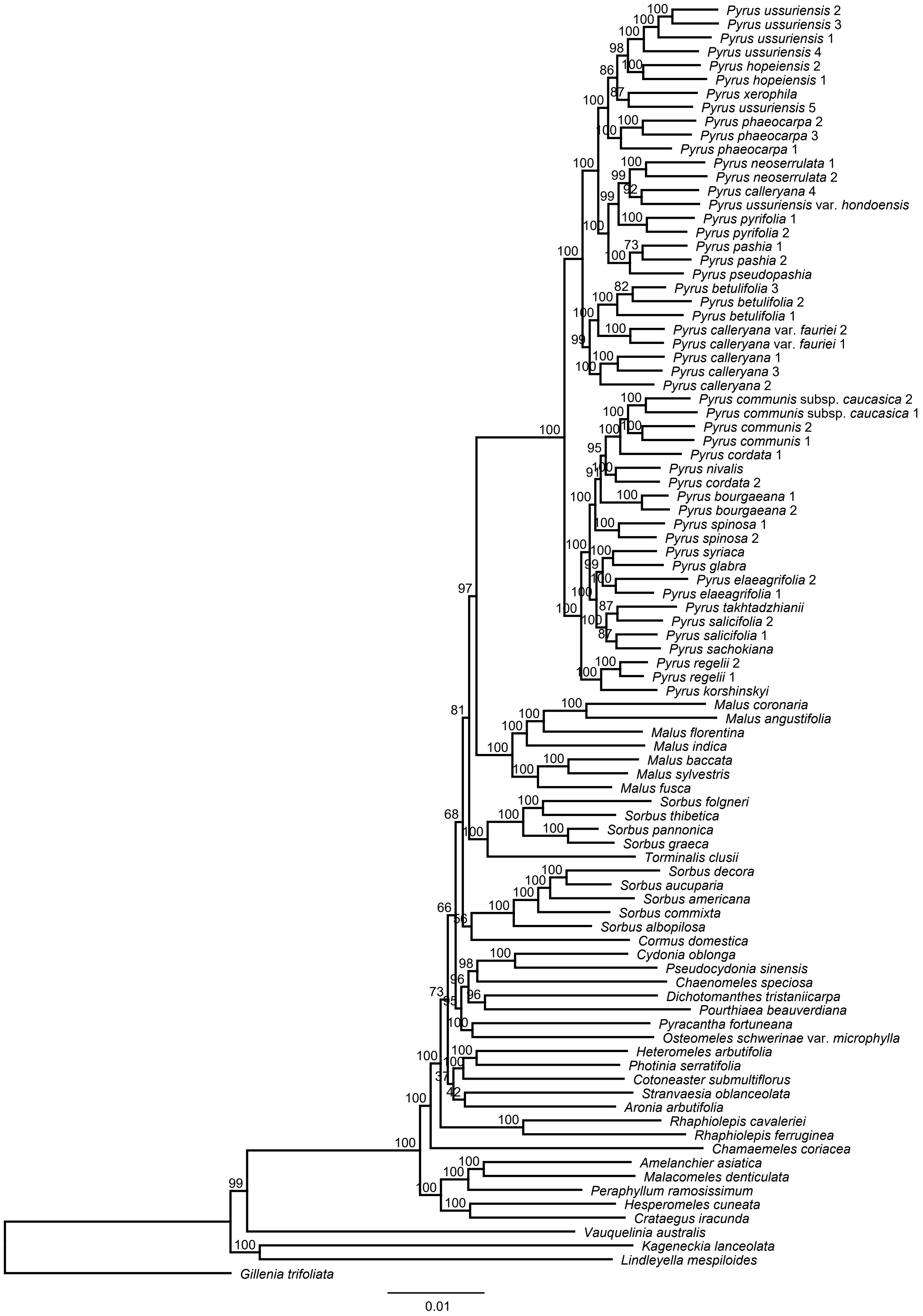


Supplementary Fig. S3. Maximum likelihood phylogeny of *Pyrus* in the framework of Maleae inferred from RAxML analysis of 771 MO orthologs. Bootstrap support (BS) is shown above branches.

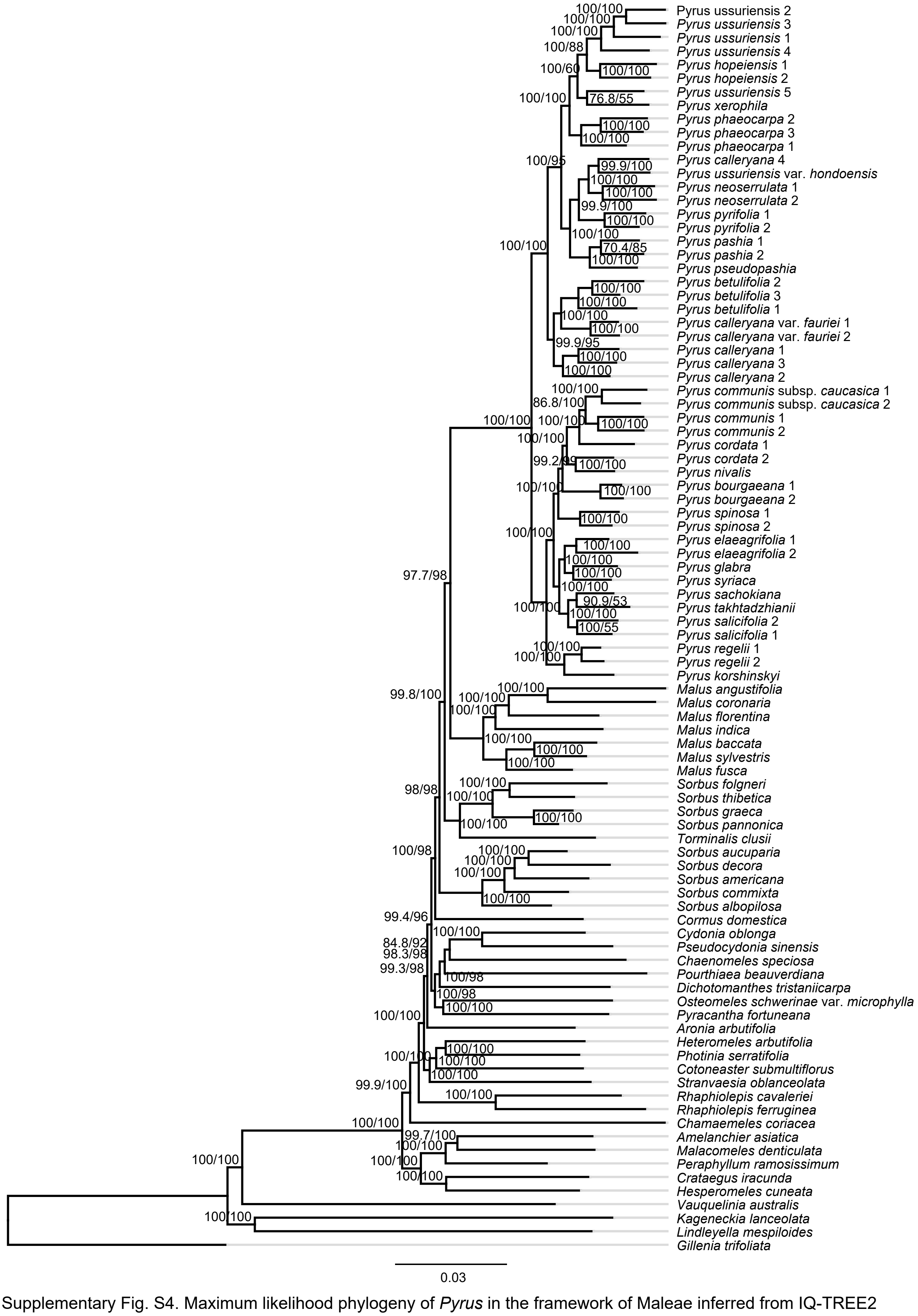


Supplementary Fig. S4. Maximum likelihood phylogeny of *Pyrus* in the framework of Maleae inferred from IQ-TREE2 analysis of 771 MO orthologs. The SH-aLRT support and Ultrafast Bootstrap support (UFBoot) are shown above branches.

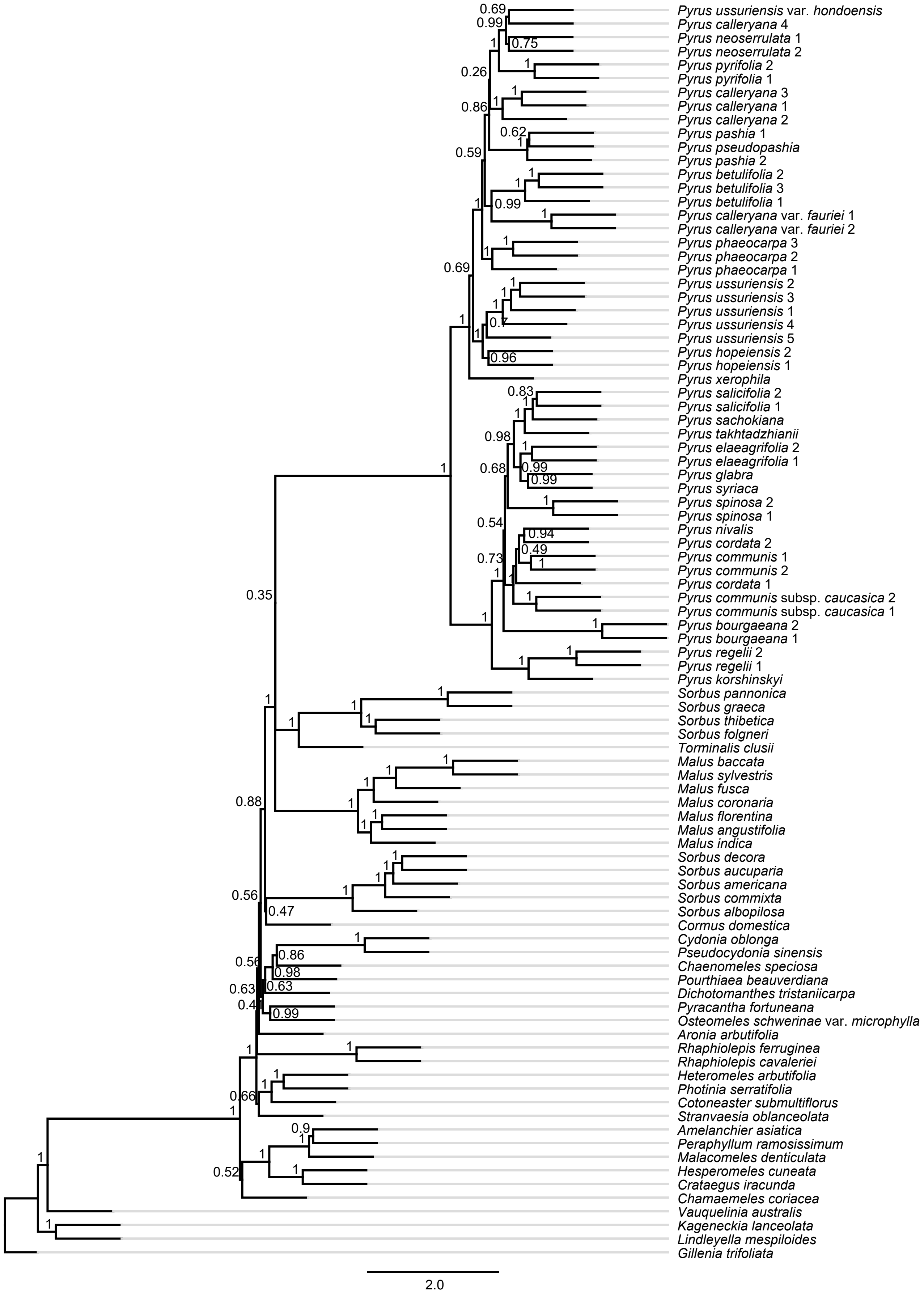


Supplementary Fig. S5. ASTRAL-III Species tree of *Pyrus* in the framework of Maleae inferred from 771 MO orthologs. Local posterior probabilities (LPP) are shown above branches.

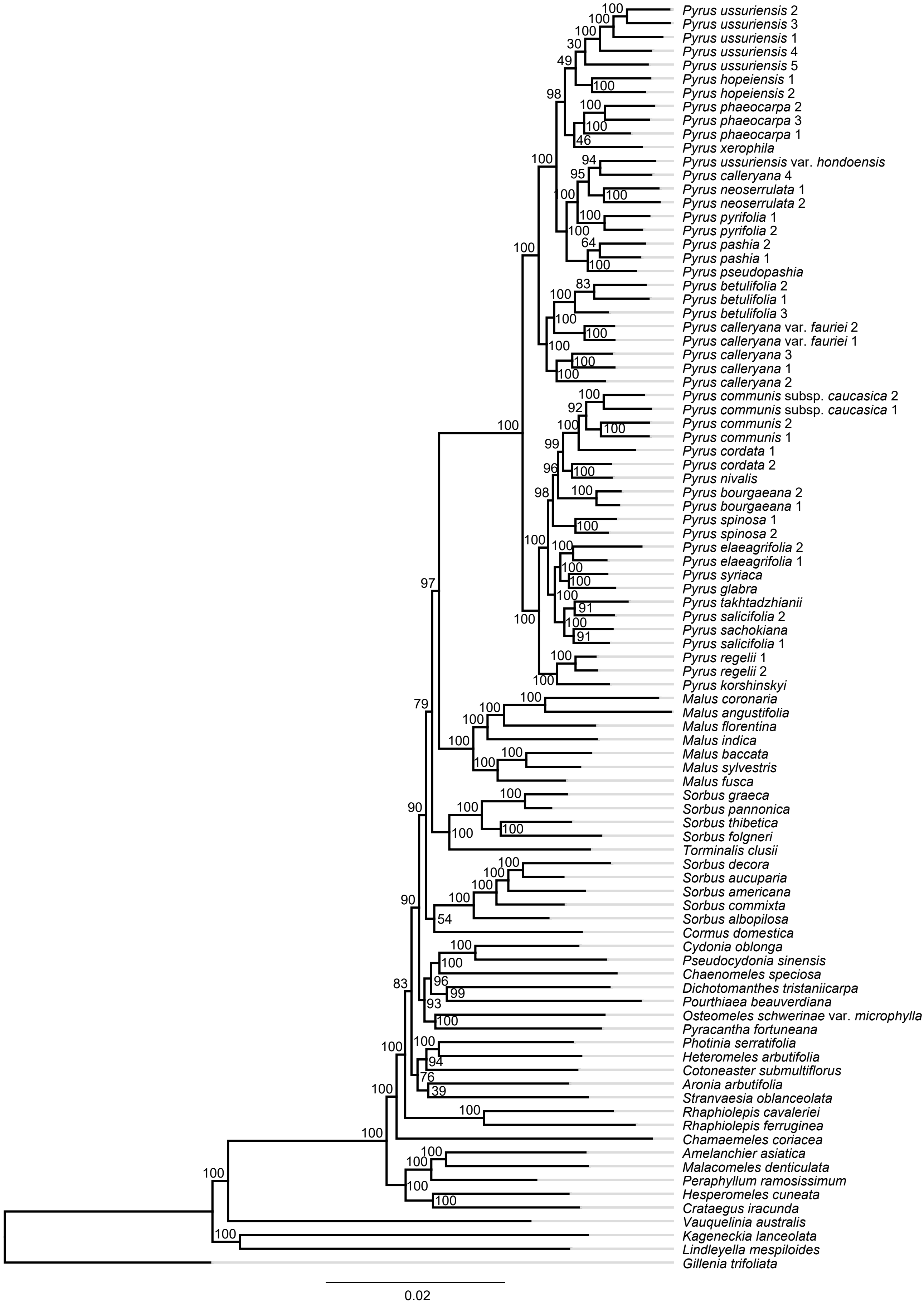


Supplementary Fig. S6. Maximum likelihood phylogeny of *Pyrus* in the framework of Maleae inferred from RAxML analysis of 905 RT orthologs. Bootstrap support (BS) is shown above branches.

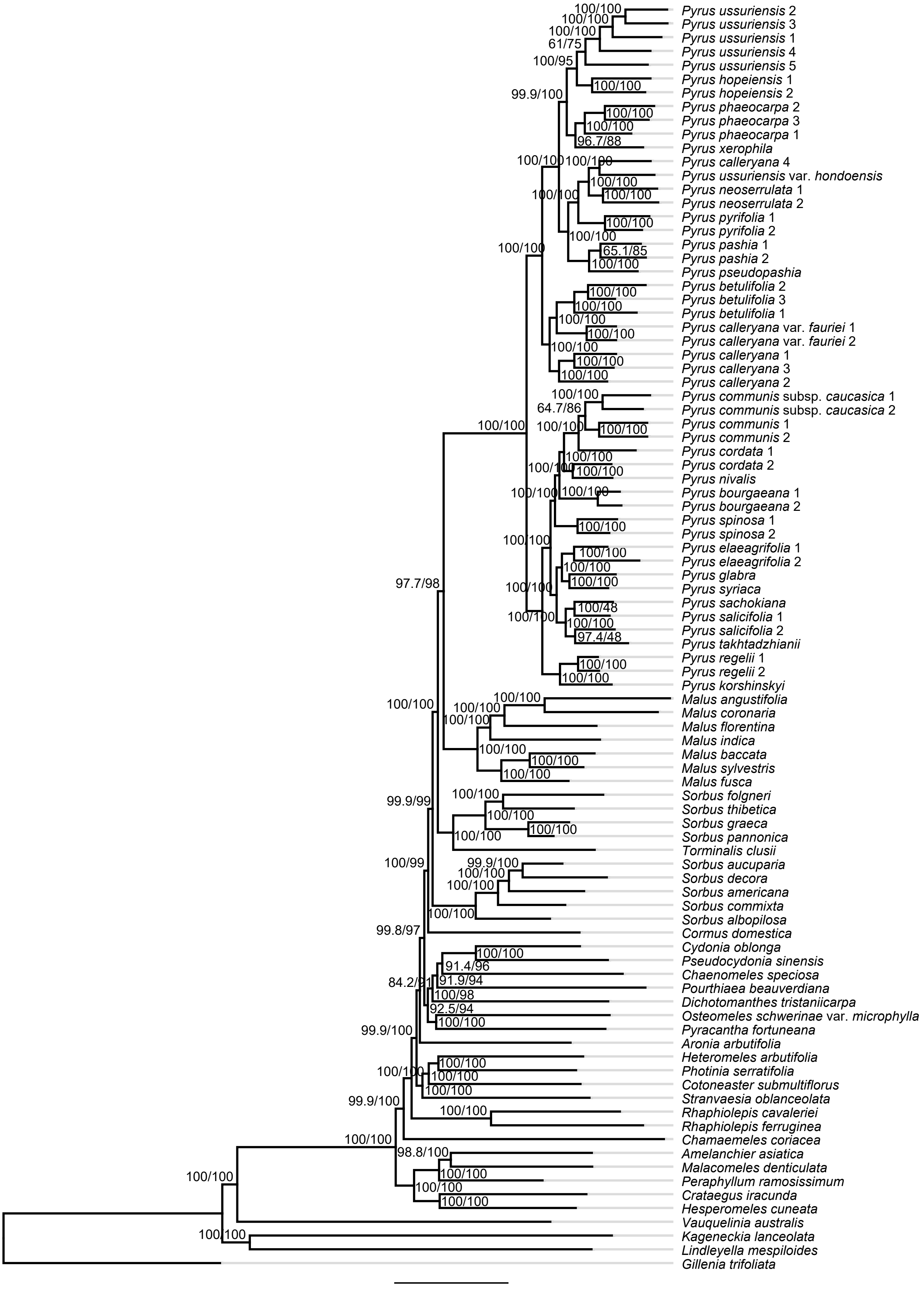


Supplementary Fig. S7. Maximum likelihood phylogeny of *Pyrus* in the framework of Maleae inferred from IQ-TREE2 analysis of 905 RT orthologs. The SH-aLRT support and Ultrafast Bootstrap support (UFBoot) are shown above branches.

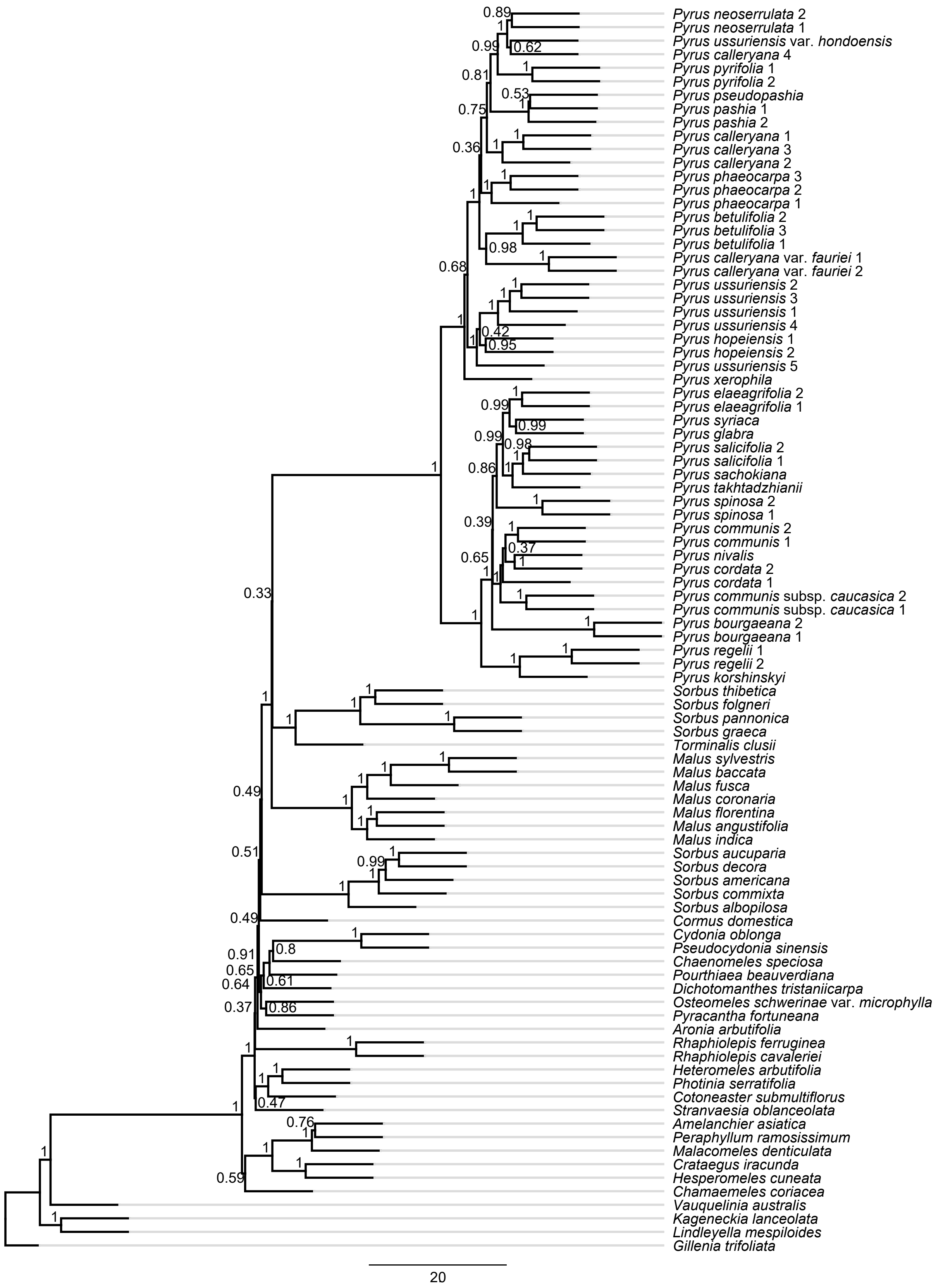


Supplementary Fig. S8. ASTRAL-III Species tree of *Pyrus* in the framework of Maleae inferred from 905 RT orthologs. Local posterior probabilities (LPP) are shown above branches.

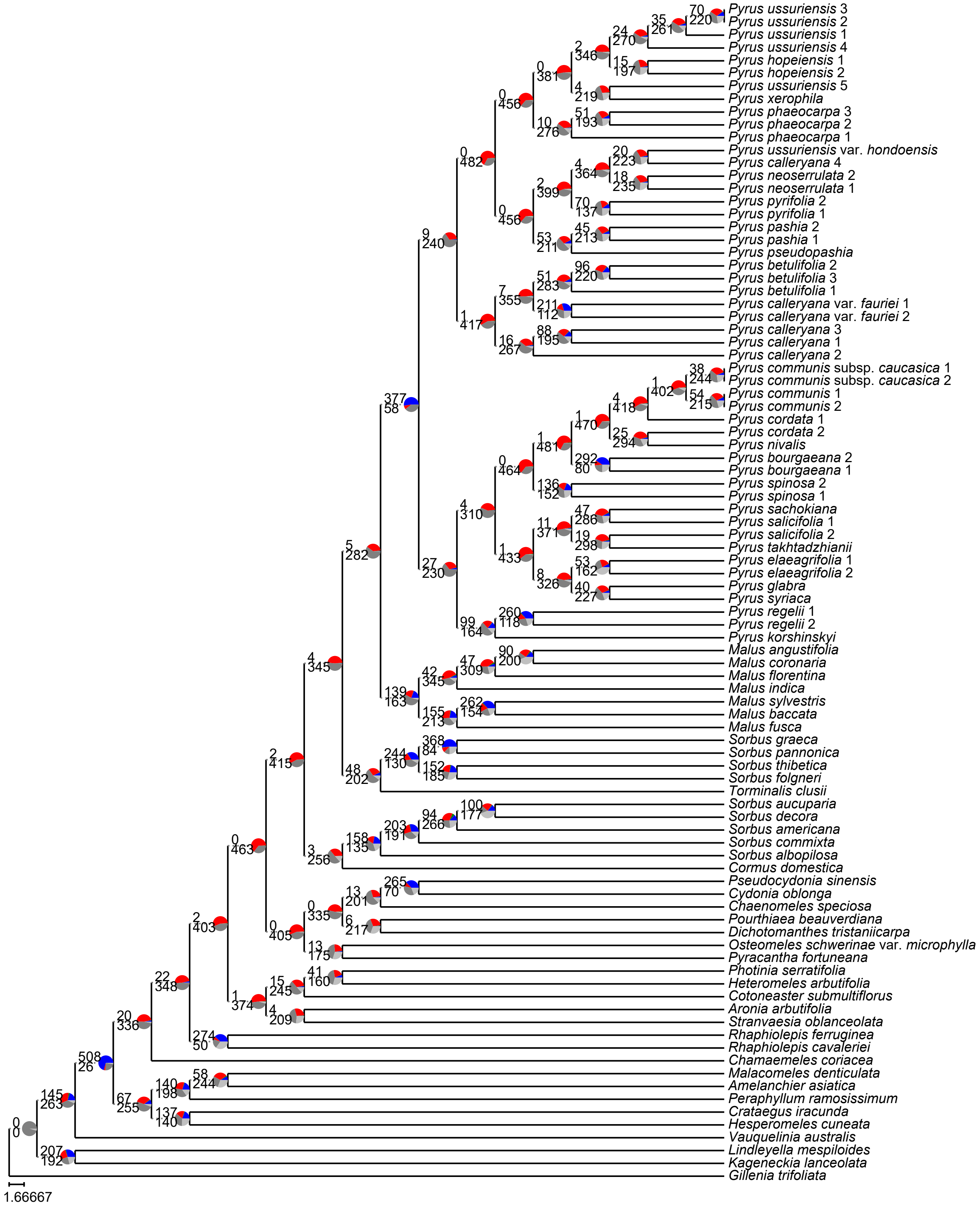


Supplementary Fig. S9. Maximum likelihood phylogeny of *Pyrus* in the framework of Maleae inferred from RAxML analysis of 771 MO orthologs. Pie charts on nodes denote the proportion of gene trees that support that clade (blue), the proportion that support the main alternative bifurcation (green), the proportion that support the remaining alternatives (red), the proportion (conflict or support) that have < 50% bootstrap support (dark grey), and the proportion that have missing taxa (light grey). The number of gene trees concordant with that node in the nuclear phylogeny are shown above branches. The number of gene trees conflicting with that node in the nuclear phylogeny are shown below branches.

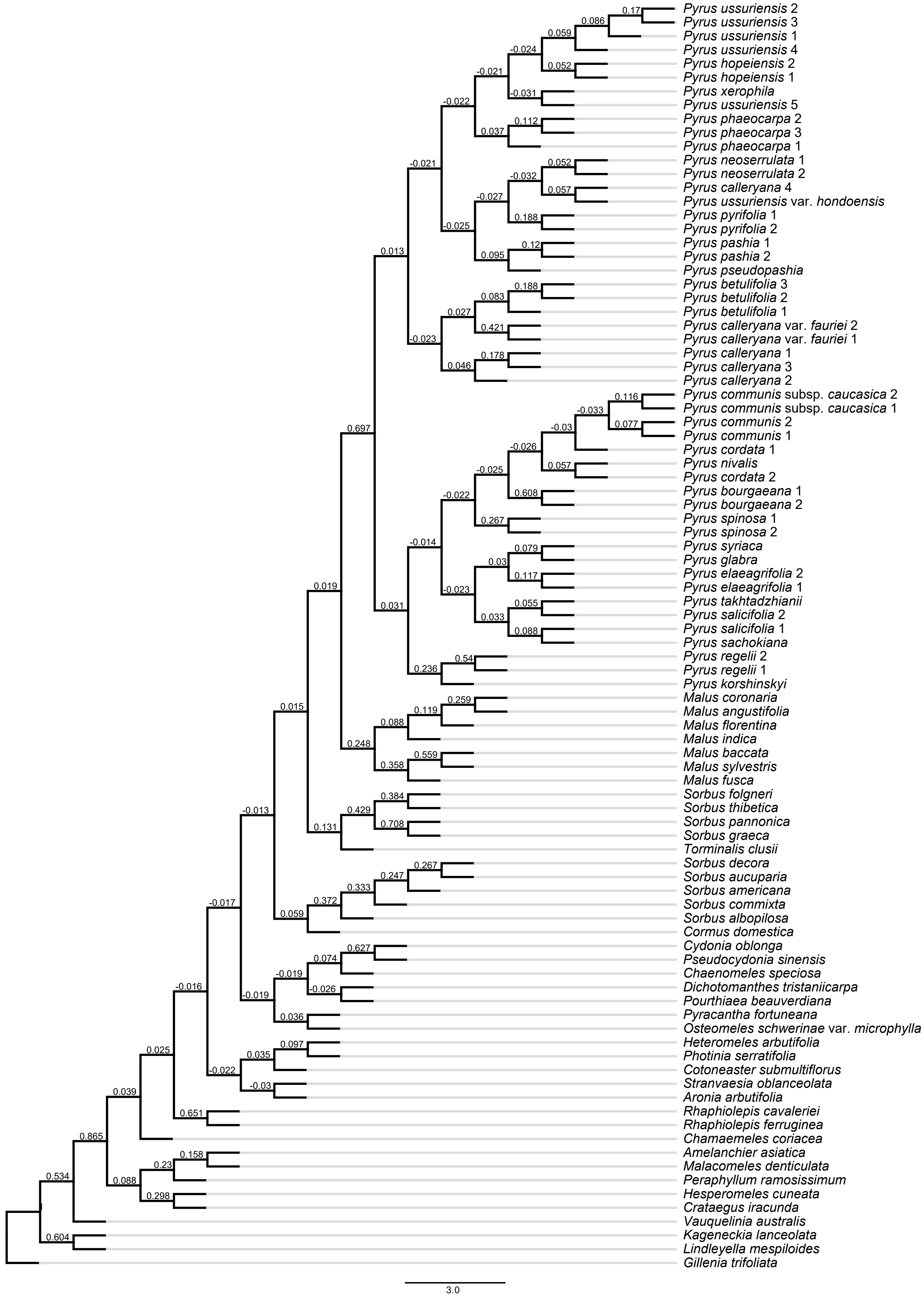


Supplementary Fig. S10. Maximum likelihood phylogeny of *Pyrus* in the framework of Maleae inferred from RAxML analysis of 771 MO orthologs. The Internode Certainty All (ICA) score are shown above branches.

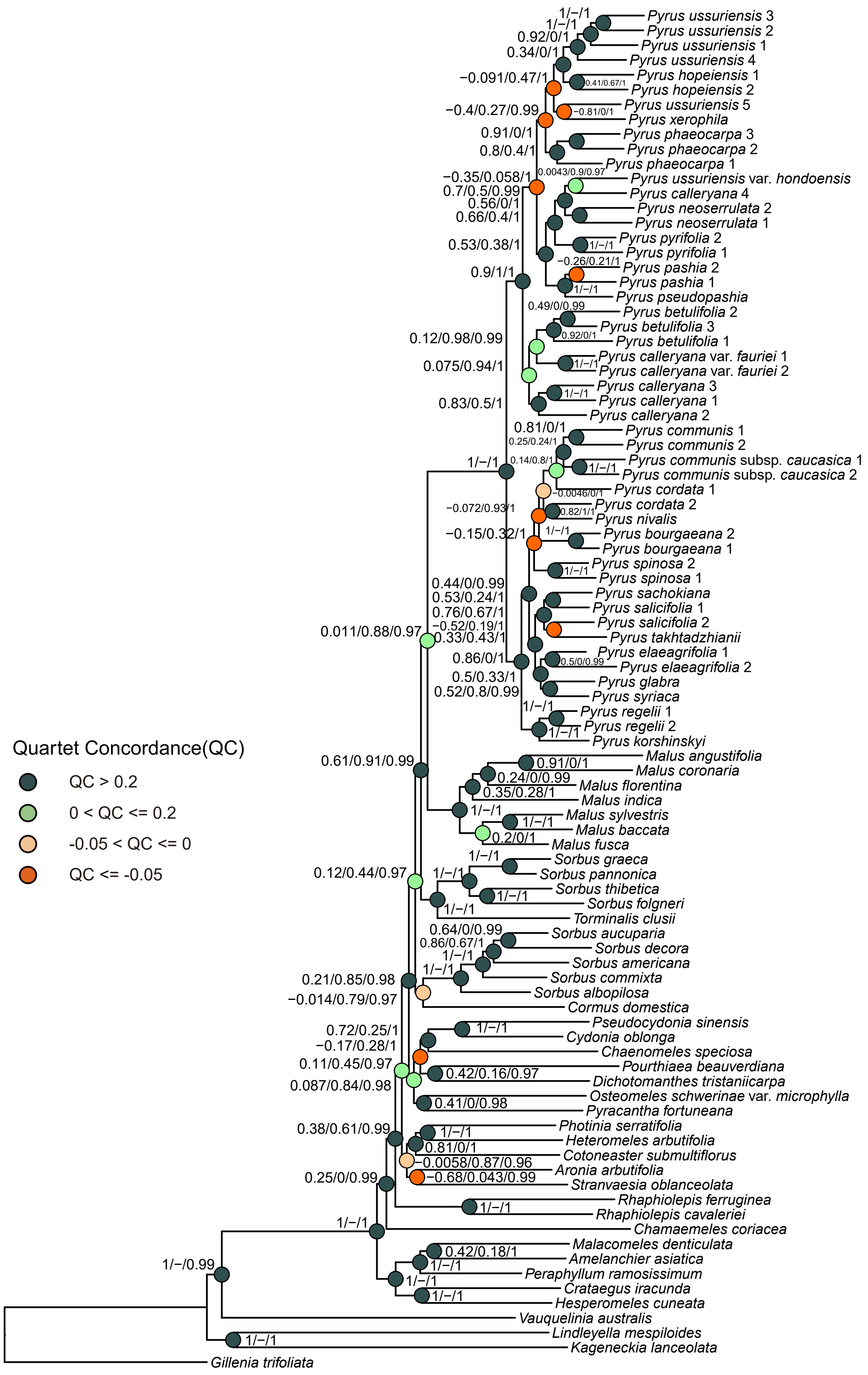


Supplementary Fig. S11. Maximum likelihood phylogeny of *Pyrus* in the framework of Maleae inferred from RAxML analysis of 771 MO orthologs. Quartet Sampling (QS) scores for each node are shown next to branches indicating Quartet Concordance (QC) / Quartet Differential (QD) / Quartet Informativeness (QI). Quartet Concordance is also showed in each node'pie chart and color-coded according to the legend.

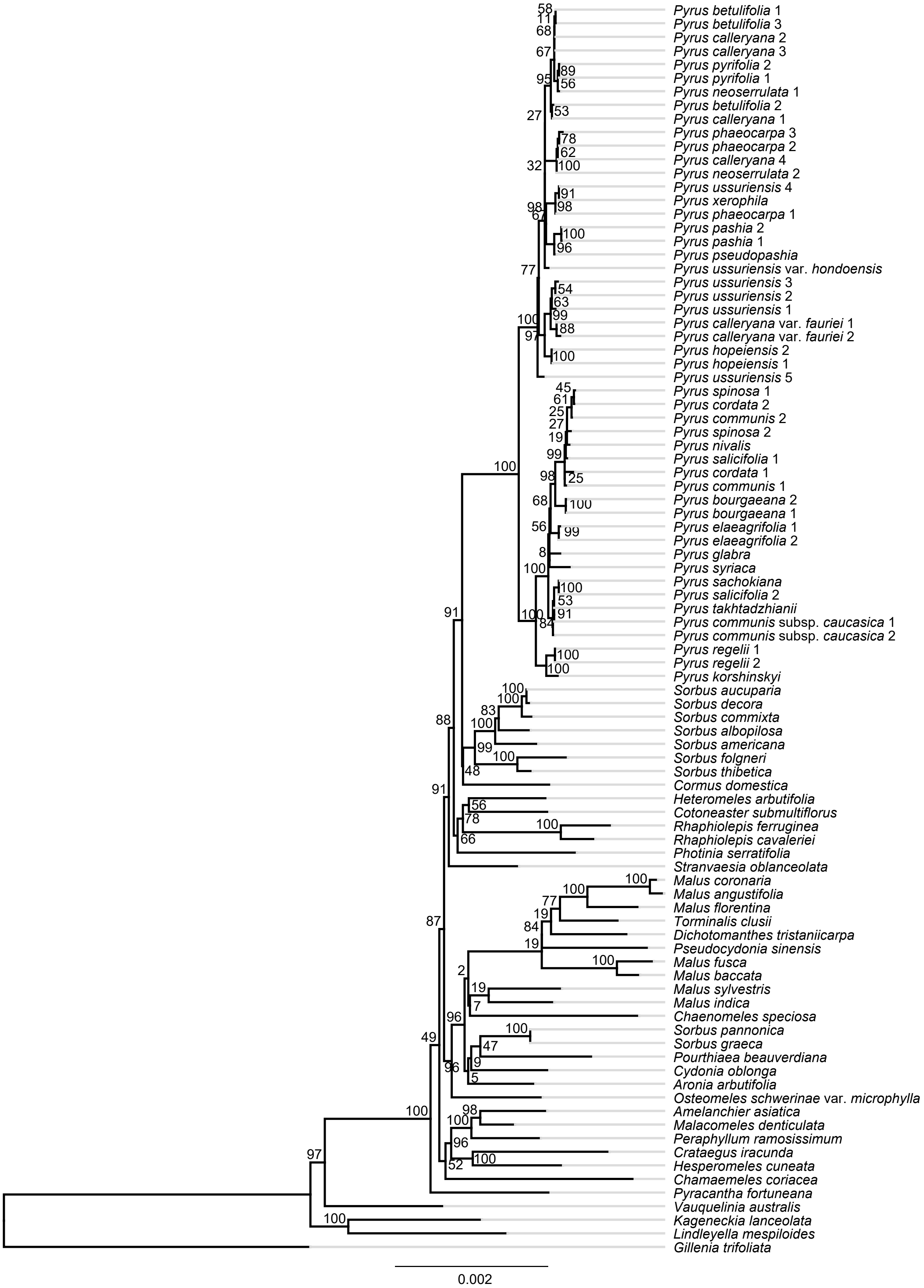


Supplementary Fig. S12. Maximum likelihood phylogeny of *Pyrus* inferred from RAxML analysis of plastid CDS dataset. Bootstrap support (BS) is shown above branches.

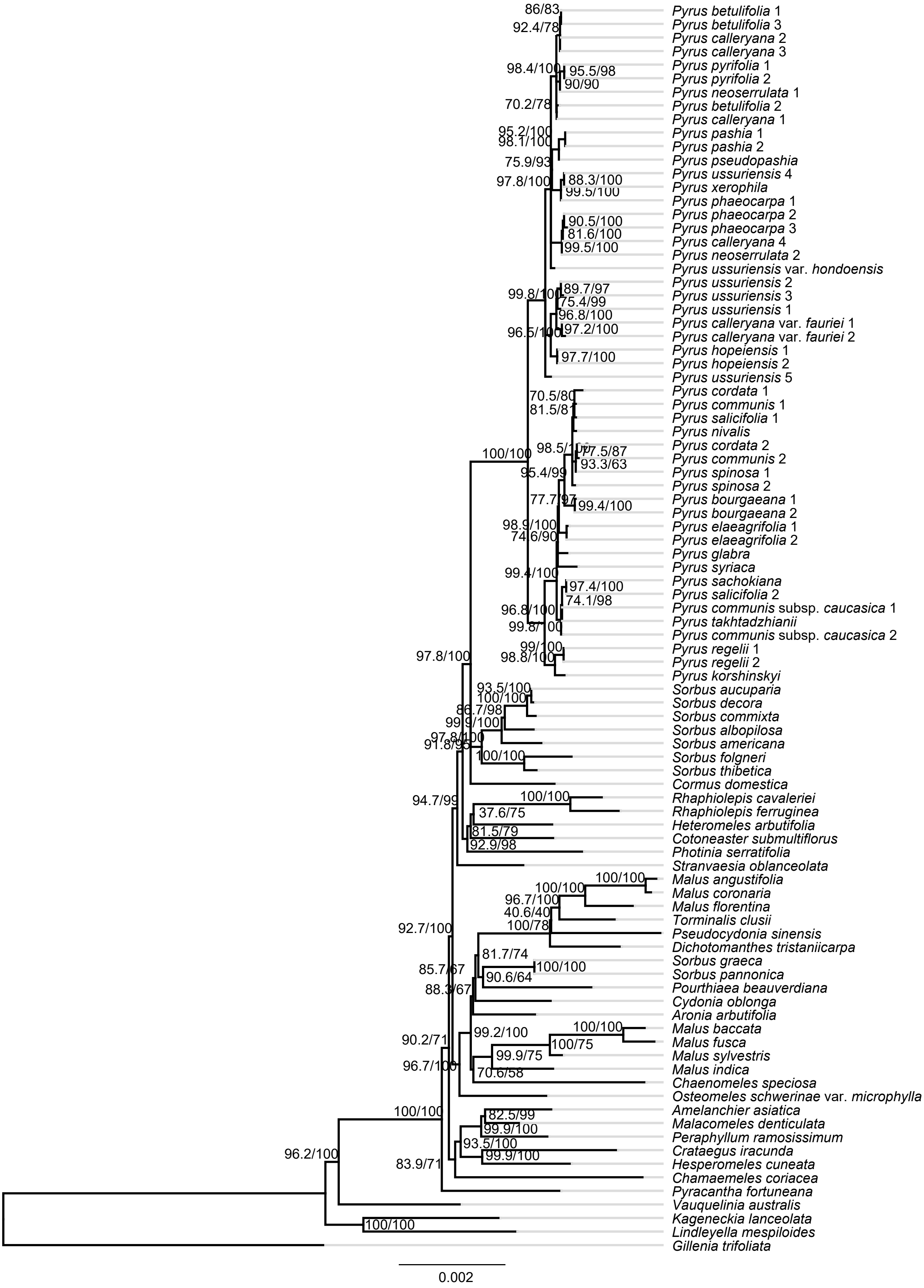


Supplementary Fig. S13. Maximum likelihood phylogeny of *Pyrus* inferred from IQ-TREE2 analysis of plastid CDS dataset. The SH-aLRT support and Ultrafast Bootstrap support (UFBoot) are shown above branches.

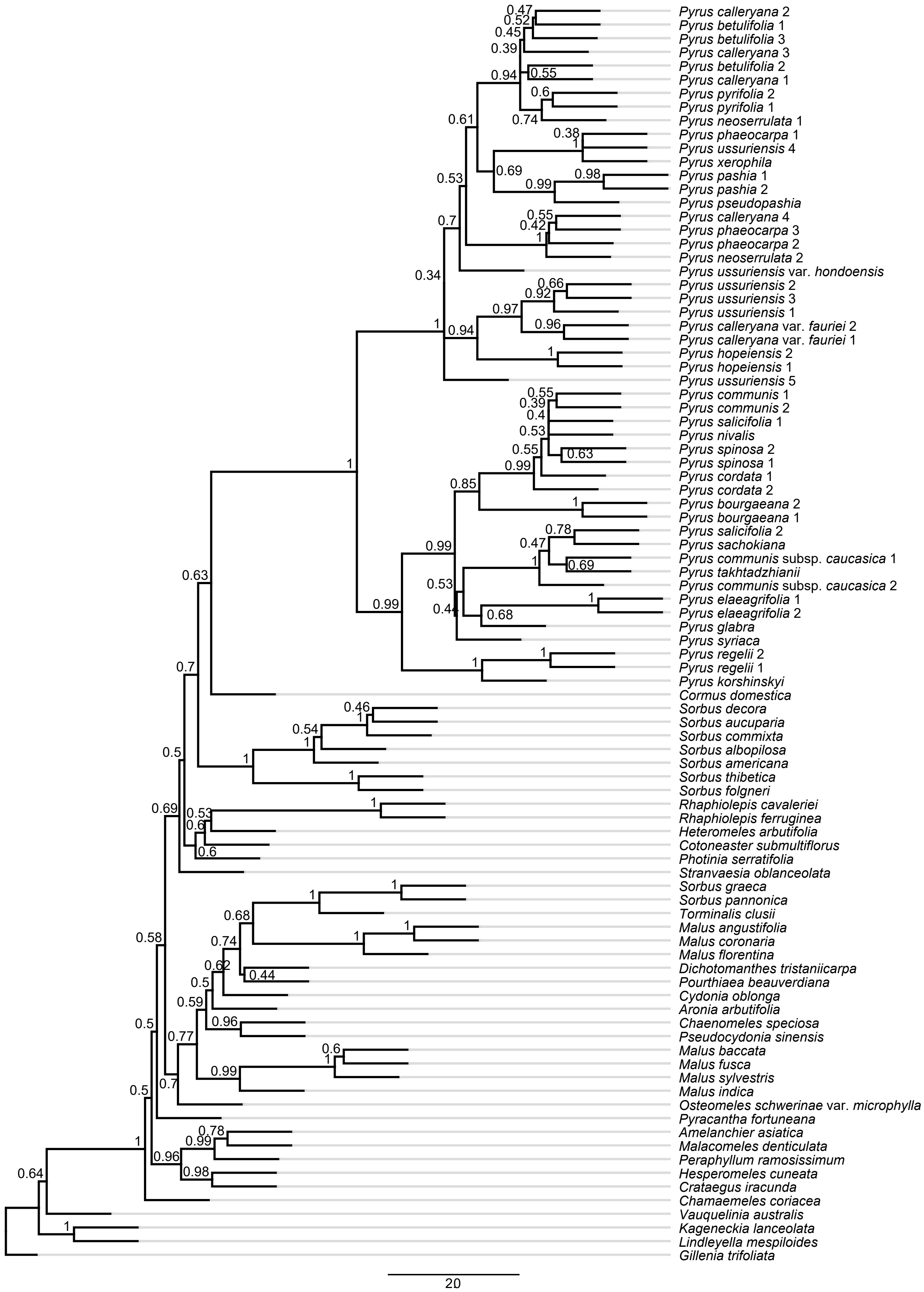


Supplementary Fig. S14. ASTRAL-III Species tree of *Pyrus* inferred from plastid CDS dataset. Local posterior probabilities (LPP) are shown above branches.

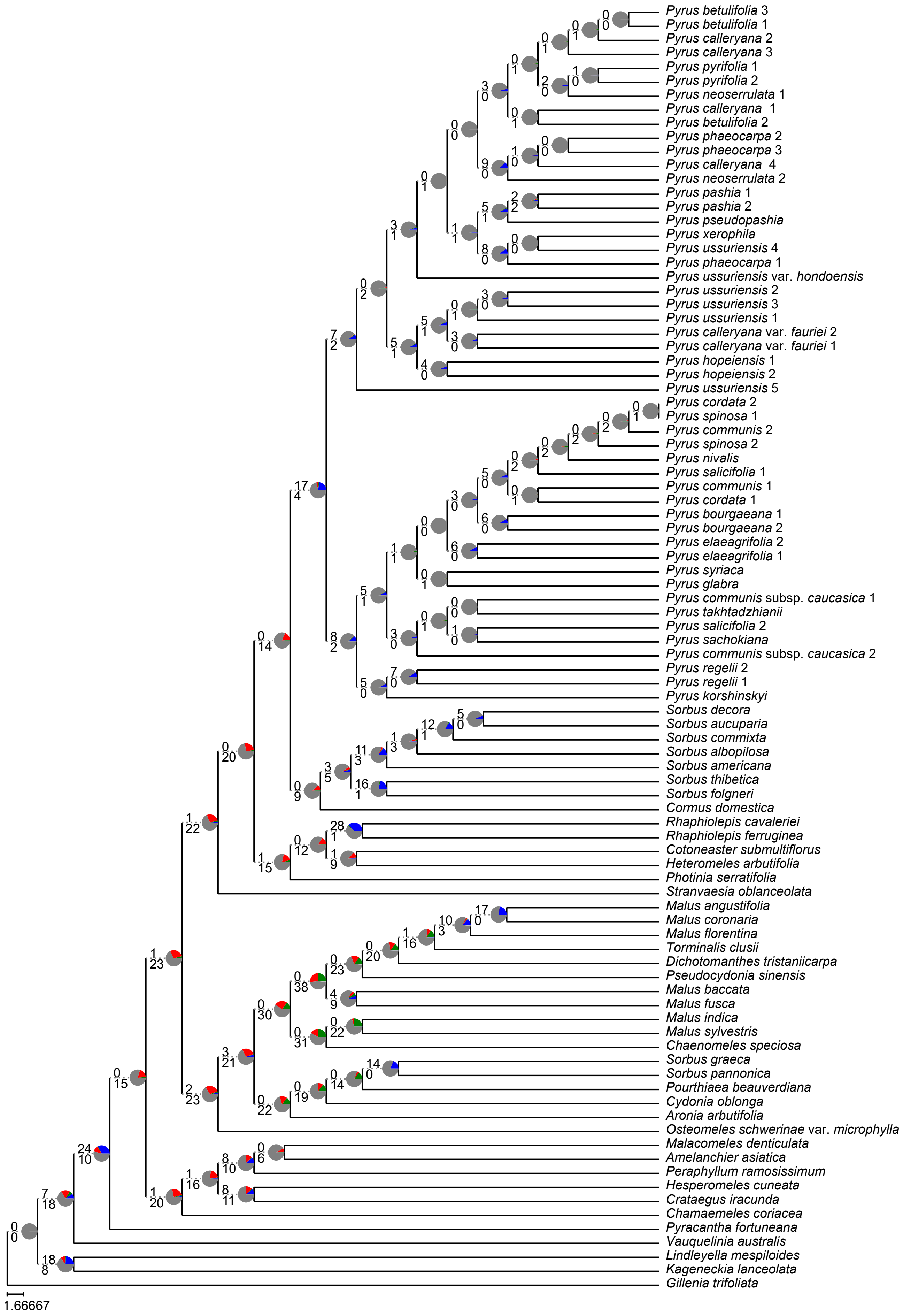


Supplementary Fig. S15. Maximum likelihood phylogeny of *Pyrus* in the framework of Maleae inferred from RAxML analysis of plastid CDS dataset. Pie charts on nodes denote the proportion of gene trees that support that clade (blue), the proportion that support the main alternative bifurcation (green), the proportion that support the remaining alternatives (red), the proportion (conflict or support) that have < 50% bootstrap support (dark grey), and the proportion that have missing taxa (light grey). The number of gene trees concordant/conflicting with that node in the nuclear phylogeny are shown above branches. The Internode Certainty All (ICA) score are shown below branches.

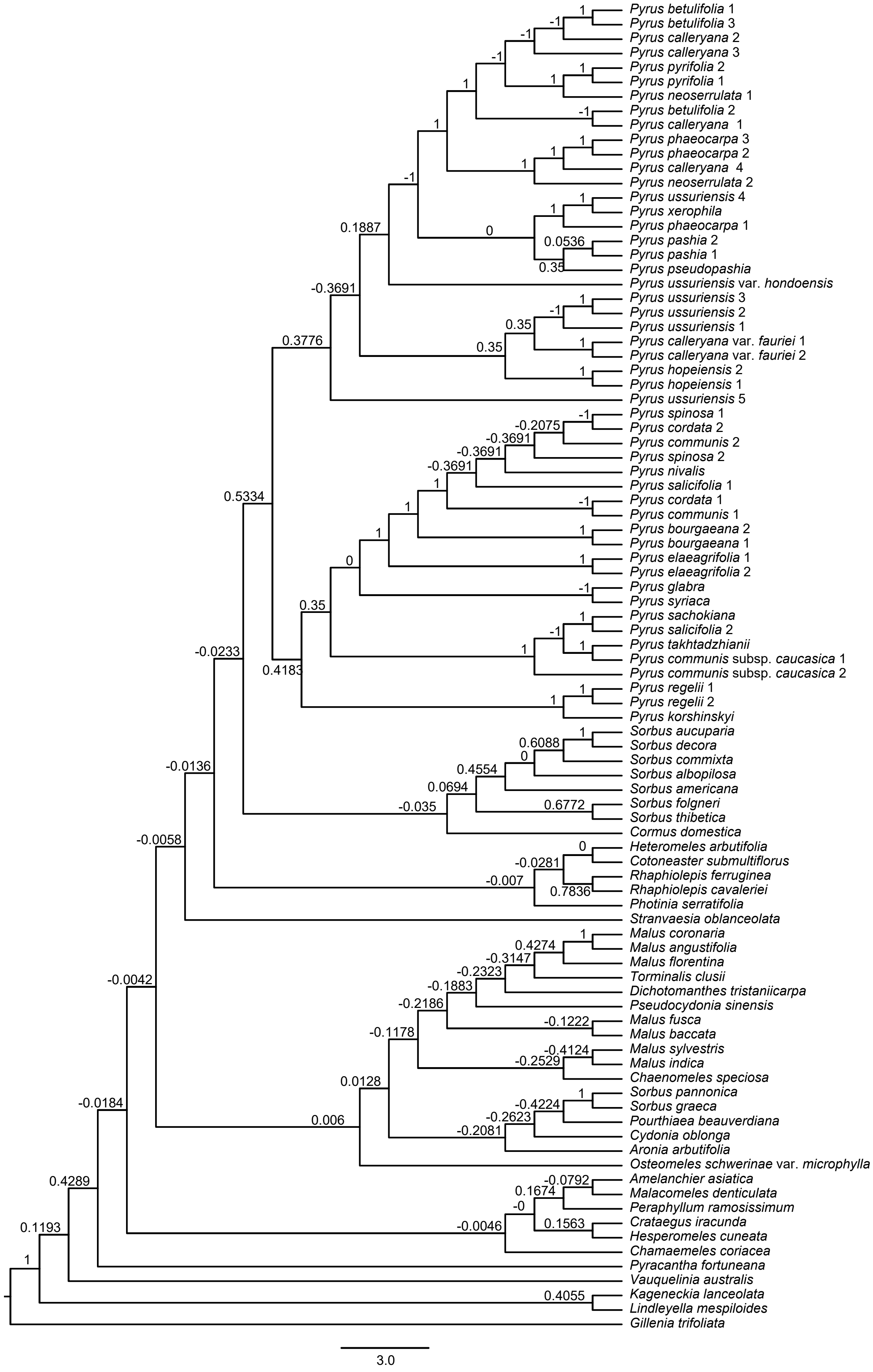


Supplementary Fig. S16. Maximum likelihood phylogeny of *Pyrus* in the framework of Maleae inferred from RAxML analysis of plastid CDS dataset. The Internode Certainty All (ICA) score are shown above branches.

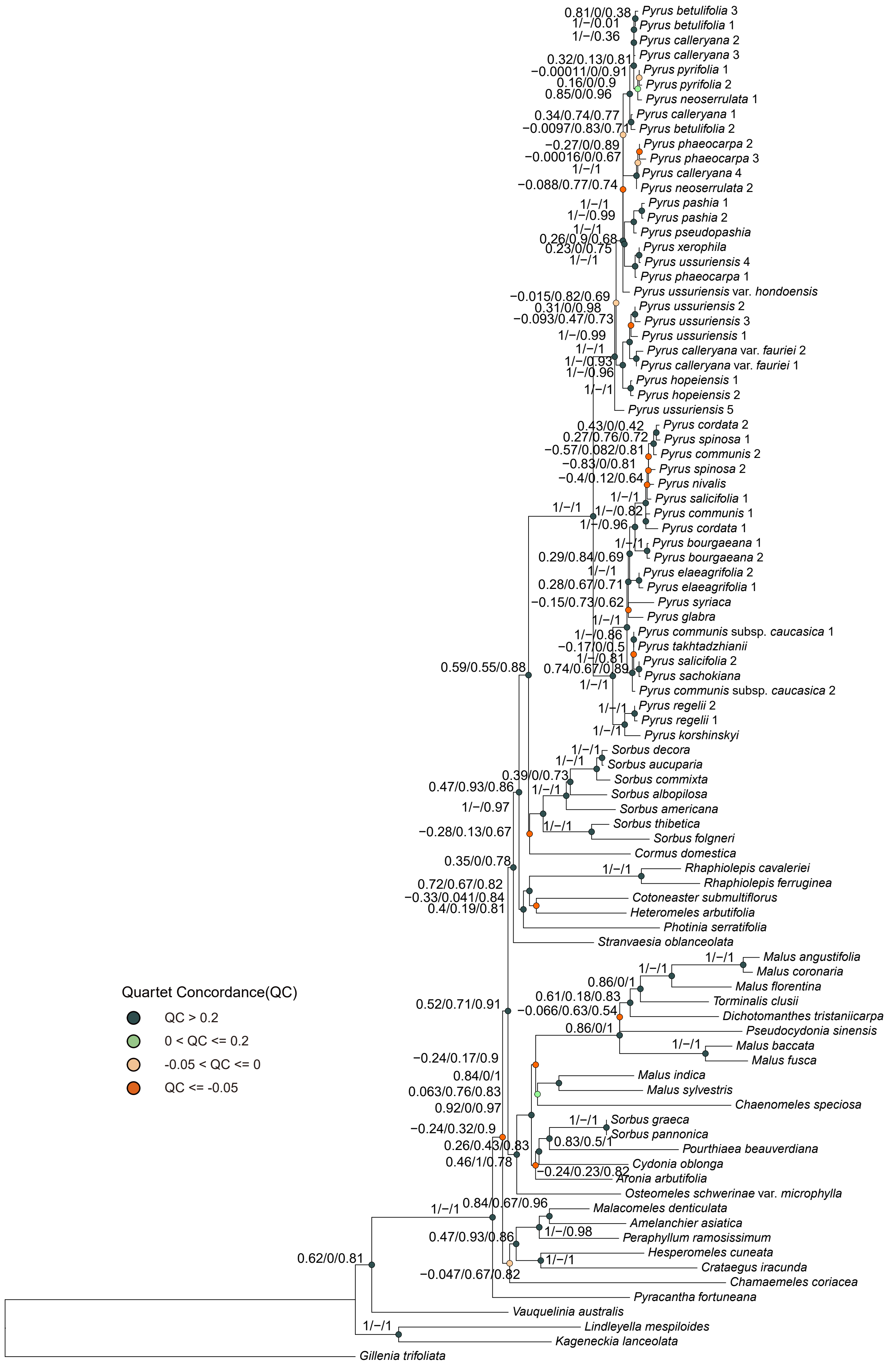


Supplementary Fig. S17. Maximum likelihood phylogeny of *Pyrus* in the framework of Maleae inferred from RAxML analysis of plastid CDS dataset. Quartet Sampling (QS) scores for each node are shown next to branches indicating Quartet Concordance (QC) / Quartet Differential (QD) / Quartet Informativeness (QI). Quartet Concordance is also showed in each node' pie chart and color-coded according to the legend.

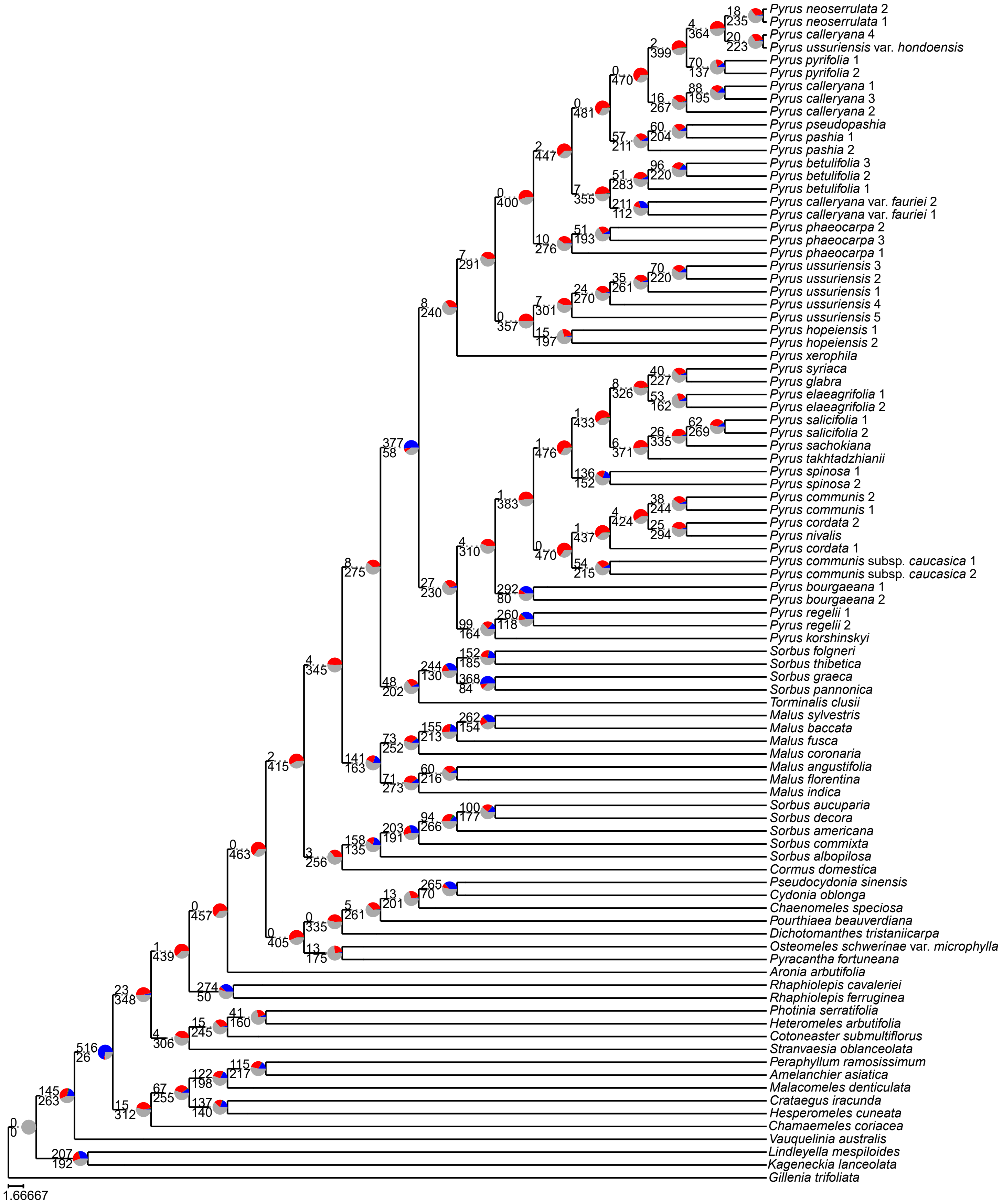


Supplementary Fig. S18. Maximum likelihood phylogeny of *Pyrus* in the framework of Maleae inferred from ASTRAL-III analysis of 771 MO orthologs. Pie charts on nodes denote the proportion of gene trees that support that clade (blue), the proportion that support the main alternative bifurcation (green), the proportion that support the remaining alternatives (red), the proportion (conflict or support) that have < 50% bootstrap support (dark grey), and the proportion that have missing taxa (light grey). The number of gene trees concordant with that node in the nuclear phylogeny are shown above branches. The number of gene trees conflicting with that node in the nuclear phylogeny are shown below branches.

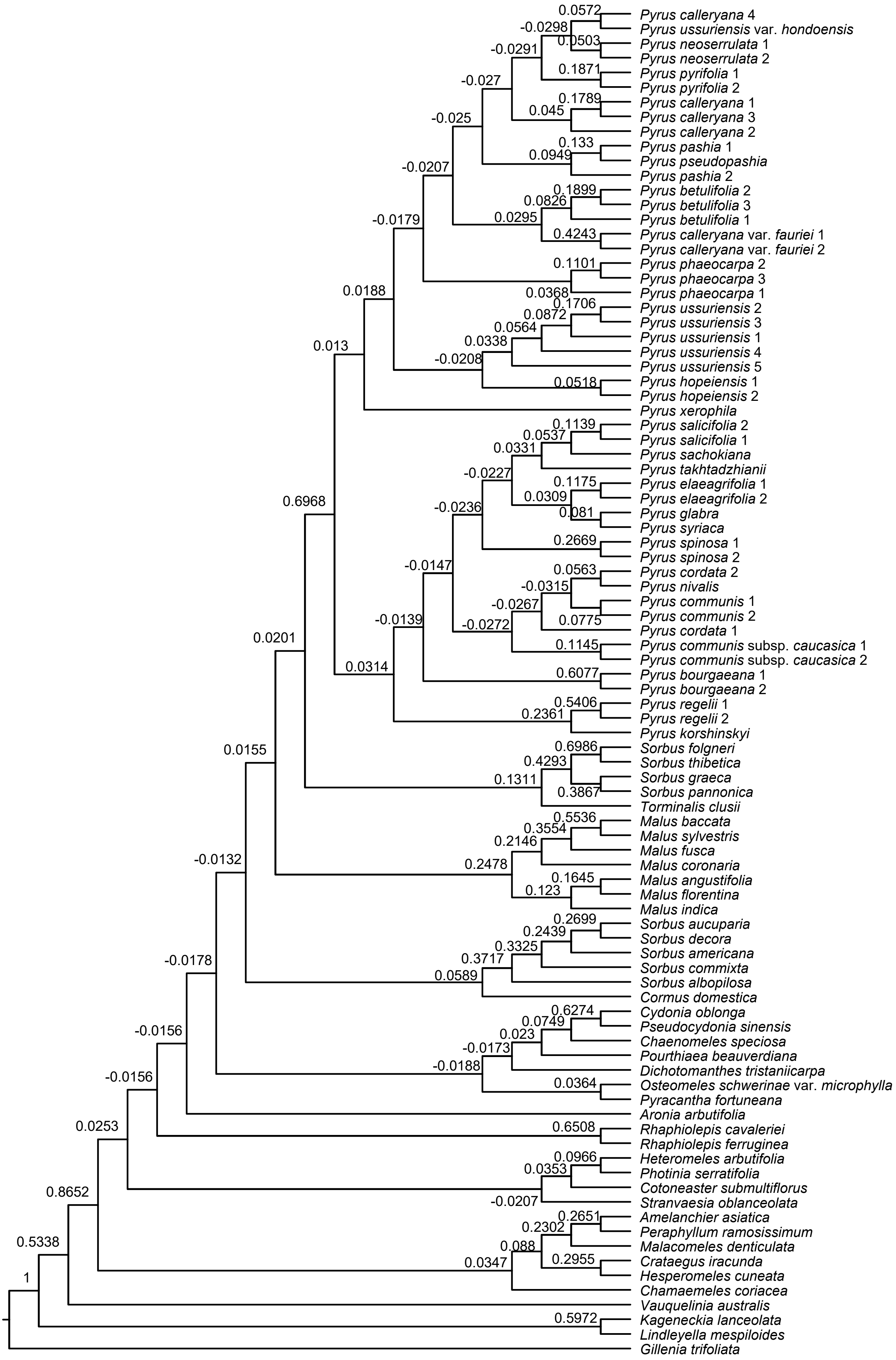


Supplementary Fig. S19. ASTRAL-III Species tree of *Pyrus* in the framework of Maleae inferred from 771 MO orthologs. The Internode Certainty All (ICA) score are shown above branches.

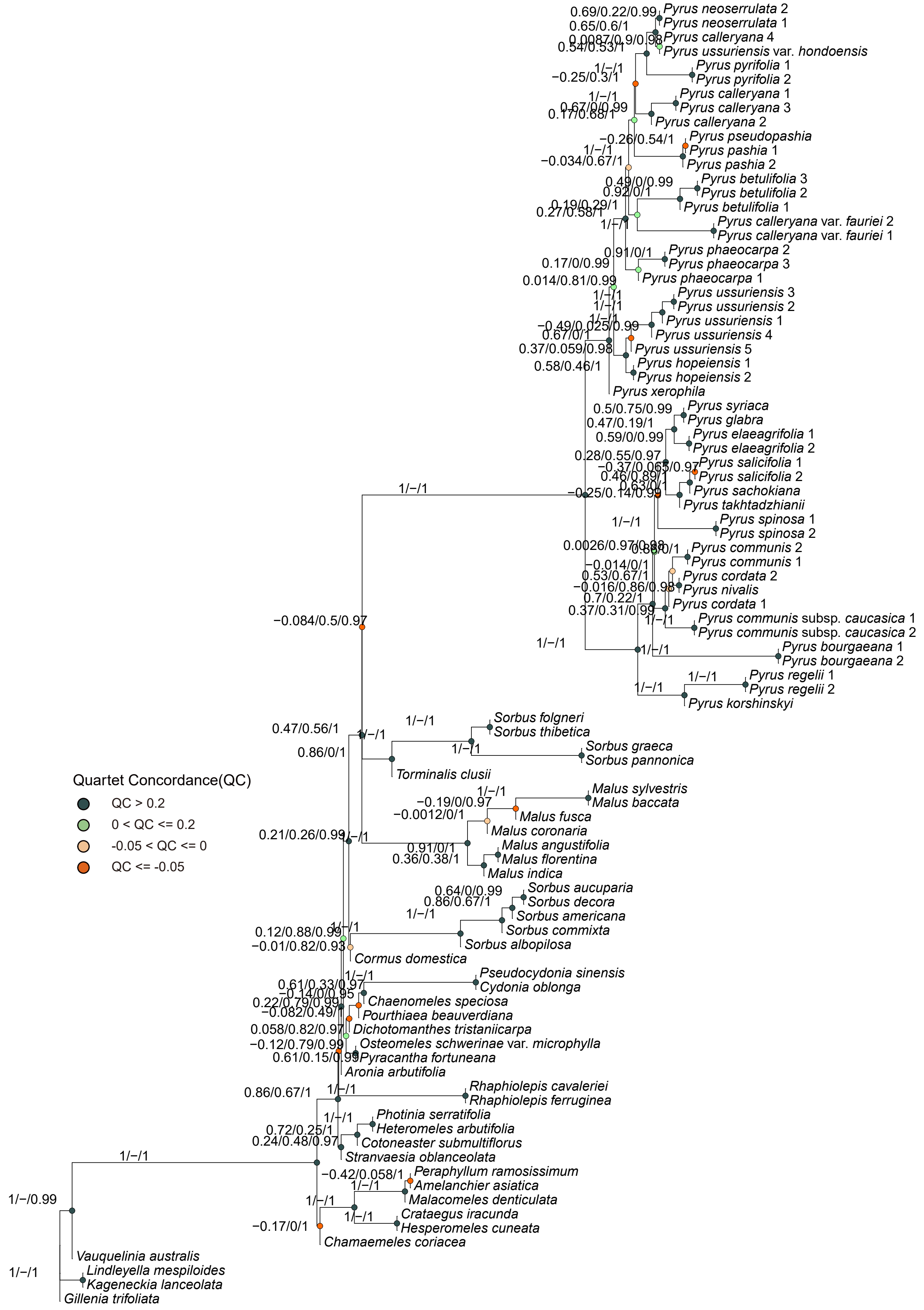


Supplementary Fig. S20. ASTRAL-III Species tree of *Pyrus* in the framework of Maleae inferred from 771 MO orthologs. Quartet Sampling (QS) scores for each node are shown next to branches indicating Quartet Concordance (QC) / Quartet Differential (QD) / Quartet Informativeness (QI). Quartet Concordance is also showed in each node'pie chart and color-coded according to the legend.

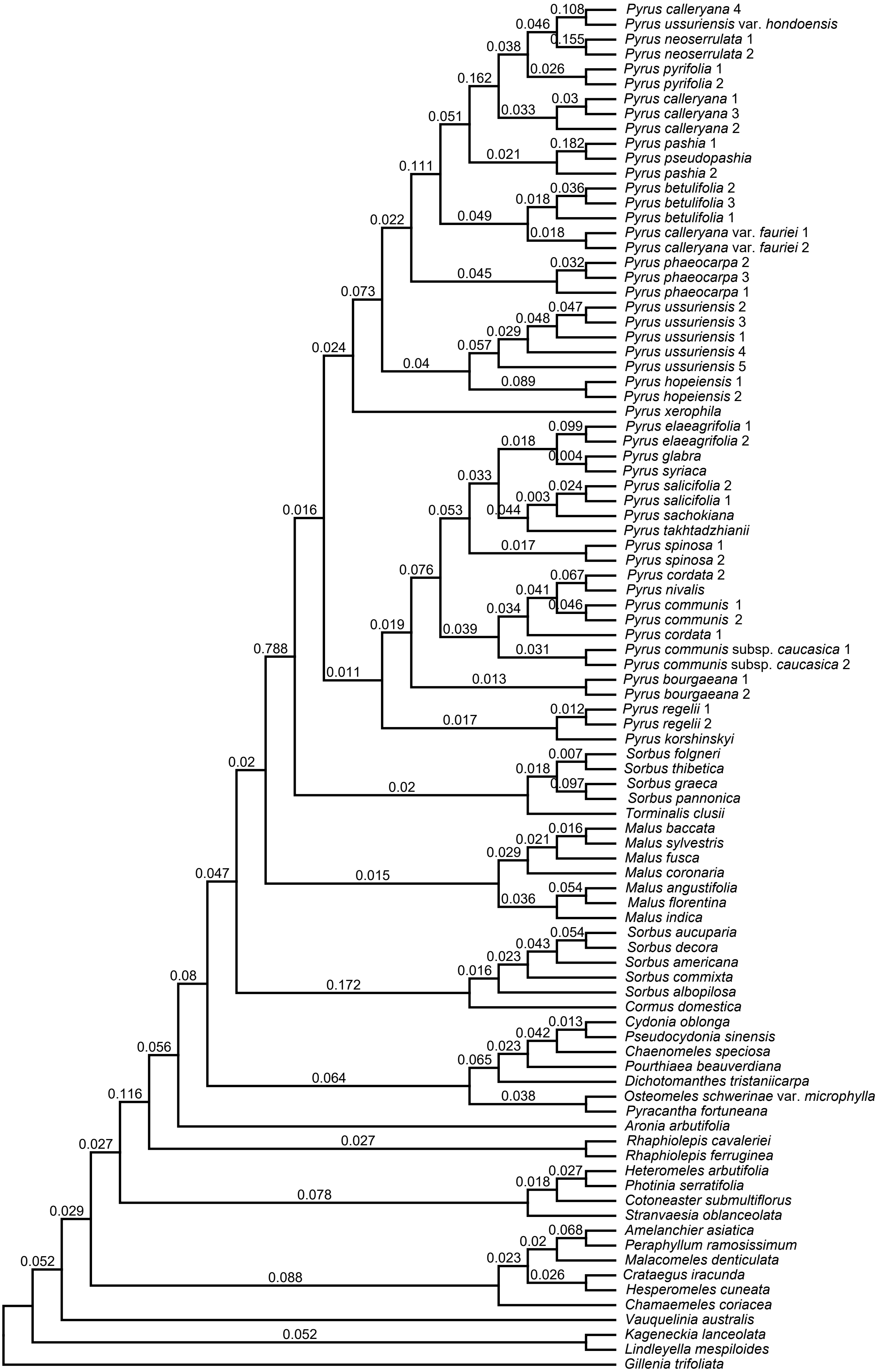


Supplementary Fig. S21. ASTRAL-III Species tree of *Pyrus* in the framework of Maleae inferred from 771 MO orthologs. Population mutatuin parameter theta are shown above branches.

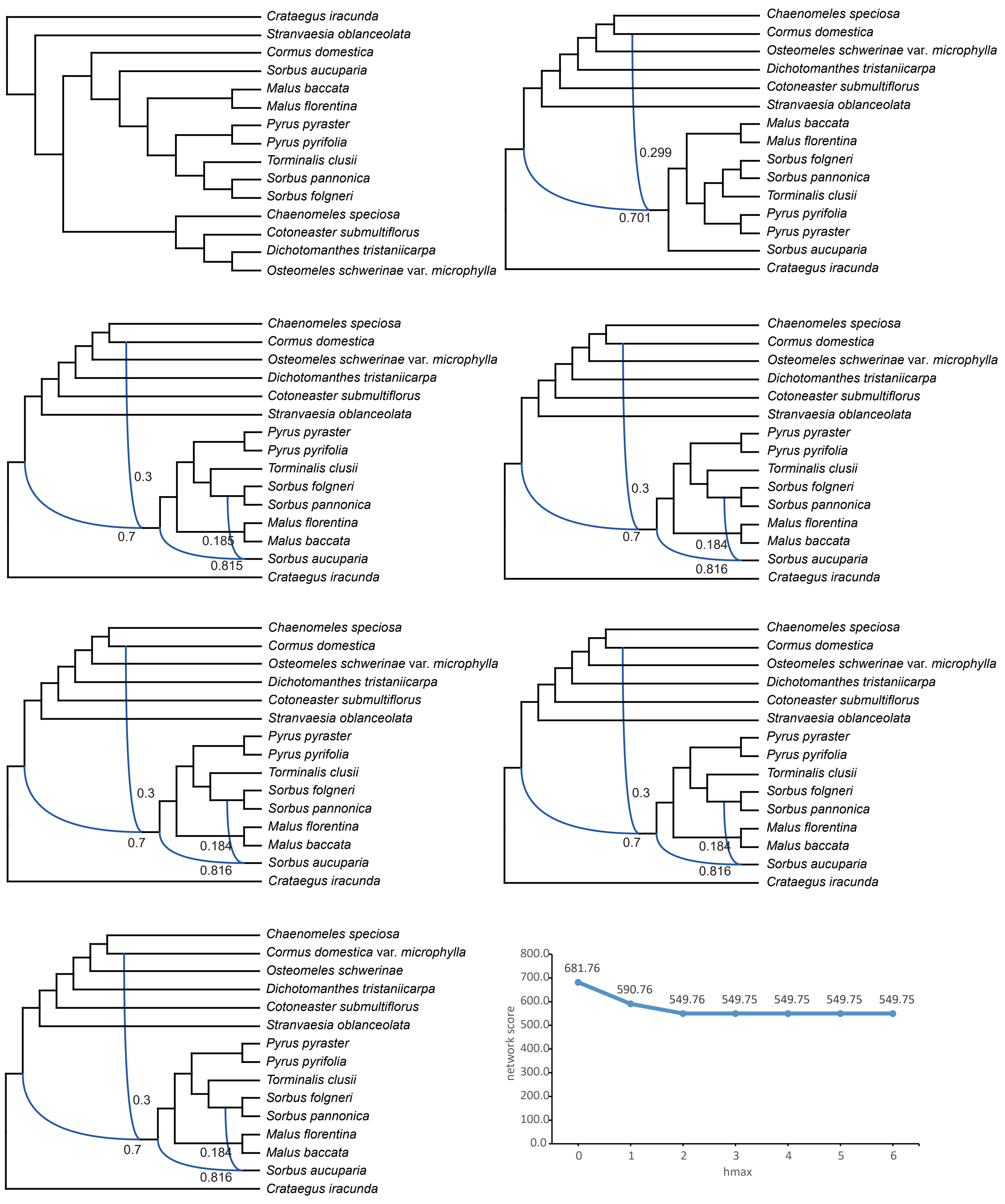


Supplementary Fig. S22. Phylogenetic network analysis from the 15 - taxa sampling of Maleae. Species networks inferred from SNaQ network. Blue curved branches indicate the possible hybridization event. Dark blue and light blue numbers indicate the major and minor inheritance probabilities of hybrid nodes.

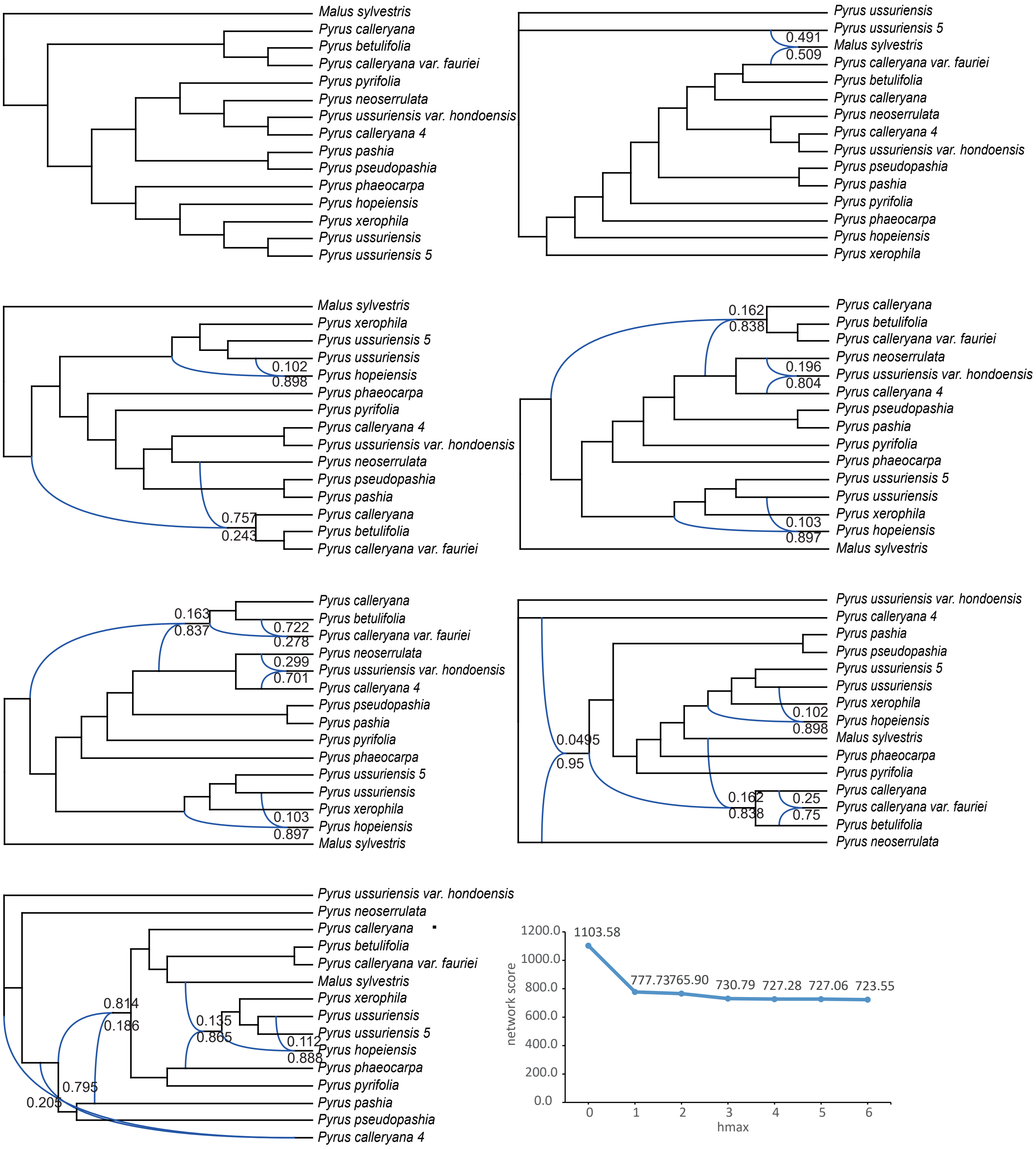


Supplementary Fig. S23. Phylogenetic network analysis from the 14 - taxa sampling of *Pyrus* representing the clade I and its closely relative genera. Species networks inferred from SNaQ network. Blue curved branches indicate the possible hybridization event. Dark blue and light blue numbers indicate the major and minor inheritance probabilities of hybrid nodes.

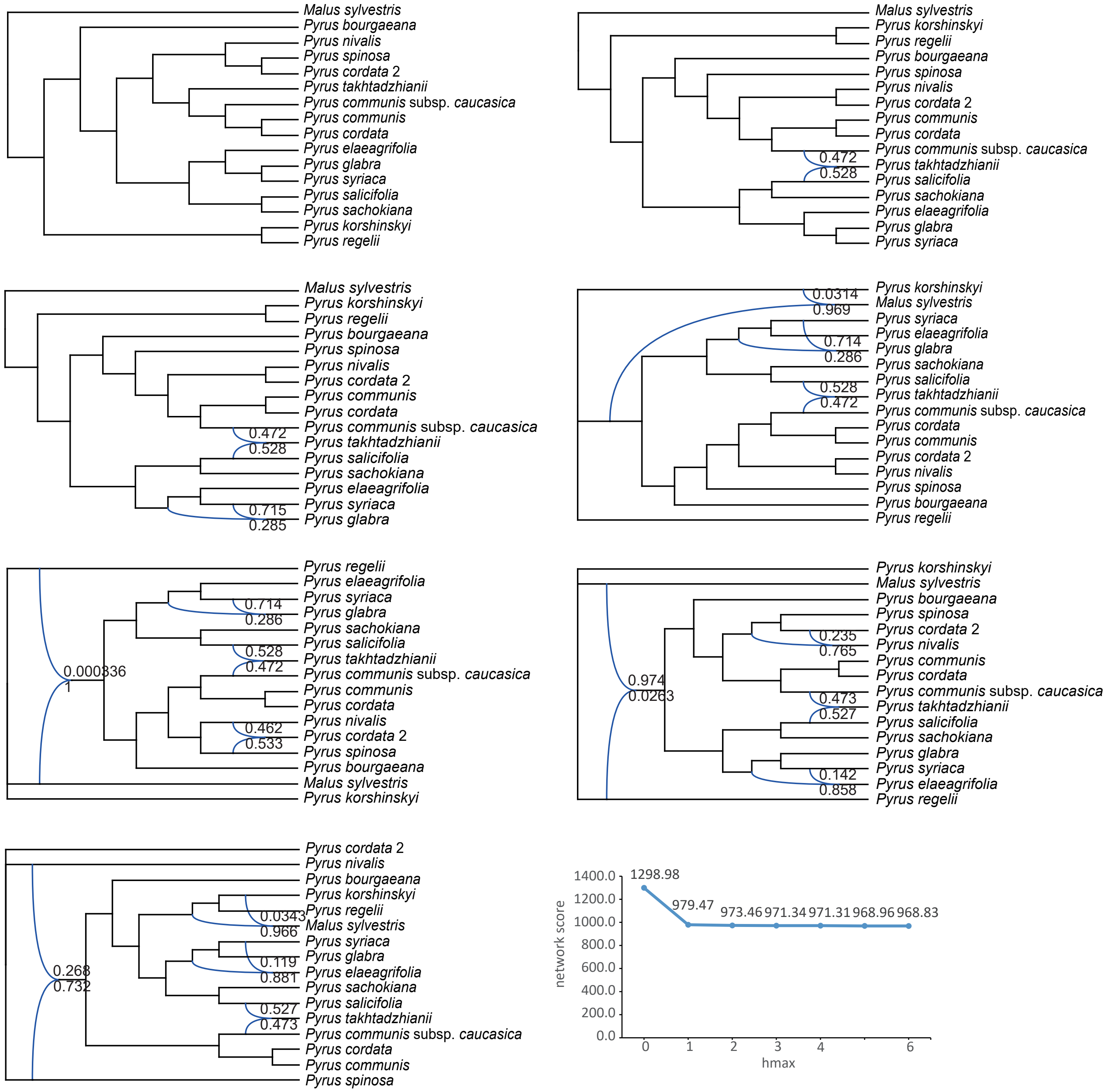


Supplementary Fig. S24. Phylogenetic network analysis from the 15 - taxa sampling of *Pyrus* representing the clade II and its closely relative genera. Blue curved branches indicate the possible hybridization event. Dark blue and light blue numbers indicate the major and minor inheritance probabilities of hybrid nodes.

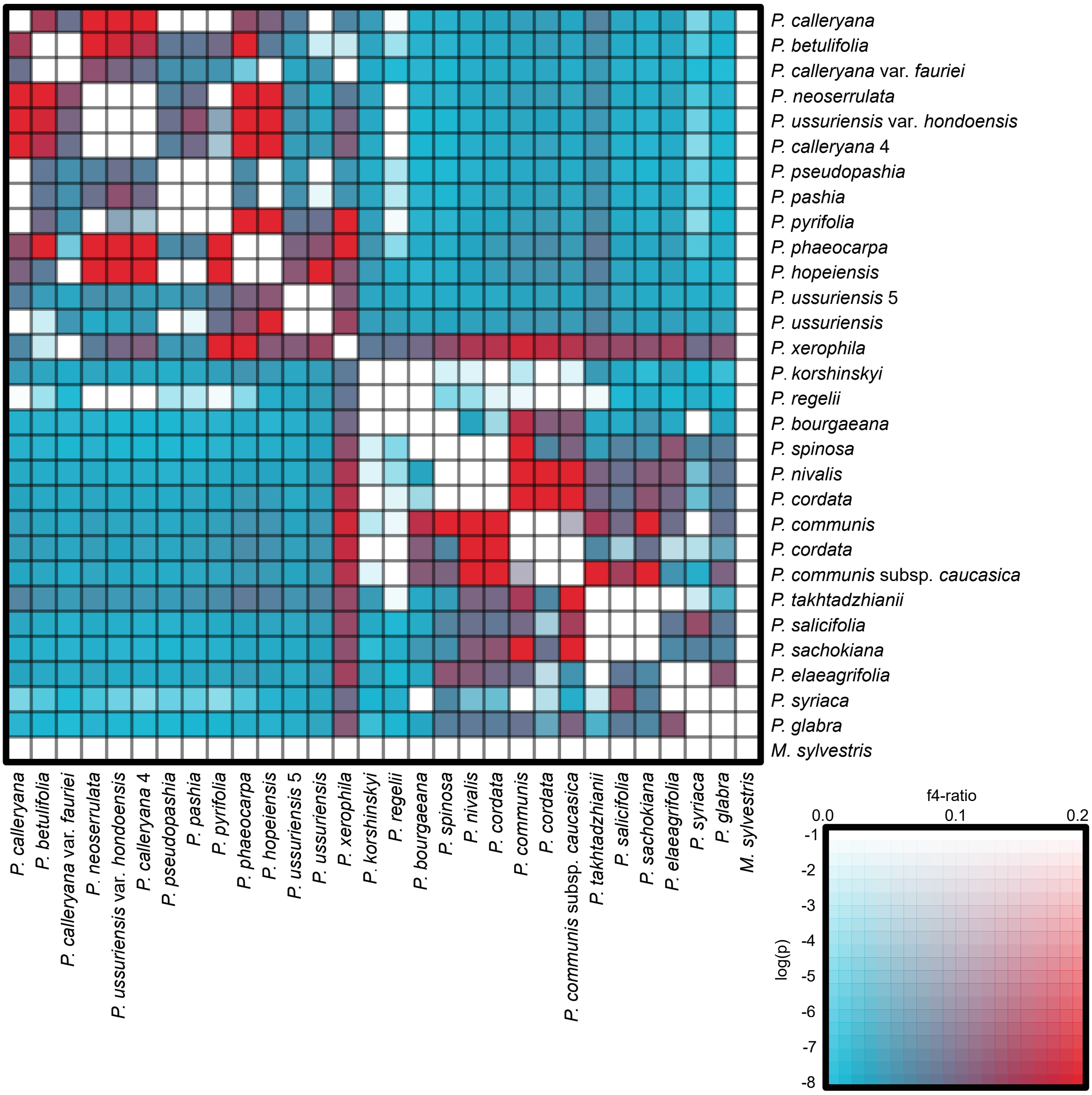


Supplementary Fig. S25. Heat map showing statical support for gene flow between pairs of species inferred from Dsuite package. The shaded scale in boxes represents the estimated f4-ratio branch value.

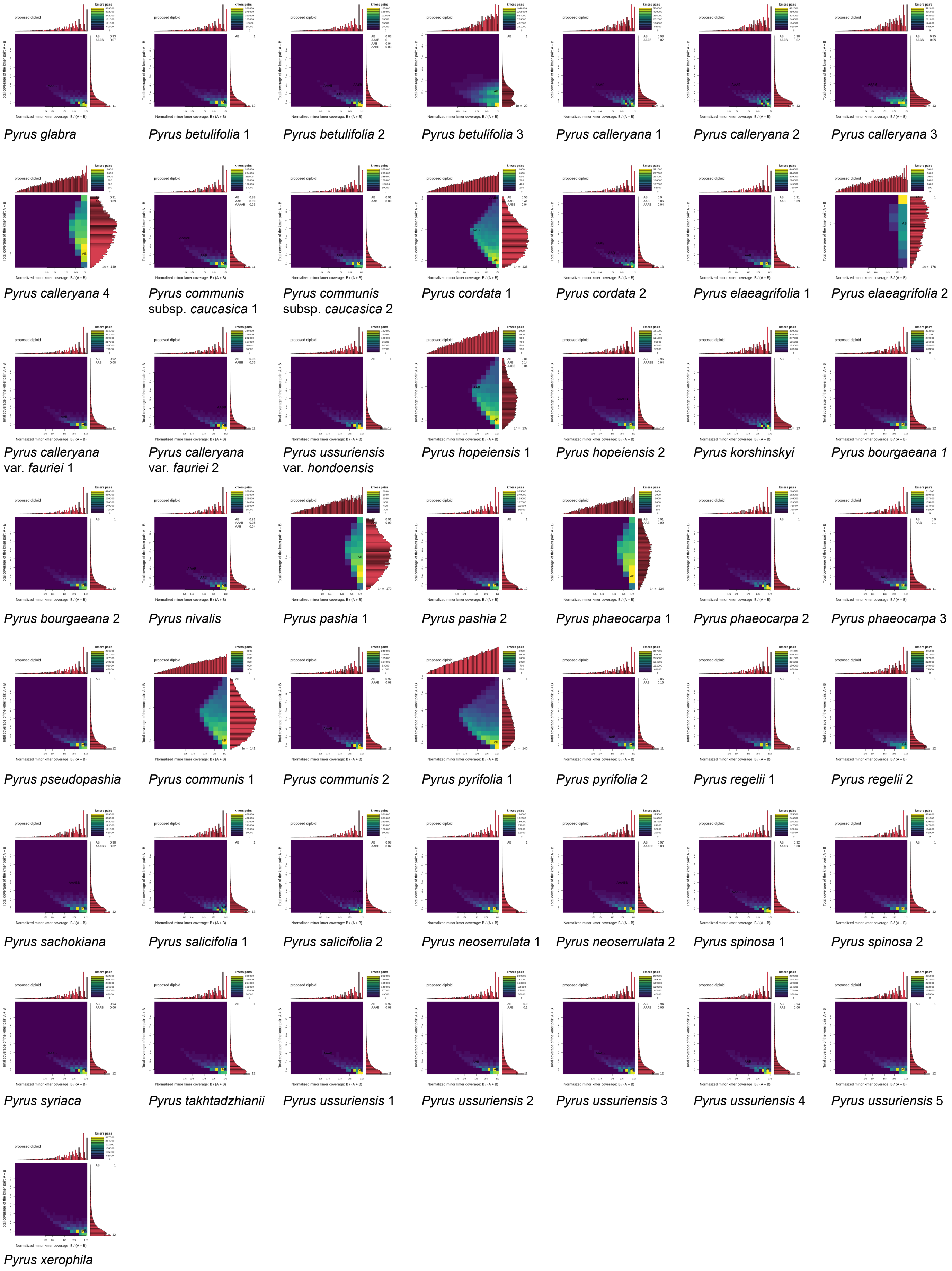


Supplementary Fig. S26. Evaluation of K-mer spectra of *Pyrus* in smudgeplot.

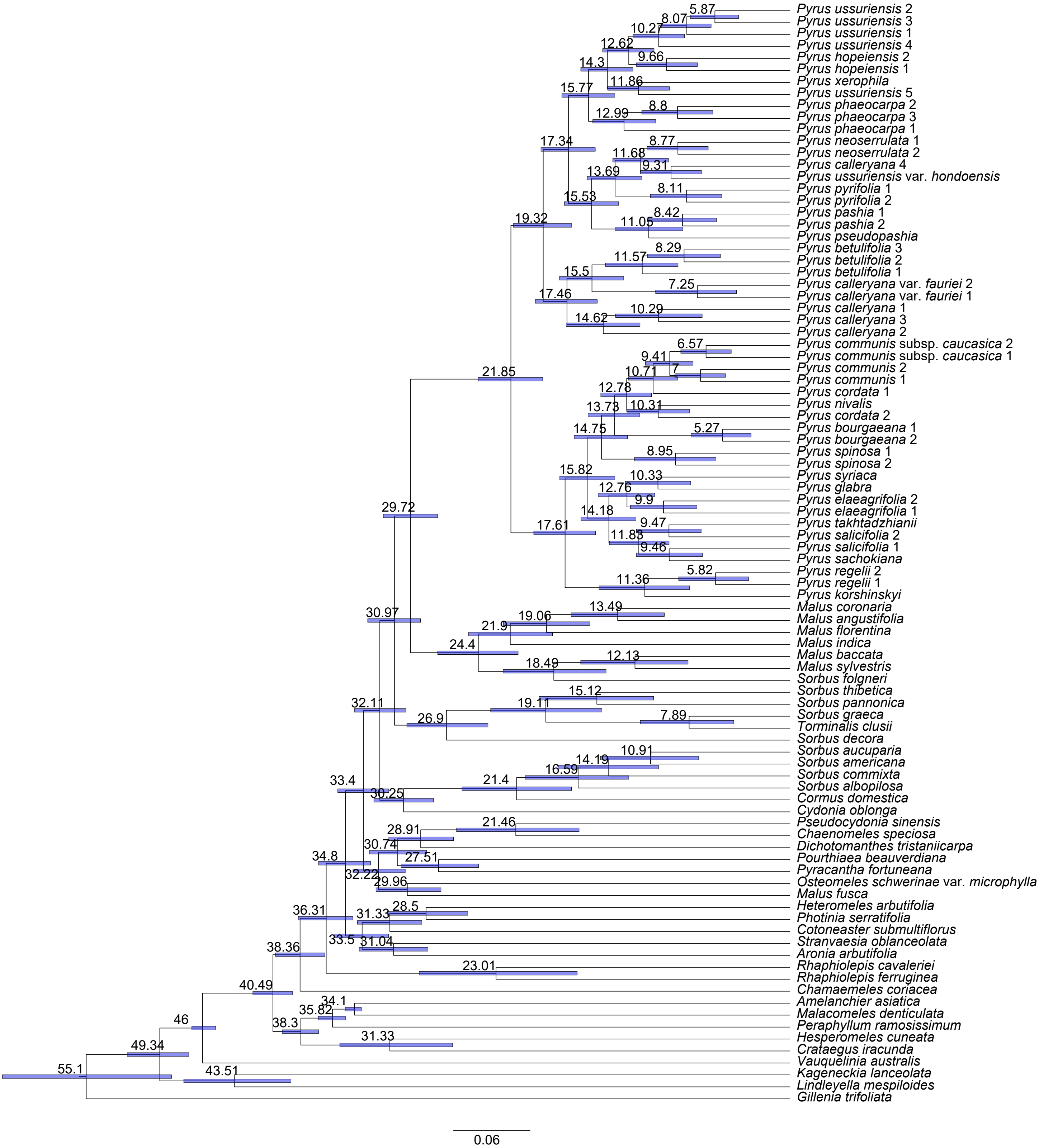


Supplementary Fig. S27. Dated chronogram of Maleae inferred from PAML based on the RAxML concatenated tree inferred from MO orthologs. Maximum clade credibility (MCC) tree showing mean ages above branches. Light blue bars on nodes represent 95% confidence intervals of divergence time estimates.

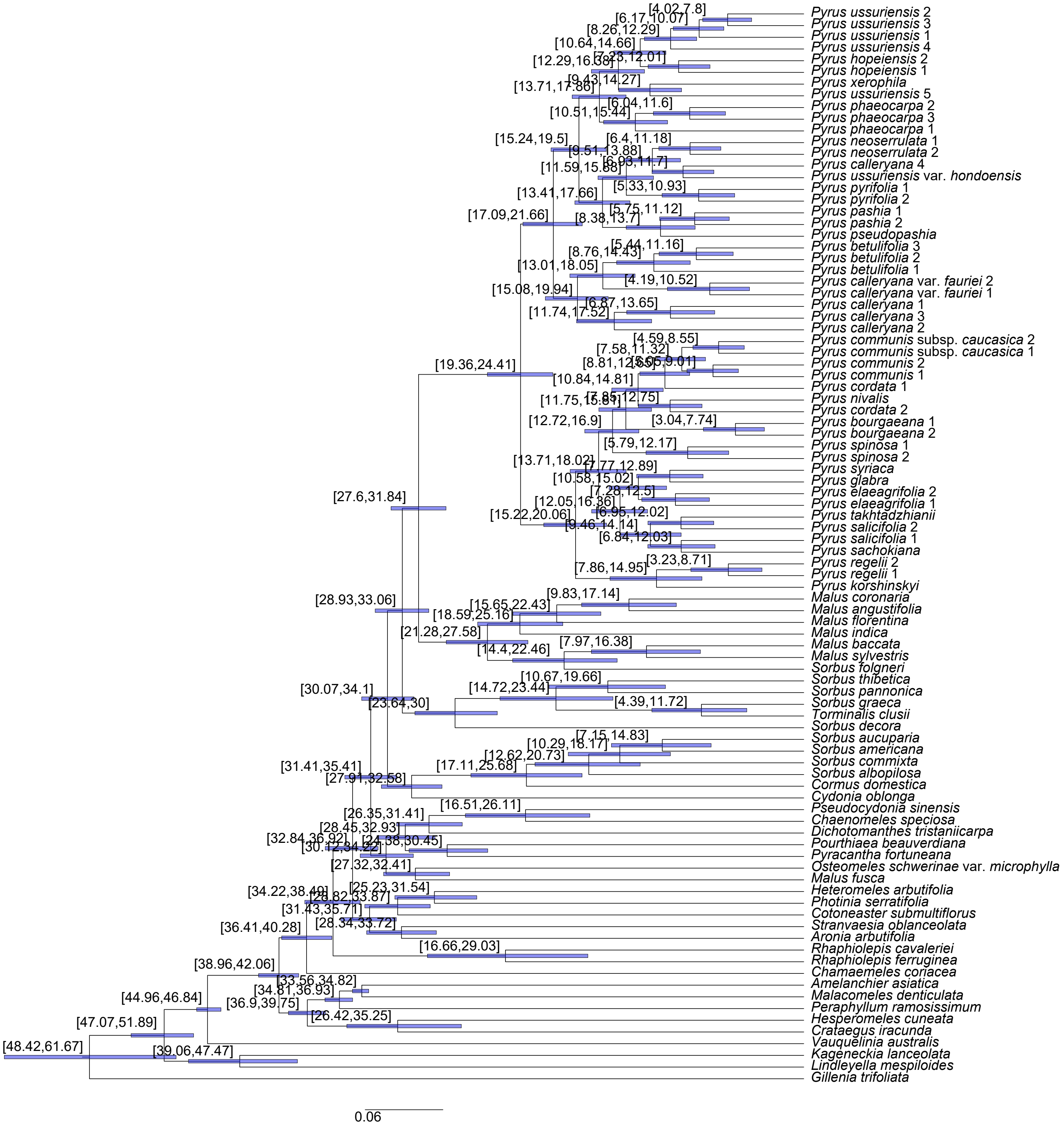


Supplementary Fig. S28. Dated chronogram of Maleae inferred from PAML based on the RAxML concatenated tree inferred from MO orthologs. Node numbers and blue bars indicate 95% confidence intervals of divergence time estimates.

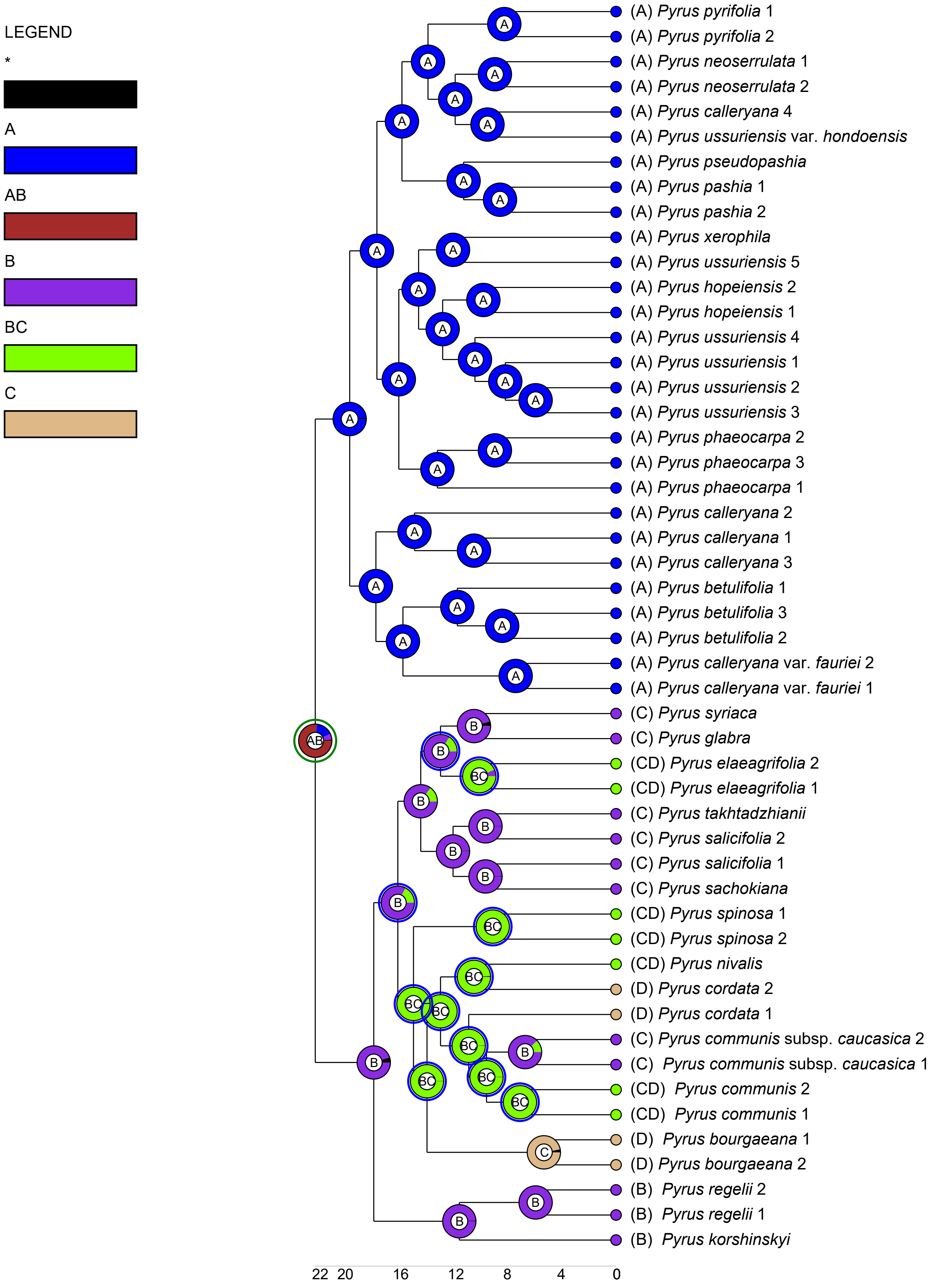


Supplementary Fig. S29. The ancestral area reconstruction using BioGeoBEARS implemented in RASP using the dated chronogram of Maleae inferred from PAML based on the RAxML concatenated tree inferred from MO orthologs, with the colored key identifying extant and possible ancestral ranges. (A) East Asia, (B) Central and West Asia, (C) Europe & North Africa.

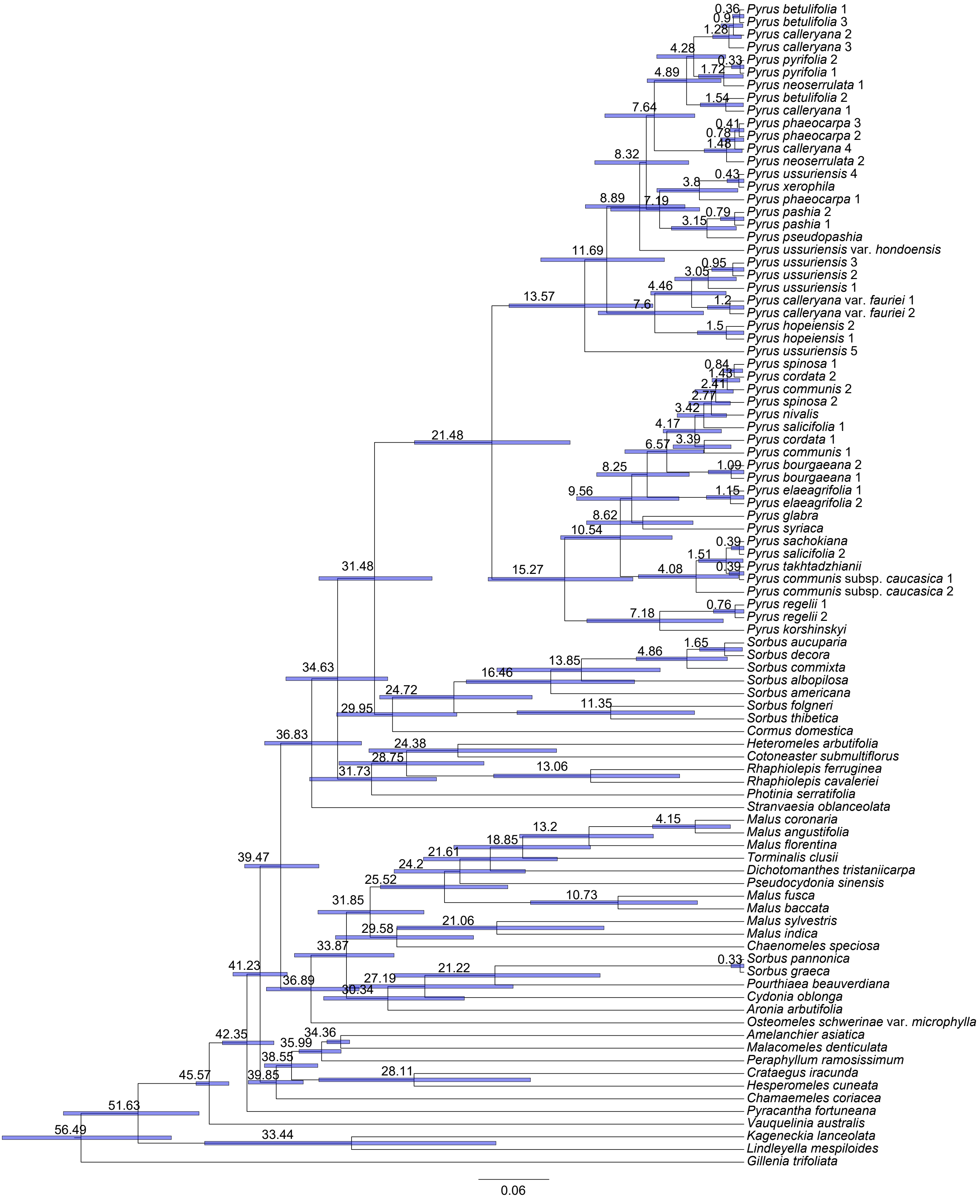


Supplementary Fig. S30. Dated chronogram of Maleae inferred from PAML based on the RAxML concatenated tree inferred from plastid CDS dataset. Maximum clade credibility (MCC) tree showing mean ages above branches. Light blue bars on nodes represent 95% confidence intervals of divergence time estimates.

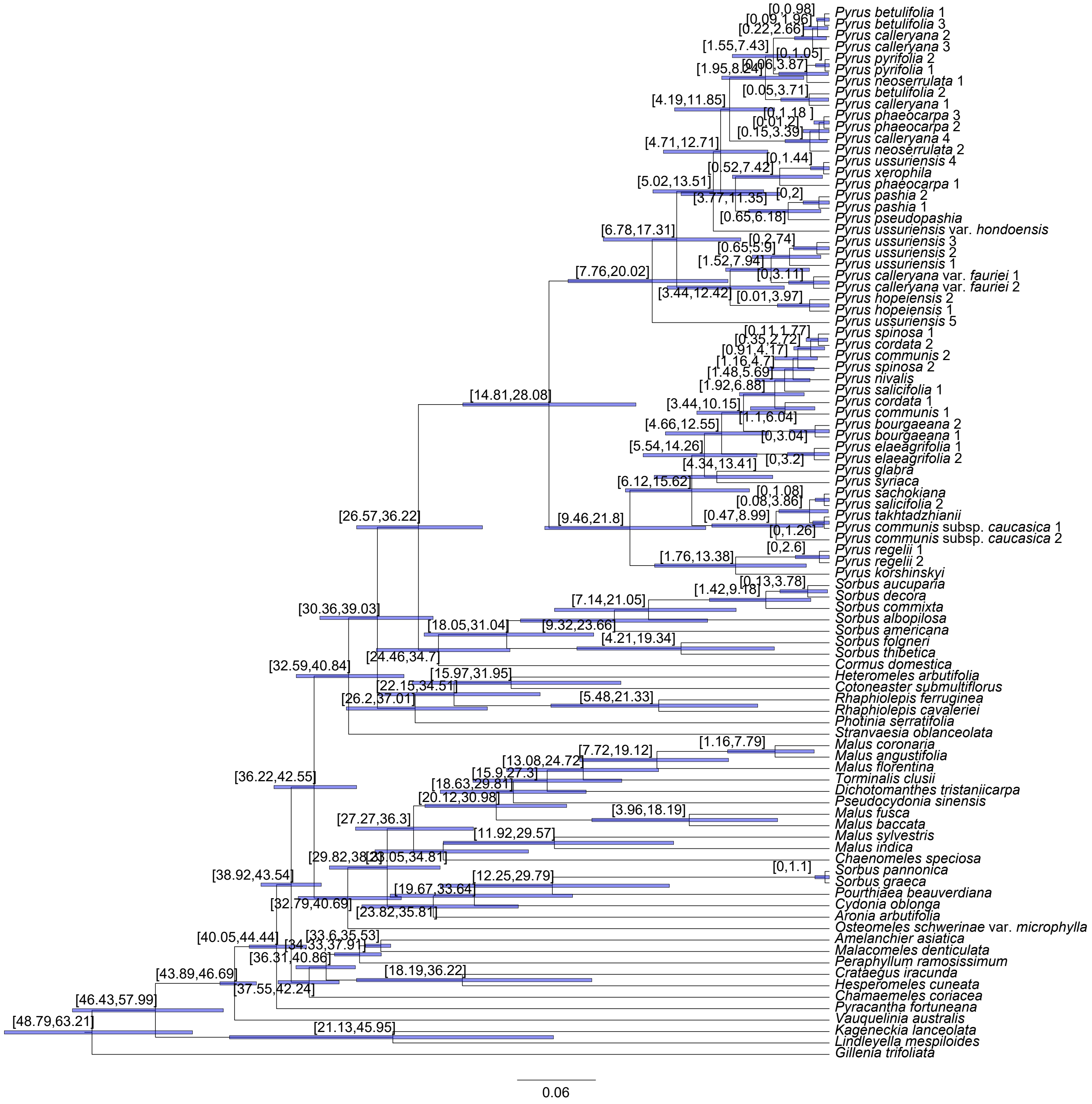


Supplementary Fig. S31. Dated chronogram of Maleae inferred from PAML based on the RAxML concatenated tree inferred from plastid CDS dataset. Node numbers and blue bars indicate 95% confidence intervals of divergence time estimates.

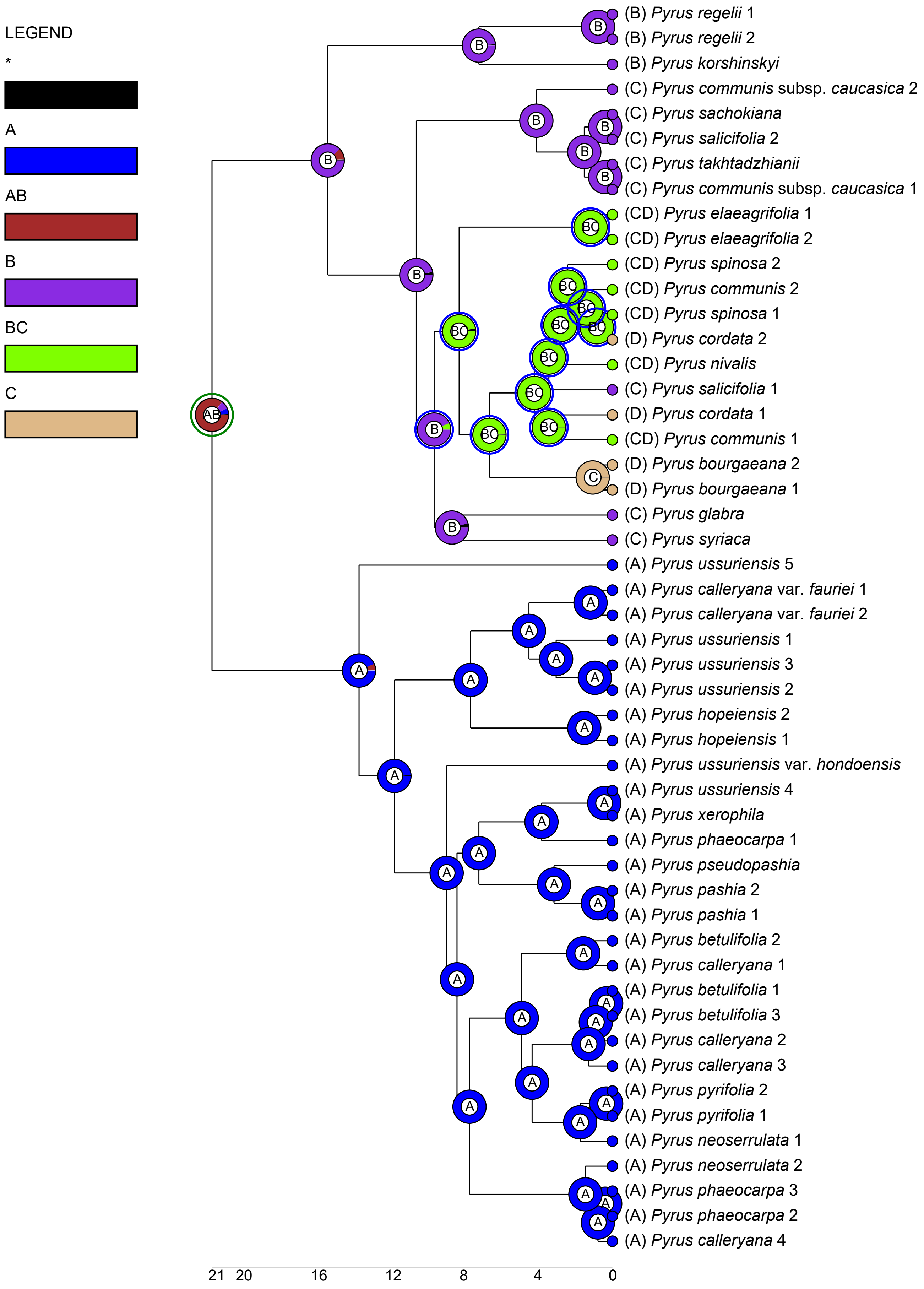


Supplementary Fig. S32. The ancestral area reconstruction using BioGeoBEARS implemented in RASP using the dated chronogram of Maleae inferred from PAML based on the RAxML concatenated tree inferred from plastid CDS dataset, with the colored key identifying extant and possible ancestral ranges. (A) Eastern Asia, (B) Central and Western Asia, (C) Africa, and (D) Euro and North Africa.
