## Supplementary material for "Unraveling the Web of Life: Incomplete lineage sorting and hybridization as primary mechanisms over polyploidization in the evolutionary dynamics of pear species": sup_information.docx

### Supplementary Methods

#### Orthology inference for the nuclear genes

In our study, we implemented the tree-based orthology inference methodology for SCN genes initially described by Yang & Smith (2014) and subsequently refined by Morales-Briones et al. (2022). Briefly, the nuclear genes recovered by Hybpiper were aligned using MAFFT v. 7.480 (Nakamura et al. 2018) with default parameters. Alignments exhibiting gaps in over 90% of sequences were subsequently trimmed using *phyx* (Brown et al. 2017). Preliminary homolog trees were estimated using RAxML v. 8.2.12 (Stamatakis 2014), utilizing a GTRCAT model and 100 bootstrap replicates. Within each homolog tree, tips exhibiting either monophyly or paraphyly and corresponding to the same taxa were systematically excised. Subsequently, the alignments that remained post-trimming, characterized by the maximal count of unambiguous characters, were retained for subsequent analytical procedures. The final homolog trees were refined by removing tips with extremely long branches using TreeShrink v. 1.3.9 (Mai and Mirarab 2018). The cleaned trees then served as the basis for ortholog inference. Two alternative orthology inference methods were applied: the Monophyletic Outgroup (MO) and Rooted Ingroup (RT) approaches (Yang and Smith 2014; Morales-Briones et al. 2022). For the MO method, homolog trees with monophyletic, non-repeating outgroups were first selected, and then orthologs with the minimal number of ingroup taxa after removing paralog from root to tip were kept in each rooted homolog tree. Utilizing the RT approach, we pruned homolog trees encompassing at least one outgroup, aiming to extract distinct ingroup clades. Following this, ortholog trees with a minimal number of ingroup taxa within each ingroup clade were selectively preserved, following the protocol described in the MO approach. In the above MO and RT approaches, homolog trees without repeating taxa and outgroups were also preserved, and the minimum number of ingroup taxa was set to 25 to maximize the number of obtained orthologs. The detailed step-by-step procedures can be referenced at https://bitbucket.org/dfmoralesb/target_enrichment_orthology.

The ortholog processing followed the pipeline described by Liu et al. (2022) to mitigate the potential impact of uneven sequencing coverage on certain orthologs, which may influence phylogenetic inference. Briefly, each locus was aligned using MAFFT v. 7.480 (Nakamura et al. 2018), and trimAL v. 1.2 (Capella-Gutiérrez et al. 2009) was used to removal of poorly aligned regions in columns. Subsequently, the refined genes were concatenated using AMAS v. 1.0 (Borowiec 2016). These concatenated sequences were then fed to Spruceup (Borowiec 2019) for detecting and clipping outlier sequences based on a lognorm criterion in rows. The Spruceup output, applying a distance cutoff of 0.95, was split back to individual loci using AMAS v. 1.0 (Borowiec 2016) and underwent an additional round of trimming with trimAL v. 1.2 (Capella-Gutiérrez et al. 2009). Aligned sequences greater than 250 bp were used to infer maximum likelihood (ML) trees in RAxML v. 8.2.12 (Stamatakis 2014) with the GTRGAMMA model and 100 rapid bootstrap replicates. Tips with disproportionately long branches, identified by TreeShrink v. 1.3.9 (Mai and Mirarab 2018) in each gene tree, were removed from the gene tree and its corresponding alignment. The subsequent phylogenetic analysis was based on these cleaned trees and sequences.

#### Dating analysis

Due to the lack of reliable fossils for calibrating the divergence time of the *Pyrus* genus, we adopted a step-by-step inference approach to enhance the precision of our temporal divergence estimates. For the dating analysis using PAML v. 4.9j (Yang 2007), we designated the ML tree inferred from the nuclear MO dataset as the user-specified tree. The stem age of *Amelanchier* Medik. was calibrated using two fossils, specifically *A. peritula* and *A. scudderi*, which date back to the Late Eocene, spanning 33.9 to 37.2 Mya. In addition, the leaf fossil *Vauquelinia comptoniifolia* from the Eocene discovered in the United States provided the basis for constraining the stem age of the genus *Vauquelinia* to an estimated range of 40.4-46.2 Mya (MacGinitie 1969). As a secondary calibration point, the crown clade of the Maleae tribe was temporally set to 60 Mya, following the findings of Zhang et al. (2017). Given the substantial computational requirements inherent in divergence time estimation for large datasets, we applied the approximate likelihood method within the MCMCTree algorithm. The priors for molecular rates were set according to an independent-rates model (specified as ‘clock = 2’), while the site substitution models adhered to the GTR models (denoted as ‘model = 7’). Two separate Markov Chain Monte Carlo (MCMC) runs, each initialized with different seed values, were executed. The initial one million iterations of each run were discarded as burn-in. Sampling was then conducted every 10 iterations until a total of 500,000 samples were collected. The analysis of stationarity and convergence for each run was performed using Tracer v. 1.7.1, ensuring that the effective sample size (ESS) for all parameters exceeded the threshold of 200. In order to facilitate a comparative analysis with the findings derived from the nuclear MO dataset, we also estimated the ultrametric tree for the plastid CDS dataset.

For the BEAST2 (Bouckaert et al. 2014) analysis, we generated a 30-taxa dataset to mitigate computational intensity. This dataset was curated to include 29 high-quality samples representing 26 currently recognized *Pyrus* species, complemented by an additional sample from the *Malus* genus, serving as the outgroup. The GTR+G model was employed for site substitution, the fossilized birth-death model for the tree prior, and the uncorrelated relaxed lognormal model for the tree prior. The MCMC chains were extended over 100 million generations, with a sampling frequency set at every 1,000 generations. The convergence and adequacy of the parameter estimations were rigorously evaluated using Tracer v. 1.7, ensuring that the ESS for all parameters surpassed the threshold of 200. This assessment was conducted after combining the log files from six independent runs of BEAST2 using LogCombiner v. 1.10. Subsequently, the final Maximum Credibility Clade (MCC) tree was summarized using TreeAnnotator v. 1.10, based on the combined tree files generated by LogCombiner v. 1.10.
